# Synergistic induction of blood-brain barrier properties

**DOI:** 10.1101/2023.02.09.527899

**Authors:** Gergő Porkoláb, Mária Mészáros, Anikó Szecskó, Judit P. Vigh, Fruzsina R. Walter, Ricardo Figueiredo, Ildikó Kálomista, Zsófia Hoyk, Gaszton Vizsnyiczai, Ilona Gróf, Jeng-Shiung Jan, Fabien Gosselet, Melinda K. Pirity, Monika Vastag, Natalie Hudson, Matthew Campbell, Szilvia Veszelka, Mária A. Deli

## Abstract

Blood-brain barrier (BBB) models derived from human stem cells are powerful tools to improve our understanding of cerebrovascular diseases and to facilitate drug development for the human brain. Yet providing stem cell-derived endothelial cells with the right signaling cues to acquire BBB characteristics while also retaining their vascular identity remains challenging. Here, we show that the simultaneous activation of cyclic AMP and Wnt/β-catenin signaling, and inhibition of the TGF-β pathway in endothelial cells robustly induce BBB properties *in vitro*. To target this novel interaction, we present a small molecule cocktail named cARLA, which synergistically enhances barrier tightness in a range of BBB models across species. Mechanistically, we reveal that the three pathways converge on Wnt/β-catenin signaling to mediate the effect of cARLA *via* the tight junction protein claudin-5. We demonstrate that cARLA shifts the gene expressional profile of human stem cell-derived endothelial cells towards the *in vivo* brain endothelial signature, with a higher glycocalyx density and efflux pump activity, lower rates of endocytosis and a characteristic endothelial response to proinflammatory cytokines. Finally, we illustrate how cARLA can improve the predictive value of human BBB models regarding the brain penetration of drugs and targeted nanoparticles. Due to its synergistic effect, high reproducibility and ease of use, cARLA has the potential to advance drug development for the human brain by improving BBB models across laboratories.

**Significance Statement:** The blood-brain barrier (BBB) hinders drug delivery to the brain and is implicated in neurological diseases. To better understand these processes in humans, there is a need for culture models that mimic the complexity of the BBB. However, state-of-the-art human BBB models either suffer from a non-physiological, mixed epithelial-endothelial identity or have weak barrier tightness, which greatly limits their usability. We identified a molecule combination that synergistically enhances barrier tightness in several *in vitro* models and induces complex BBB properties in human stem cell-derived endothelial cells by targeting a novel link between three signaling pathways. The molecule combination has the potential to improve BBB culture models across laboratories to advance both basic research and drug development for the human brain.

## Introduction

Endothelial cells (ECs) lining the smallest blood vessels in the brain possess unique anatomical and functional properties, collectively known as the BBB^1^. Through specialized tight junction complexes, low rates of vesicular transcytosis, a negatively charged glycocalyx as well as efflux and influx transport systems, the BBB precisely controls the composition of the neural microenvironment^2^. While these characteristics protect the brain from harmful substances, the barrier also prevents most drugs from entering the central nervous system, hindering drug development for neurological diseases^3^. Additionally, BBB breakdown is a hallmark of a range of neuropathologies^4,5^, yet these processes are incompletely understood in humans at the molecular level. To provide mechanistic insight on BBB (dys)function and improve the prediction of drug delivery to the brain, there is a need for *in vitro* models that faithfully mimic the human BBB^6^.

Culture models based on primary brain microvascular ECs isolated from animal tissue have been widely used as *in vitro* systems to study the BBB since the 1980s^7^. However, there are major interspecies differences in the BBB proteome related to drug transport^8,9,10^, which negatively impacts the translatability of findings from animal models to clinical trials. As a human alternative with good scalability, several BBB models have been established using induced pluripotent stem cells^11,12,13,14,15,16,17,18,19,20,21^ or CD34^+^ hematopoietic stem cells isolated from umbilical cord blood^22,23,24,25^. Although these protocols represent major technological advances, state-of-the-art human BBB models either have high barrier tightness but suffer from a mixed epithelial-endothelial identity or are definitive vascular ECs with weak barrier properties^26,27^. In either case, the usability of these models is limited to only certain applications^26,27^.

To solve this problem, two-step differentation has emerged as a promising strategy, in which stem cells are first differentiated into vascular ECs and brain-like features are then induced in a second step^15,21,22^. This is critical as BBB properties are not intrinsic to ECs but are promoted and maintained by organ-specific signaling cues from pericytes^28,29,30^, astrocytes^31,32,33^, and the microenvironment *in vivo*. While stem cell-derived vascular ECs are mostly responsive to inductive cues from co-culture with pericytes or astrocytes^15,21,22^, they need additional factors for proper junctional maturation and to acquire a complex BBB phenotype *in vitro*. Motivated by the unmet need for an approach that is efficient, reproducible and easily adaptable by other laboratories, we focused on targeting developmentally relevant signaling pathways at the BBB using soluble factors.

During development, Wnt/β-catenin signaling controls brain angiogenesis^34,35,36^ and BBB formation^37,38,39^, and this pathway has been activated *in vitro* by Wnt ligands (Wnt-3a, Wnt-7a/b) or small molecules (LiCl, 6-BIO, CHIR99021) to induce a subset of barrier properties^22,40,41,42,43^. Recently, TGF-β receptor antagonists (RepSox, A83-01) have also been shown to elevate junctional tightness in human stem cell-derived ECs^44,45^ *via* claudin-5, the dominant tight junction protein at the BBB^46^. Notably, treatment with cyclic AMP-elevating agents (pCPT-cAMP, forskolin and the cAMP-specific phosphodiesterase inhibitor Ro-20-1724) has previously been demonstrated to increase the resistance across bovine brain EC monolayers alone, and especially in combination with astrocyte-conditioned medium^46,48,49^. Since astrocytes and their conditioned media can be a source of Wnt ligands^50,51^ and Wnt signaling-associated genes were reported to be induced by RepSox^44^, we hypothesized that the cAMP, Wnt/β-catenin and TGF-β pathways converge during BBB maturation, which can be leveraged to increase barrier properties in culture.

Here, we show that the simultaneous activation of cAMP and Wnt/β-catenin signaling while inhibiting the TGF-β pathway synergistically increases BBB tightness in ECs. Using our small molecule cocktail containing pCPT-cAMP+Ro-20-1724+LiCl+A83-01, which we termed as cARLA, we demonstrate a simple and efficient approach to induce barrier properties *in vitro* that is reproducible across a range of BBB models from different species. We provide mechanistic insight into how these signaling pathways interact at the molecular level, and illustrate how the complex phenotype induced by cARLA can potentially improve the prediction of drug– and nanoparticle delivery across the human BBB.

## Results

### cARLA synergistically enhances BBB tightness *via* claudin-5

First, we performed an impedance-based screen of small molecules and recombinant proteins for their ability to enhance barrier tightness in cultured human stem cell-derived ECs supplemented with pericyte-conditioned medium (**Fig. 1A**). This approach allowed us to test selected activators of cAMP and Wnt as well as inhibitors of TGF-β signaling combinatorially in a 96-well plate format, in real time (**Fig. 1B**). After optimizing treatment durations and concentrations (**SI Appendix, Fig. S1A-F**), we asked whether combining Wnt activation with TGF-β inhibition increases barrier integrity compared to targeting single pathways. Interestingly, we observed an increase but only by specific combinations, most effectively by LiCl+A83-01 (**SI Appendix, Fig. S2A-H**).

**Figure 1.**
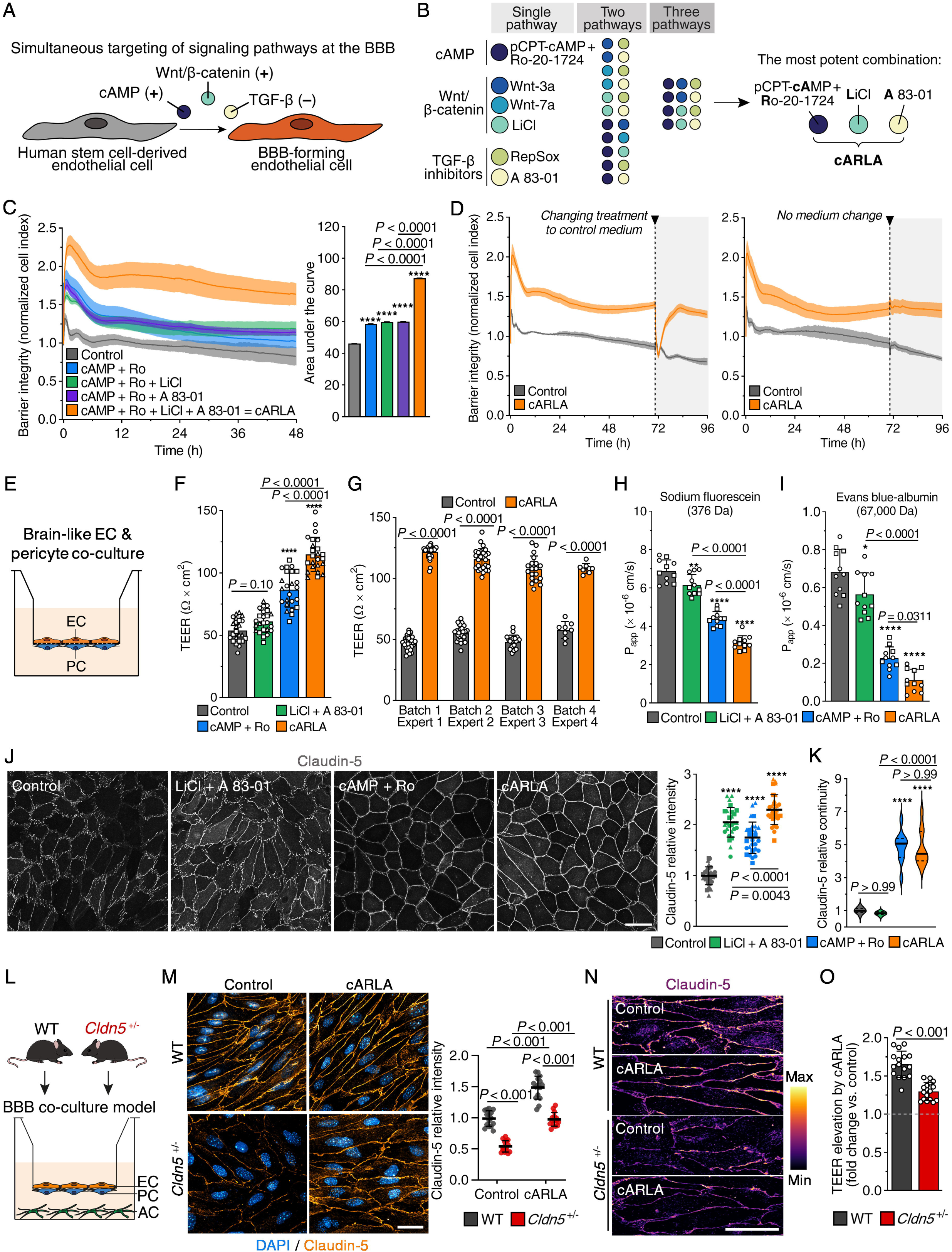
cARLA synergistically enhances barrier tightness *via* claudin-5. **A**) Rationale. **B)** Schematic drawing of single compounds and their combinations tested in the impedance-based screen. **C)** Barrier integrity of human stem cell-derived EC monolayers supplemented with PC-conditioned medium, measured by impedance. Higher normalized cell index values and a higher area under the curve indicates increased barrier integrity. Mean ± SD, ANOVA with Bonferroni’s post-hoc test, \*\*\*\**P*<0.0001 compared to the control group, *n*=6. **D)** Impedance kinetics of EC monolayers with or without changing cARLA treatment to control medium at 72 h. Mean ± SD, *n*=6. **E)** Schematic drawing of the BBB model: human stem cell-derived ECs acquire brain-like characteristics upon co-culture with PCs. **F)** Transendothelial electrical resistance (TEER) in the co-culture model after 48 h treatment. Mean ± SD, ANOVA with Bonferroni’s post– hoc test, \*\*\*\**P*<0.0001 compared to the control group, *n*=24 from 3 independent experiments. **G)** Reproducibility of TEER measurements after 48 h cARLA treatment across experiments, measured by different experts using different batches of cells. Mean ± SD, ANOVA with Bonferroni’s post-hoc test, *n*=82 from 4 independent experiments. **H)** Permeability of sodium fluorescein and **I)** Evans blue-albumin across the co-culture model after 48 h treatment. P_app_: apparent permeability coefficient. Mean ± SD, ANOVA with Bonferroni’s post-hoc test, \**P*<0.05, \*\**P*<0.01, \*\*\*\**P*<0.0001 compared to the control group, *n*=11 from 2 independent experiments. **J)** Claudin-5 immunostaining in human brain-like ECs. Bar: 50 µm. Mean ± SD for intensity and **K)** median ± quartiles for continuity, ANOVA with Bonferroni’s post-hoc test, \*\*\*\**P*<0.0001 compared to the control group, *n*=27-30 from 3 independent experiments. **L)** Schematic drawing of BBB co-culture models isolated from wild type (WT) and *Cldn5^+/-^* mice. **M-N)** Claudin-5 immunostaining in the mouse BBB co-culture model. Bar: 50 µm in both subpanels. Mean ± SD, Two-way ANOVA with Bonferroni’s post-hoc test, *n*=15. **O)** Measurement of TEER in the mouse BBB co-culture models. Values are presented as fold change (cARLA vs. control). Mean ± SD, unpaired *t*-test, *t*=6,903, *df*=30, *n*=16.

We then asked if Wnt activation by LiCl or TGF-β inhibition by A83-01 can potentiate the barrier integrity-promoting effect of cAMP signaling in ECs. Targeting the cAMP pathway alone by pCPT-cAMP supplemented with the cAMP-specific phosphodiesterase inhibitor Ro-20-1724 (cAMP+Ro) resulted in an immediate and potent increase in barrier integrity that diminished over time (**Fig. 1C; SI Appendix, Fig. S3A,B**). The addition of either LiCl or A83-01 did not enhance the effect of cAMP+Ro (**Fig. 1C**). However, the combination of cAMP+Ro+LiCl+A83-01, which we termed as cARLA, synergistically elevated barrier integrity in ECs (**Fig. 1C**), *i.e*. the effect of cARLA was larger than the sum of its parts. This effect could be reproduced by other combinations that simultaneously activate cAMP and Wnt as well as inhibit TGF-β signaling (**SI Appendix, Fig. S3C-E**), but we have found cARLA to be the most potent combination in our screen (**Fig. 1B; SI Appendix, Fig. S3E**). Added to this, the effect of cARLA on barrier integrity was long-lasting (>72 h) and resilient to withdrawal of treatment from ECs by replacing cARLA with control medium (**Fig. 1D; SI Appendix Fig. S3E, S4A**).

We validated these findings in a BBB model in which human stem cell-derived ECs acquire brain-like characteristics (from now on referred to as brain-like ECs) upon co-culture with pericytes (**Fig. 1E**)^22^. Transendothelial electrical resistance (TEER) was synergistically increased upon cARLA treatment by 2.1-fold compared to the control group (**Fig. 1F; SI Appendix, Fig. S4B**), an effect that was highly reproducible across experiments (**Fig. 1G**). Moreover, we measured a lower permeability of tracers sodium fluorescein (2.2-fold) and Evans blue-albumin (7.4-fold compared to the control group) across the co-culture model upon cARLA treatment (**Fig. 1H-I**), indicating that cARLA tightened both the para– and transcellular BBB. As a proposed mediator of its barrier tightening effect, cARLA elevated both the staining intensity (2.3-fold) and continuity (4.5-fold) of the tight junction protein claudin-5 at cell borders (**Fig. 1J-K**). Importantly, similar results were obtained in EC monoculture (**SI Appendix, Fig. S5A-F**), indicating that the effect of cARLA does not depend on other soluble factors from pericytes. We also confirmed the effect of cARLA on barrier tightness, and its specificity for claudin-5 in an induced pluripotent stem cell-derived human BBB model with definitive vascular characteristics (EECM-BMEC-like cells^21^, **SI Appendix, Fig. S6A-K**).

To test if the effect of cARLA is dependent on claudin-5, we assembled BBB models consisting of primary brain ECs, pericytes and astrocytes isolated from wild type (WT) mice or claudin-5 heterozygous (*Cldn5^+/-^*) mice that have only one copy of *Cldn5* (**Fig. 1L**). Compared to WT cultures, *Cldn5^+/-^* cultures had 49% less claudin-5 at the protein level, which could be rescued by cARLA treatment (**Fig. 1M**). Similarly to human brain-like ECs and EECM-BMEC-like cells, cARLA not only increased the staining intensity of claudin-5 (**Fig. 1M**) but also its junctional localization (**Fig. 1N**) in both genotypes. TEER was elevated by cARLA compared to the control group in both genotypes, but approximately 50% less so in *Cldn5^+/-^* cultures (**Fig. 1O**), suggesting that cARLA increases TEER by inducing *Cldn5* gene expression.

We repeated the main experiments in additional, widely used BBB culture models of rodent origin to see if the synergistic effect of cARLA is conserved between species. For this purpose, we selected the mouse brain EC line bEnd.3 in monoculture and a rat primary brain EC-pericyte-astrocyte co-culture model^52^, which considerably differ in their complexity (**Fig. 2A**). Barrier integrity measured by impedance was robustly elevated by cARLA in both models, albeit to a different extent and with different kinetics (**Fig. 2B,C**). Notably, the optimal treatment concentrations of cARLA and its components were different between mouse bEnd.3 cells (**SI Appendix, Fig. S7A-F**), primary rat brain ECs (**SI Appendix, Fig. S8A-F**) and human brain-like ECs. Therefore, we emphasize the need for optimizing these parameters first when testing cARLA in a new BBB model. As validation, we measured a 2.1-fold increase in TEER (**Fig. 2D**) and decreased permeability of fluorescein (1.6-fold) and albumin (3.4-fold) tracers across bEnd.3 monolayers upon cARLA treatment (**Fig. 2E-F**). In the rat primary co-culture model, a 2.9-fold increase in TEER (**Fig. 2G**) as well as a 2.1-fold and 1.93-fold decrease in permeability was seen for fluorescein and albumin, respectively (**Fig. 2H-I**). Similarly to human brain-like ECs, treatment with cARLA synergistically elevated the staining intensity of claudin-5 in both mouse bEnd.3 cells (**Fig. 2J)**, and in rat primary brain ECs (**Fig. 2K)**, but did not change the staining intensity of ZO-1 (**SI Appendix, Fig. S7G, S8G**). Taken together, our results reveal that cARLA synergistically enchances barrier tightness in a range of BBB models and this effect is dependent on claudin-5.

**Figure 2.**
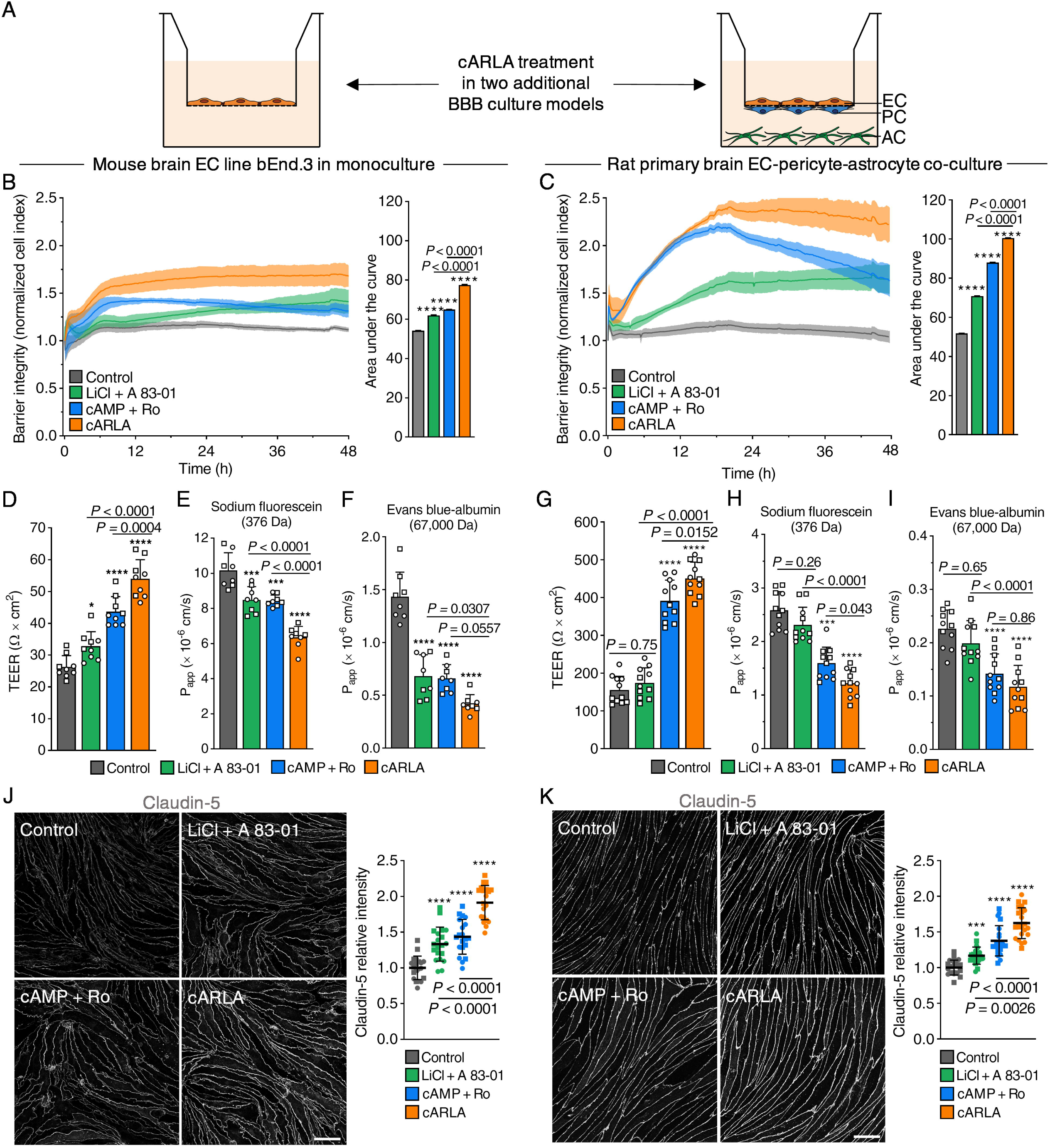
The effect of cARLA on barrier tightness is reproducible in additional BBB culture models. **A**) Schematic drawing of additional BBB culture models in which cARLA treatment was tested. **B)** Real-time measurement of barrier integrity by impedance in mouse bEnd.3 cells and **C)** in rat primary brain capillary ECs. Higher normalized cell index values and a higher area under the curve indicates increased barrier integrity. Mean ± SD, ANOVA with Bonferroni’s post-hoc test, \*\*\*\**P*<0.0001 compared to the control group, *n*=5-6 for both panels. **D)** Measurement of TEER as well as permeability of **E)** sodium fluorescein and **F)** Evans blue-albumin across mouse bEnd.3 monolayers after 48h treatment. Similarly, **G)** TEER as well as permeability of **H)** sodium fluorescein and **I)** Evans blue-albumin was measured across the rat primary cell-based co-culture BBB model after 24h treatment. Mean ± SD, ANOVA with Bonferroni’s post-hoc test, \**P*<0.05, \*\*\**P*<0.001, \*\*\*\**P*<0.0001 compared to the control group, *n*=8 in mouse bEnd.3 cells *n*=11 in the rat primary BBB model from 2-2 independent experiments. **J)** Claudin-5 immunostaning in mouse bEnd.3 cells and **K)** in the rat primary BBB model. Bar: 50 µm in both panels. Mean ± SD, ANOVA with Bonferroni’s post-hoc test, \*\*\*\**P*<0.0001 compared to the control group, *n*=20-22 from 2 independent experiments in both panels.

### cARLA promotes barrier maturation and brain endothelial identity revealed by MACE-seq

To investigate gene expressional changes in human brain-like ECs upon cARLA treatment, we performed Massive Analysis of cDNA Ends (MACE-seq, **Fig. 3A**). Principal component analysis revealed a sharp separation of control and cARLA samples along principal component 1 (PC1), with low heterogeneity of replicates within each group (**Fig. 3B**). In line with our findings on barrier integrity, *CLDN5* was the most differentially expressed gene based on statistical significance (**SI Appendix, S9A**) among the 619 upregulated and 546 downregulated genes upon cARLA treatment (**Fig. 3C**). Key transcripts related to different aspects of BBB function were also differentially expressed (**Fig. 3C**). For example, treatment with cARLA upregulated *ABCB1* encoding the efflux pump P-glycoprotein, *ST6GALNAC3,* an enzyme involved in the synthesis of the EC glycocalyx, and *SLC2A1,* the glucose transporter GLUT1. Conversely, genes involved in pathological BBB disruption, including *VCAM1* (vascular cell adhesion molecule-1), *CCL2* (the C-C motif ligand-2 chemokine) and *SERPINE1* (plasminogen activator inhibitor-1) were downregulated by cARLA (**Fig. 3C**).

**Figure 3.**
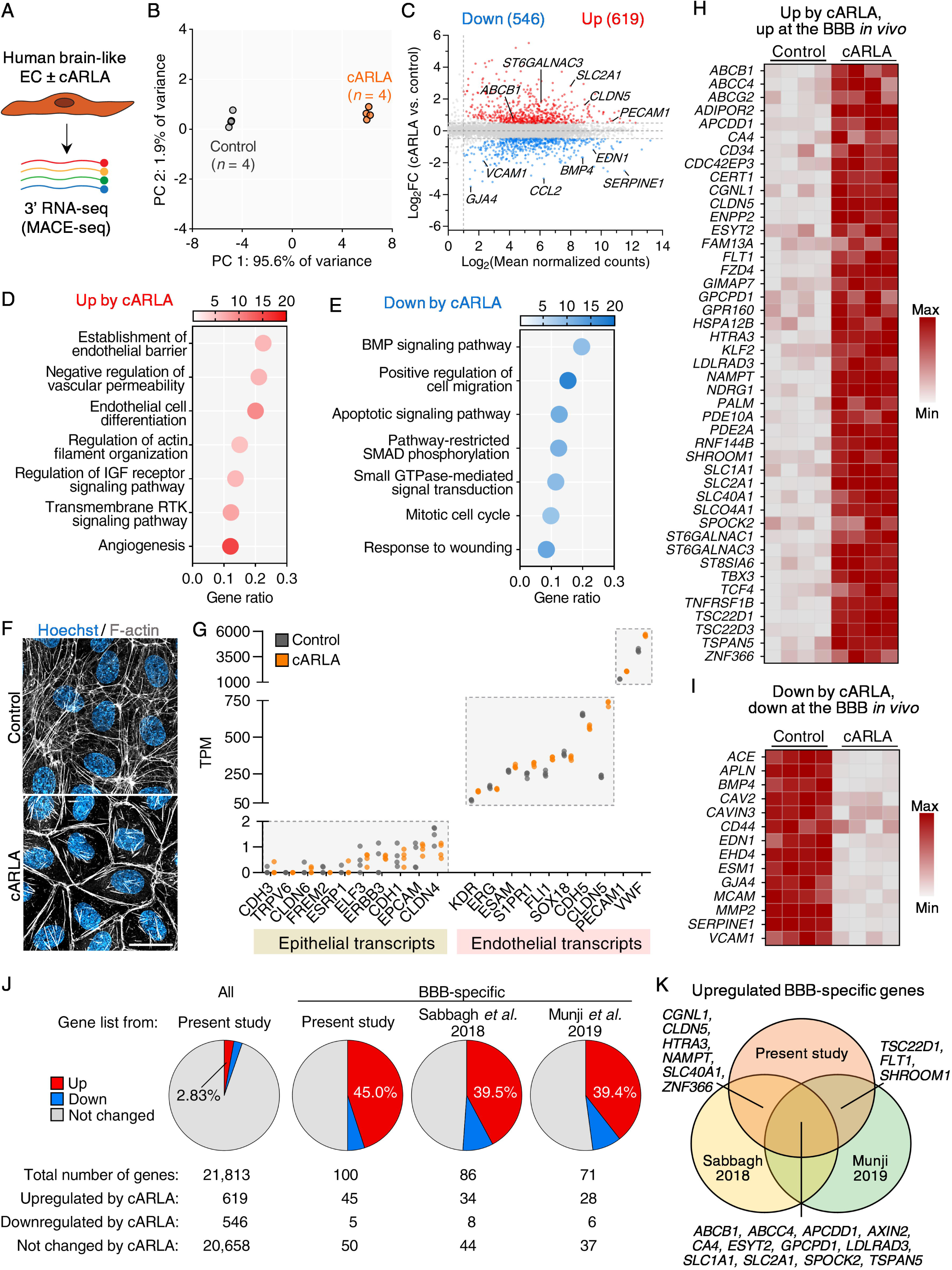
cARLA promotes barrier maturation and brain endothelial identity in human brain-like ECs. **A**) Schematic drawing of the experimental setup. **B)** Principal component analysis (PCA) of control and cARLA-treated samples. **C)** Mean-difference (MD) plot of cARLA vs. control samples showing up– and downregulated genes and their expression levels. Key transcripts related to different aspects of BBB function are highlighted. **D**) Functional enrichment analysis of upregulated– and **E)** downregulated gene sets upon cARLA treatment. Gene Ontology Biological Process (GO:BP) terms are ranked based on their gene ratio and are colored based on statistical significance. **F)** F-actin immunostaining, with or without cARLA treatment. Bar: 40 µm. **G)** Validation of the vascular endothelial nature of human stem cell-derived brain-like ECs. Expression levels (TPM: transcript per million) of key epithelial and endothelial transcripts from our dataset are plotted on a 3-segment y-axis covering multiple orders of magnitude. **H)** Scaled heatmap of 45 BBB-specific genes that are upregulated by cARLA and are enriched in brain ECs *in vivo*. **I)** Scaled heat map of 14 genes that are downregulated by cARLA and are enriched in peripheral ECs *in vivo*. **J)** The ratio of upregulated, not changed and downregulated genes by cARLA for all genes and BBB-specific genes using our and previously published^4,53^ lists. **K)** Venn diagram of BBB-specific genes from the three lists upregulated by cARLA.

On a pathway level, we observed an overrepresentation of genes related to EC differentiation and barrier formation as well as quiescence and maturation (**Fig. 3D,E**). Accordingly, cARLA reduced the staining intensity of cytoplasmic vimentin fibers (**SI Appendix, Fig. S10A**), and the number of Ki-67^+^ cells (**SI Appendix, Fig. S10B**) in EC monoculture. This was accompanied by a marked redistribution of F-actin from stress fibers to the cortical actin cytoskeleton (**Fig. 3F; SI Appendix, Fig. S11A**) and a more continous ZO-1 staining at cell borders (**SI Appendix, Fig. S11B**), supporting the maturation of cell-cell contacts. Other upregulated junctional genes include *F11R* (JAM-A), *CGNL1* (paracingulin/JACOP), and *PECAM1* (CD31; **SI Appendix, Fig. S11C-E**).

Notably, the expression of key endothelial transcripts in human brain-like ECs was orders of magnitude higher than that of epithelial-associated transcripts^27^, confirming the vascular endothelial nature of the model (**Fig. 3G**). To test if cARLA further shifts the gene expression of ECs towards the *in vivo* brain EC signature, we examined a set of 100 transcripts (**SI Appendix, Table S1, Fig. S9B**) that are enriched in brain *vs*. peripheral ECs in mice^4,50,53^ and are robustly expressed at the human brain endothelium^54,55,69,70^. We observed an upregulation of 45 BBB-specific genes by cARLA, including BBB-enriched transporters, enzymes and transcription factors (**Fig. 3H**). By contrast, 14 BBB-depleted genes participating in cell-cell adhesion, proliferation, and vesicular transport, were downregulated by cARLA (**Fig. 3I**). The ratio of BBB-specific genes upregulated by cARLA was 45% according to our list of 100 genes, and 39.4-39.5% using two other published gene lists from mice^4,53^(**Fig. 3J,K; SI Appendix, Table S2**). This indicates that BBB-specific genes were 14-16× more likely to be upregulated by cARLA compared to all genes (2.8% upregulated, **Fig. 3J**). The effect of cARLA positively correlated with the gene pattern differences observed between mouse brain *vs*. peripheral ECs^53^, but only partially matched the differences between mouse brain vs. lung ECs^4^ (**SI Appendix, Fig. S12A**). Importantly, for BBB-specific genes, cARLA shifted the expression profile of human brain-like ECs *in vitro* towards values in ECs from isolated human brain microvessels^70^ (**SI Appendix, Fig. S12B**). Together, these results demonstrate that cARLA promotes quiescence, barrier maturation, and the acquisition of a brain EC-like identity.

### cARLA induces a complex BBB phenotype at the mRNA, protein and functional level

We then verified our MACE-seq results at the protein and functional level for four selected BBB properties in human brain-like ECs (also shown by category in **SI Appendix, S13A-J**). First, genes encoding key enzymes that participate in the synthesis of the negatively charged EC glycocalyx, such as *ST6GALNAC3* and *ST8SIA6*, were upregulated by cARLA (**Fig. 4A**). This was also reflected by a higher intensity of wheat germ agglutinin (WGA) lectin staining, which recognizes negatively charged sialic acid residues in the glycocalyx, upon cARLA treatment (**Fig. 4B; SI Appendix, Fig. S14A**). Accordingly, we measured a more negative surface charge (Δ1.5 mV) in cARLA-treated ECs (**Fig. 4C**). Secondly, genes encoding BBB efflux transporters, such as P-glycoprotein (*ABCB1*), breast cancer-related protein (BCRP, *ABCG2*) and multidrug resistance-related protein 4 (MRP4, *ABCC4*), were upregulated by cARLA (**Fig. 4D**). At the protein level, we observed a 20-35% elevation in the staining intensity of P-glycoprotein (**Fig. 4E; SI Appendix, Fig. S15A**) which was accompanied by a marked increase in its luminal-abluminal polarization in ECs treated with cARLA (**SI Appendix, Fig. S15B**). BCRP protein levels were also robustly induced by cARLA, especially in EC monoculture (**SI Appendix, Fig. S15B**). As a functional test of efflux pump activity, we measured the permeability of rhodamine 123, a ligand of efflux pumps, across the co-culture model in blood-to-brain and brain-to-blood directions. The efflux ratio of rhodamine 123 was 2.1 upon cARLA treatment (a 1.6-fold increase compared to the control group), which could be reduced using verapamil, an efflux pump inhibitor (**Fig. 4F**). These results were also confirmed in accumulation assays using additional efflux pump ligands and inhibitors, with or without pericyte-conditioned medium (**SI Appendix, Fig. S15D-G**). Together, our data reveal a higher glycocalyx density and better polarity and functionality of efflux pumps in ECs treated with cARLA.

**Figure 4.**
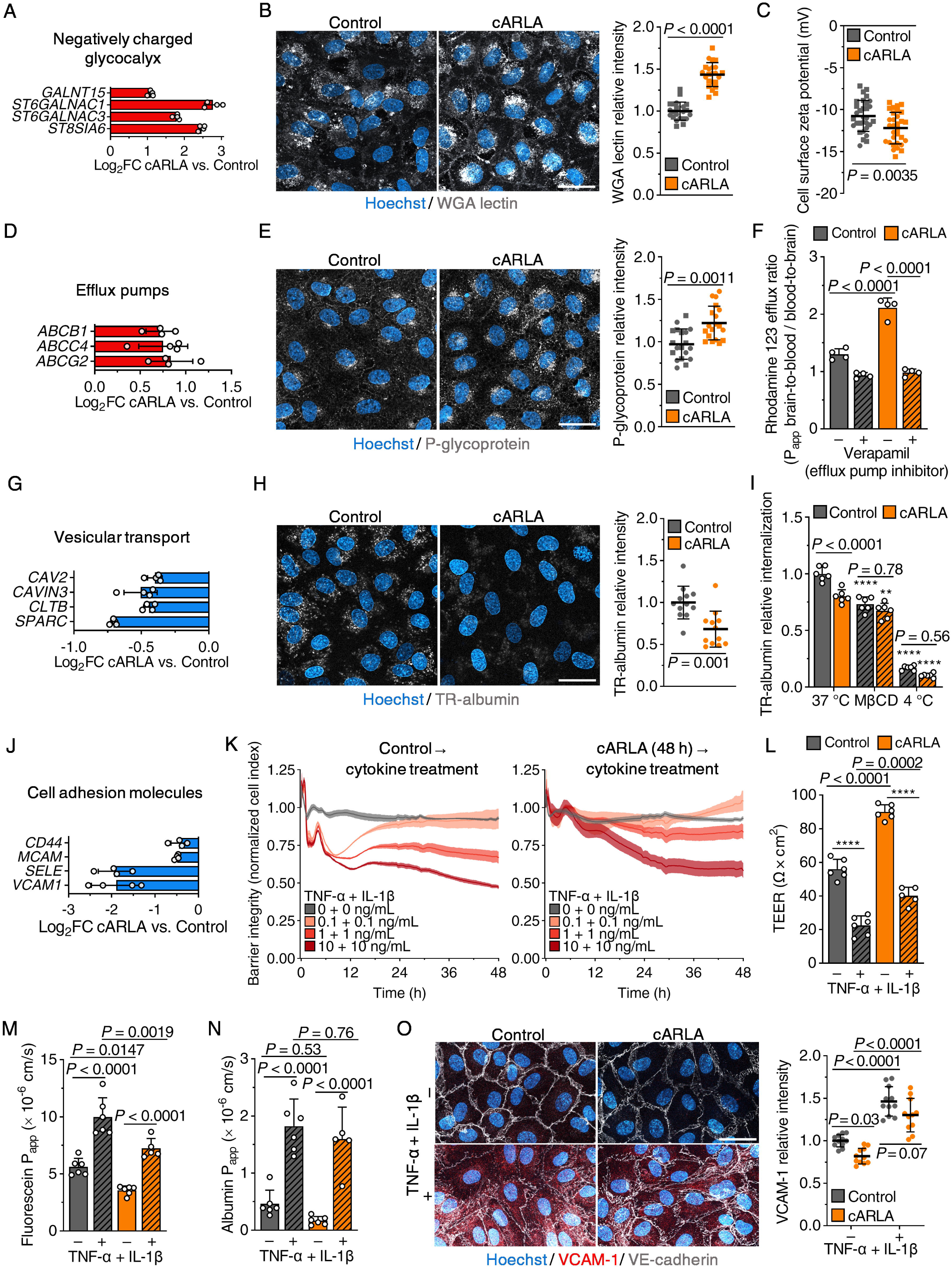
cARLA induces a complex BBB phenotype at the mRNA, protein and functional level in human brain-like ECs. **A**) Key glycocalyx synthesis enzymes are upregulated upon cARLA treatment. Mean ± SD. **B)** Wheat germ agglutinin (WGA) lectin staining labeling negatively charged sialic acid residues in the EC glycocalyx. Bar: 50 µm. Mean ± SD, unpaired t-test, *t*=11.43, *df*=43, *n*=22-24 from 2 independent experiments. **C)** Cell surface zeta potential measurement in ECs. Mean ± SD, unpaired t-test, *t*=3.045, *df*=60, *n*=31 from 2 independent experiments. **D)** Key BBB efflux transporters are upregulated upon cARLA treatment. Mean ± SD. **E)** P-glycoprotein (P-gp, *ABCB1*) immunostaining in ECs. Bar: 50 µm. Mean ± SD, unpaired t-test, *t*=4.065, *df*=36, *n*=19 from 2 independent experiments. **F)** Efflux ratio of rhodamine 123, a ligand of efflux pumps, across the BBB co-culture model in the presence or absence of verapamil, an efflux pump inhibitor. Mean ± SD, two-way ANOVA with Bonferroni’s post-hoc test, *n*=4. **G)** Key mediators of endocytic vesicle formation and nonspecific albumin uptake are downregulated by cARLA. Mean ± SD. **H)** Texas Red (TR)-labeled albumin internalization in ECs visualized by live cell confocal microscopy. Bar: 50 µm. Mean ± SD, unpaired t-test, *t*=3.786, *df*=22, *n*=12. **I)** TR-albumin internalization in ECs quantified by fluorescent spectrophotometry. MβCD: randomly methylated β-cyclodextrin, an inhibitor of lipid raft/caveolin-mediated endocytosis. Incubation at 4 °C inhibits energy-dependent internalization. Mean ± SD, two-way ANOVA with Bonferroni’s post-hoc test, \*\**P*<0.01, \*\*\*\**P*<0.0001 compared to its respective 37 °C group, *n*=6. **J)** Key immune cell adhesion molecules are downregulated by cARLA in the absence of inflammatory stimuli. Mean ± SD. **K)** Barrier integrity of control and cARLA-treated ECs upon TNF-α + IL-1β treatment, measured by impedance. Higher normalized cell index values indicate better-preserved barrier integrity. Mean ± SD, *n*=4-6. **L)** Measurement of TEER in control and cARLA-treated co-cultures upon TNF-α + IL-1β treatment. Mean ± SD, two-way ANOVA with Bonferroni’s post-hoc test, \*\*\*\**P*<0.0001 compared to non-stimulated ECs, *n*=6. **M)** Permeability of sodium fluorescein and **N)** Evans blue-albumin across the control or cARLA-treated co-culture model upon TNF-α + IL-1β treatment. Mean ± SD, two-way ANOVA with Bonferroni’s post-hoc test, *n*=5-6. **O)** VCAM-1 (red) and VE-cadherin (grey) immunostaining in control or cARLA-treated ECs, with or without TNF-α + IL-1β treatment. Mean ± SD, two-way ANOVA with Bonferroni’s post-hoc test, *n*=11.

On the other hand, genes associated with endocytic vesicle formation and nonspecific albumin uptake, suppressed pathways at the BBB *in vivo*, were also downregulated by cARLA (**Fig. 4G**). In line with this, we observed 20-30% lower levels of albumin internalization in ECs upon cARLA treatment (**Fig. 4H-I)**. Moreover, the internalization process was sensitive to methyl-β-cyclodextrin, which inhibits lipid raft/caveolin-mediated endocytosis, but this effect was smaller in cARLA-treated cells (**Fig. 4I**). Finally, genes encoding cell adhesion molecules involved in leukocyte trafficking across the inflamed BBB, such as *VCAM1* and E-selectin (*SELE*), showed a low expression at basal conditions (**SI Appendix, Fig. S13D**), and were downregulated by cARLA (**Fig. 4J**). When challenged by a combination of TNF-α and IL-1β in increasing concentrations, both control and cARLA-treated ECs responded to cytokine treatment by a decrease in impedance (**Fig. 4K**). ECs treated with cARLA maintained a higher barrier integrity at lower cytokine concentrations, but a robust reduction in impedance was observed at the highest cytokine concentration, similarly to the control group (**Fig. 4K**). Treatment with proinflammatory cytokines also decreased barrier integrity in the BBB model both in control and cARLA groups, as measured by TEER (**Fig. 4L**), fluorescein (**Fig. 4M**) and albumin permeability (**Fig. 4N**). In agreement with this and our MACE-seq results, VCAM1 protein levels without TNF-α and IL-1β treatment were low in ECs and were decreased by cARLA (**Fig. 4O, SI Appendix, Fig. S14B**), but were robustly increased in both control and cARLA-treated ECs by proinflammatory cytokines (**Fig. 4O**). These findings suggest that cARLA-treated ECs have lower basal rates of endocytosis, and retain their ability to respond to proinflammatory cytokines.

### Targeted pathways converge on Wnt/β-catenin signaling to mediate the effect of cARLA

We then asked how the synergistic effect of cARLA on barrier tightness is orchestrated in brain-like ECs. Bioinformatic analyses using the STRING database revealed a network of interactions between the effector transcription factors of cAMP, Wnt and TGF-β pathways, with β-catenin taking center stage (**Fig. 5A**). Indeed, we observed a robust induction of Wnt/β-catenin signaling-associated genes at the mRNA level (**Fig. 5B; SI Appendix Fig. S13E, S16A**). In addition, both the staining intensity (**Fig. 5C,D**) and nuclear accumulation of β-catenin (**Fig. 5E**) were synergistically elevated by cARLA, indicating a convergence of the targeted pathways on Wnt signaling. Active β-catenin levels were also increased by LiCl and A83-01 (**SI Appendix, Fig. S16B**), but the nuclear translocation of β-catenin was most evident in the cARLA group (**Fig. 5C)**. This was not accompanied by the disruption of junctional β-catenin or VE-cadherin (**SI Appendix, Fig. S17A**) complexes, and was consistent with the upregulation of *RAPGEF5*, a mediator of β-catenin nuclear import, by cARLA (**Fig. 5B**). Notably, the effect of cARLA was synergistic (**SI Appendix, Fig. S16E**) for 14 out of the 100 BBB-specific genes in our set, 6 of which are direct targets of Wnt signaling.

**Figure 5.**
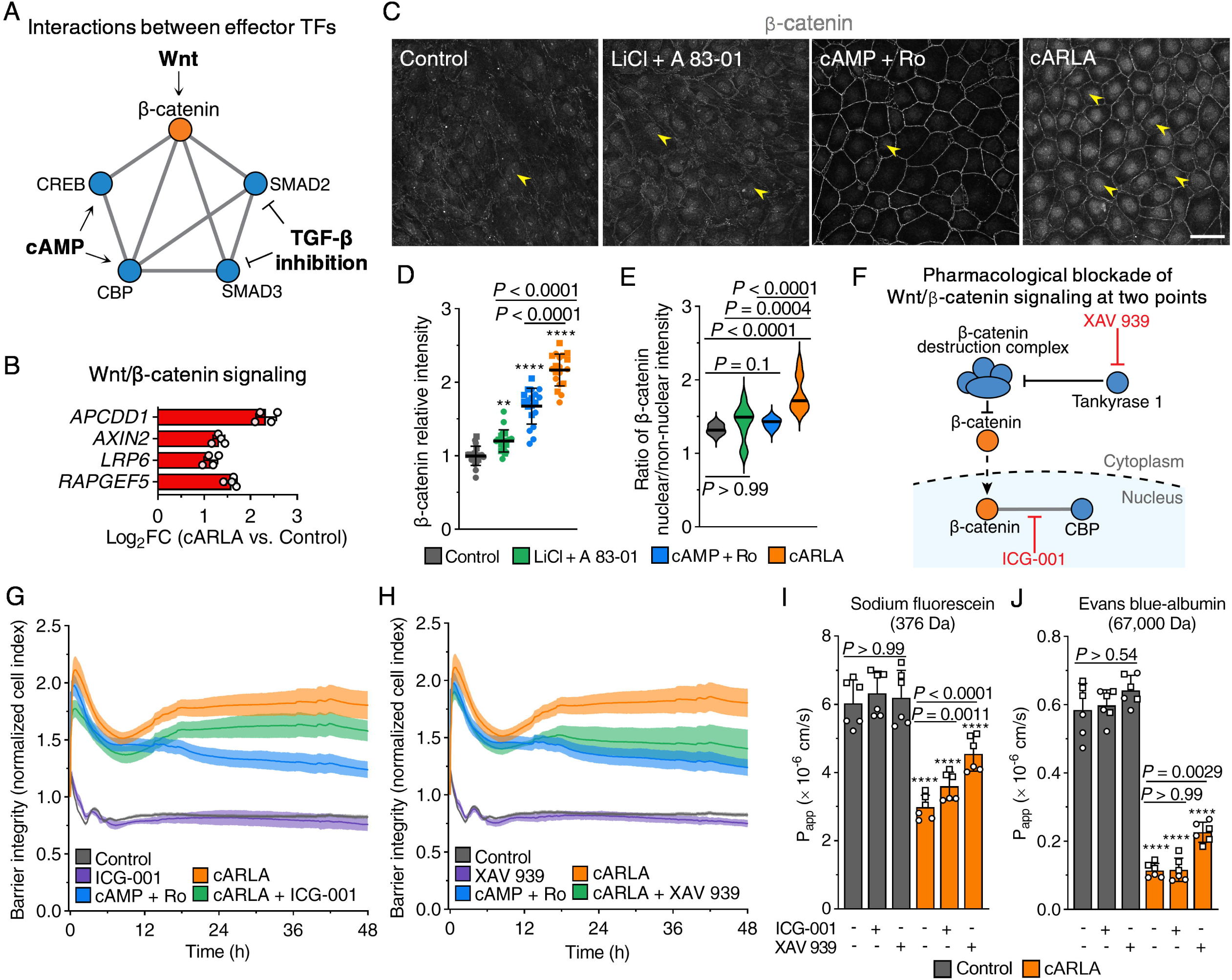
Targeted pathways converge on Wnt/β-catenin signaling to mediate the effect of cARLA. **A**) Schematic drawing of known interactions between effector transcription factors (TFs) of cAMP, Wnt and TGF-β signaling. **B** Canonical target genes of Wnt/β-catenin signaling are upregulated by cARLA. Mean ± SD. **C)** β-catenin immunostaining in ECs. Arrowheads point to nuclear β-catenin, a hallmark of active Wnt signaling. Bar: 50 µm. **D)** Quantification of β-catenin staining intensity and **E)** β-catenin nuclear/non-nuclear ratio. Mean ± SD for intensity, median ± quartiles for nuclear/non-nuclear ratio, ANOVA with Bonferroni’s post-hoc test, \*\**P*<0.01, \*\*\*\**P*<0.0001 compared to the control group, *n*=20 from 2 independent experiments. **F**) Schematic drawing of the mechanism of action of Wnt signaling inhibitors ICG-001 and XAV 939. **G)** Barrier integrity measured by impedance in EC monolayers, with or without ICG-001 (5 µM) and **H)** XAV 939 (1 µM). Higher normalized cell index values indicates higher barrier integrity. Mean ± SD, *n*=5-6. **I)** Permeability of sodium fluorescein and **J)** Evans blue-albumin across the co-culture model, with or without ICG-001 and XAV 939. Mean ± SD, ANOVA with Bonferroni’s post-hoc test. \*\*\*\**P*<0.0001 in cARLA-treated ECs compared to its respective treatment in the control group, *n*=6 from 2 independent experiments.

To investigate how a highly active state of Wnt/β-catenin signaling contributes to the effect of cARLA, we pharmacologically blocked this pathway at two different points **(Fig. 5F**). ICG-001 specifically inhibits the β-catenin-CREB-binding protein (CBP) interaction in the nucleus, while XAV 939 blocks the pathway upstream of nuclear translocation. We used these inhibitors in concentrations that do not interfere with basal barrier integrity (**SI Appendix, Fig. S17B,C**) but block the effect of elevated levels of Wnt signaling on impedance (**Fig. 5G,H**). ICG-001 moderately decreased (**Fig. 5G**), whereas XAV 939 more potently reduced the effect of cARLA on barrier tightness, almost to the level of cAMP+Ro treatment (**Fig. 5H**). By contrast, the elevation in barrier integrity by cAMP+Ro alone was not affected by these inhibitors (**SI Appendix, Fig. S16C,D**). Treatment with XAV 939 increased the permeability of both fluorescein and albumin, whereas ICG-001 only increased the permeability of fluorescein but not albumin, across the cARLA-treated co-culture model (**Fig. 5I,J**). These findings suggest that the three pathways converge on Wnt/β-catenin signaling to mediate the effect of cARLA on barrier integrity, and that specifically, the β-catenin-CBP interaction is involved in establishing paracellular tightness.

### cARLA improves the prediction of drug– and nanoparticle transport across the human BBB

Finally, to put cARLA into practice, we assessed the penetration of ten clinically used small molecule drugs across the co-culture model, with or without cARLA treatment. Drugs were selected to represent different routes of penetration across the BBB, such as passive diffusion, efflux and influx transport (**SI Appendix, Table S3,4**). In blood-to-brain direction, the permeability of efflux pump ligands vinblastine, loperamide, salicylic acid and verapamil was lower upon cARLA treatment (**Fig. 6A**). Conversely, cARLA increased the permeability of tacrine (influx transport) and propranolol (passive diffusion, influx transport) across the co-culture model. As a consequence, the cARLA-treated human BBB model could correctly discriminate between drugs that readily enter the central nervous system *in vivo* and those that do not (**Fig. 6A**). Treatment with cARLA also elevated the efflux ratio of loperamide and methotrexate, indicating an improved vectorial transport across the BBB model (**Fig. 6B**). We observed similar results regarding the extent of drug penetration (**Fig. 6C**) given as *in vitro* unbound brain-to-plasma partition coefficients (K_p,uu,brain_)^22,25,56^. Importantly, cARLA increased the overall correlation between human *in vitro* K_p,uu,brain_ values and *in vivo* brain penetration data from rats^56^ and non-human primates^57^(**Fig. 6D**). In the case of verapamil, the only drug on our list with human clinical K_p,uu,brain_ data available, the cARLA-treated BBB model closely approximated the human *in vivo* value (**Fig. 6E, SI Appendix, Fig. S18A**), being second only to non-human primate *in vivo* data among predictive models. In cases of drugs where no human clinical data were available, we selected non-human primate values as reference points. For all drugs in this group, cARLA shifted *in vitro* K_p,uu,brain_ values towards non-human primate *in vivo* values, compared to the control BBB model (**SI Appendix, Fig. S18B**).

**Figure 6.**
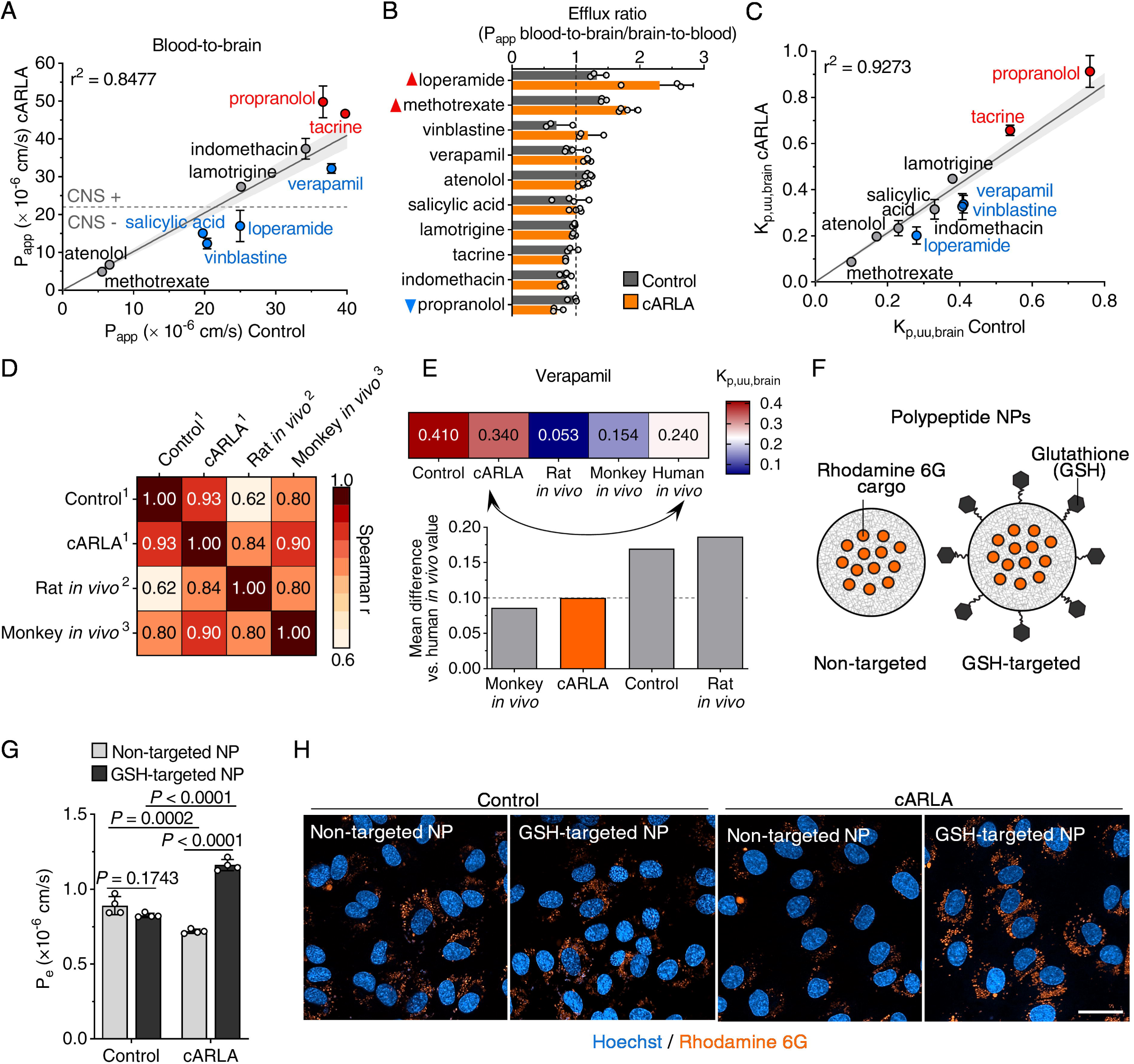
cARLA improves the *in vitro* prediction of drug– and nanoparticle delivery across the human BBB. **A**) Permeability of ten clinically used drugs in blood-to-brain direction across the BBB co-culture model, with or without cARLA treatment. CNS: central nervous system. Red and blue color indicates higher and lower permeability, respectively, for a given drug upon cARLA treatment (*P* < 0.05). Mean ± SD for symbols, *n*=3-4. Line of best fit with 95% convidence intervals and R^2^ value from a simple linear regression are shown. **B)** Drug efflux ratios in the co-culture model. Mean ± SD, multiple unpaired t-tests with Welch’s correction, *n*=3-4. Upward red and downward blue triangles indicate higher and lower efflux ratios, respectively (*P* < 0.05). **C)** *In vitro* unbound brain-to-plasma partition coefficient (K_p,uu,brain_) of drugs across the co-culture model, with or without cARLA treatment. Red and blue color indicates higher and lower values, respectively, for a given drug upon cARLA treatment (*P* < 0.05). Mean ± SD for symbols, *n*=3-4. Line of best fit with 95% convidence intervals and R^2^ value from a simple linear regression are shown. **D)** Correlation heatmap of *in vitro* K_p,uu,brain_ values for the whole set of drugs from our experiments with *in vivo* K_p,uu,brain_ data from rats and non-human primates. Numbers inside boxes are Spearman’s correlation coefficients. Data were obtained from: ^1^present study, ^2^Fridén *et al*., 2009 and ^3^Sato *et al*., 2021. **E)** Comparison of K_p,uu,brain_ measured *in vivo* in humans with its predictive models for the drug verapamil. Upper panel: heat map of K_p,uu,brain_ values. Lower panel: Mean difference of K_p,uu,brain_ values compared to human *in vivo* data. Lower values indicate better prediction. **F)** Schematic drawing of non-targeted and GSH-targeted polypeptide nanoparticles (NPs) carrying rhodamine 6G cargo. **G)** Permeability of non-targeted and GSH-targeted NPs across the co-culture BBB model, with or without cARLA treatment. P_e_: endothelial permeability coefficient. Mean ± SD, two-way ANOVA with Bonferroni’s post-hoc test, *n*=4. **H)** Internalization of non-targeted and GSH-targeted NPs carrying rhodamine 6G cargo (orange) in ECs, visualized by live cell confocal microscopy. Bar: 50 µm.

We also tested the effect of cARLA on the penetration of brain-targeted nanocarriers by assessing the permeability of polypeptide nanoparticles (NPs) decorated with or without the BBB-specific targeting ligand glutathione^58,59,60^ (GSH; **Fig. 6F**). In agreement with lower basal levels of endocytosis upon cARLA treatment (**Fig. 4G-I**), the permeability of non-targeted NPs was lower across the cARLA-treated BBB model compared to the control group (**Fig. 6G**). Notably, the BBB-specific targeting effect of GSH was apparent in the cARLA-treated but not in the control model, based on both the permeability (**Fig. 6G**) and internalization of NPs in ECs (**Fig. 6H**). Taken together, our results demonstrate that by inducing several aspects of BBB function, cARLA can improve the *in vitro* prediction of therapeutic drug– and NP transport across the human BBB.

## Discussion

In the past decade, several laboratories established BBB models derived from human stem cells^11,12,13,14,15,16,17,18,19,20,21,22,23,24,25^. Not only can this technology revolutionize drug development for the human brain by eliminating problems with species differences, it can also provide mechanistic insight into how the human BBB functions in health and disease. However, state-of-the-art human BBB models either have high barrier tightness but suffer from a mixed epithelial-endothelial identity or the cells are vascular ECs but have weak barrier properties^26,27^. This is a major limitation and thus, there is consensus in the field that existing differentiation protocols need serious improvements to better mimic both the endothelial nature and the complexity of the human BBB^26,61,62,63^. In this study, we describe an efficient, reproducible yet easy-to-use approach to induce complex BBB properties in vascular ECs *in vitro*.

The small molecule cocktail cARLA, which consists of pCPT-cAMP+Ro-20-1724+LiCl+A83-01, simultaneously activates cAMP and Wnt/β-catenin signaling, and inhibits the TGF-β pathway. So far, soluble factors targeting these pathways have been separately used to elevate BBB tightness *in vitro*. For example, cAMP elevating agents (pCPT-cAMP, Ro-20-1724, forskolin) rapidly increased junctional integrity in bovine and rodent primary brain ECs^47,48,49^. The activation of Wnt/β-catenin signaling^22,40,41,42,43^ by Wnt ligands (Wnt-3a, Wnt-7a/b) or small molecules (LiCl, 6-BIO, CHIR99021) as well as the inhibition of TG-β signaling by receptor antagonists (RepSox, A83-01)^44,45^ have also been shown to induce a subset of barrier properties individually. Yet we and others hypothesized that signaling cues do not act separately in ECs but rather interact with one another during BBB maturation. Indeed, overexpression of a combination of general and brain EC-specific transcription factors has been shown to increase barrier integrity in stem cell-derived ECs by ∼40%^65^. In contrast to targeting signaling pathways separately, Praça *et al*. used Wnt-3a, VEGF and retinoic acid to simultaneously activate multiple pathways during EC differentiation^15^. These factors, however, did not effectively potentiate each others’ effect on barrier tightness^21^. The strength of our approach compared to previous studies is that we performed a combinatorial screen of selected compounds for their ability to increase barrier tightness in ECs. This allowed us to identify a novel link between cAMP, Wnt/β-catenin and TGF-β signaling at the BBB *in vitro* and thus, we could rationally develop a small molecule cocktail with optimal composition to target this interaction. Treatment with cARLA has a greater than 2-fold, synergistic effect on barrier tightness *via* claudin-5, the dominant tight junction protein at the BBB. Based on our results, cARLA not only increases claudin-5 expression but can also promote its localization and stabilization at the intercellular junctions. We further support these findings by showing that the effect of cARLA is mediated by a convergence of targeted pathways on Wnt/β-catenin signaling.

The other main strength of our approach is that cARLA induces several aspects of BBB function, not just junctional tightness. Recapitulating these different BBB properties *in vitro* is necessary for both physiological studies and to predict the peneration of drugs and nanocarriers across the BBB. However, the induction of a complex BBB phenotype remains challenging in human stem cell-derived vascular ECs using current differentiation protocols^15,21,22,27^. Notably, cARLA does not induce epithelial-associated genes, and cARLA-treated ECs retain their characteristic endothelial response to proinflammatory cytokines. We also demonstrate that cARLA upregulates 39-45% of BBB-specific genes, and shifts the gene expressional profile of human stem cell-derived ECs towards the *in vivo* brain endothelial signature. As a limitation, 10 out of the selected 100 BBB-specific genes, including transcription factors *LEF1*, *FOXF2* and *ZIC3*, were not expressed in human brain-like ECs, and could not be induced by cARLA. Treatment with our small molecule cocktail also did not elevate the expression of 45 other BBB-specific genes, such as *MFSD2A* or *SLC38A5* (SNAT5). By contrast, cARLA induced a higher glycocalyx density and efflux pump activity as well as lower rates of transcytosis, which we validated at mRNA, protein and functional levels. This is important as pathological factors modify, and therapeutics interact with the EC glycocalyx, influx and efflux transporters as well as transcytotic pathways at the BBB. Accordingly, we observed a higher *in vitro-in vivo* correlation, and a substantially better prediction of the penetration of clinically used small molecule drugs in the cARLA-treated human BBB model compared to the control group and previously published human *in vitro* models. Among these, the effect of cARLA was best seen for drugs that are efflux pump ligands or have an influx transport mechanism across the BBB. Our data also indicate that cARLA increases the paracellular tightness of the human BBB model to a level that is suitable for testing larger biopharmaceuticals, such as antibody constructs, enzymes, and brain-targeted NPs.

Importantly, the aim of our study was not to establish a new human BBB model but rather to create an approach that can be used in a range of models across laboratories to move the field forward. To this end, we selected commercially available small molecules as an easy, affordable and rapid way to target developmentally relevant signaling pathways. Our method uses a single ‘hit-and-run’ treatment with cARLA for 24-48 hours to induce barrier maturation in ECs before the cells can be used in experiments. Notably, the effect of cARLA on barrier tightness is maintained after removal of the treatment, which can be important for several experimental setups. Whether the efficacy of cARLA can be further improved by other cues such as fluid flow and organ-on-a-chip technology^6,19,66^, 3D microenvironment^67^ or basement membrane composition^68^, remains to be explored. As we are aware of current difficulties in the field related to the induction of BBB properties of culture models in a reproducible way, we demonstrated the consistency of cARLA treatment using different batches of cells, measured by different experts in our research group. Moreover, we verified the effectiveness of cARLA in additional, well-established BBB models, namely EECM-BMEC-like cells differentiated from human induced pluripotent stem cells^21^, the mouse brain EC line bEnd.3 and primary brain EC-PC-AC co-culture models^58^ from mice and rats. Although it is a limitation of our study that the interaction between the three pathways targeted by cARLA was not yet proven *in vivo*, the synergistic effect of cARLA is conserved between species, and is reproducible across BBB models of different complexity.

Due to its high efficacy, reproducibility and ease of use, cARLA has the potential to be applied in a variety of BBB culture models across laboratories to advance drug development for the human brain. We envision that our study will also boost future research in understanding interactions between signaling pathways that govern the formation, maturation and maintenance of the BBB.

## Materials and Methods

### Derivation and culture of human stem cell-derived ECs

CD34^+^ hematopoietic stem cells were isolated from human umbilical cord blood and differentiated towards the endothelial lineage as previously described^22^. In accordance with the World Medical Association Declaration of Helsinki, informed consent was obtained from the donors’ parents and the protocol was approved by the French Ministry of Higher Education and Research (reference no. CODECOH DC2011-1321). Endothelial cells (ECs) were derived from CD34^+^ stem cells in endothelial cell growth medium (EGM; Lonza cat# CC-3162) supplemented with 20% fetal calf serum (FCS; Sigma-Aldrich cat# F7524) and 50 ng/mL vascular endothelial growth factor (VEGF_165_; PeproTech, cat# 100-20). After differentiation, ECs were cultured in endothelial cell culture medium (ECM; ScienCell, cat# 1001) supplemented with 5% fetal bovine serum (FBS; Sigma-Aldrich, cat# F4135), 1% endothelial cell growth supplement (ECGS; ScienCell, cat# 1052) and 50 µg/mL gentamicin (Sigma-Aldrich, cat# G1397). For experiments, ECs (passage number 6-7) were either grown in co-culture with or received conditioned medium from bovine brain pericytes (PCs) to acquire brain-like properties, as described in detail below. Cells were kept in a humidified incubator at 37 °C with 5% CO_2_.

### Impedance-based measurement of barrier integrity

Real-time, label-free measurement of barrier integrity was performed in 96-well plates with integrated gold electrodes (E-plate 96; Agilent, cat# 300600910) using an xCELLigence RTCA SP device (Agilent). Plates were coated with 0.2% gelatin (Sigma-Aldrich, cat# G2500) and the cell-free background was measured in culture medium for each well. Then, ECs were seeded in the plates at a density of 6×10^3^ cells/well and grown until fully confluent, while receiving 50% PC-conditioned medium to acquire brain-like properties. Confluent monolayers were treated with 8-(4-chlorophenylthio)adenosine 3′,5′-cyclic monophosphate sodium salt (cPT-cAMP; Sigma-Aldrich, cat# C3912, 25-250 µM) Ro 20-1724 (Sigma-Aldrich, cat# 557502, 17.5 µM), recombinant human Wnt-3a protein (R&D, cat# 5036-WN, 10-300 ng/mL), recombinant human Wnt-7a protein (PeproTech, cat# 120-31, 10-300 ng/mL), lithium chloride (LiCl; Sigma-Aldrich, cat# L9650, 3-30 mM), RepSox (PeproTech, cat# 4463325, 1-30 µM), A83-01 (PeproTech, cat# 9094360 or Tocris, cat# 2939, 0.1-3 µM) and the impedance of monolayers was measured at 10 kHz at every 5 min for 48-96 h. The readout of impedance measurements is cell index, a dimensionless parameter which correlates with the strength of cell-cell contacts and cell adhesion. Cell index was defined as Z_n_−Z_0_, where Z_n_ is the impedance at time point n and Z_0_ is the impedance of the cell-free background in each well. Cell index values were normalized to the last time point before treatment.

### Optimal concentrations of soluble factors and cARLA in human ECs

In human stem cell-derived brain-like ECs, the following concentrations were optimal according to our screen and were used for treatments: pCPT-cAMP 250 µM, Ro-20-1724 17.5 µM, Wnt-3a 300 ng/mL, Wnt-7a 300 ng/mL, LiCl 3 mM, RepSox 10 µM and A83-01 3 µM. For this cell type, cARLA treatment consisted of 250 µM pCPT-cAMP + 17.5 µM Ro-20-1724 + 3 mM LiCl + 3 µM A83-01. The optimal duration of cARLA treatment in human stem cell-derived brain-like ECs was found to be 48 h. The optimal composition and duration of cARLA treatment in mouse bEnd.3 cells as well as primary mouse and rat brain ECs are described below.

### Construction of the human brain-like EC and bovine brain PC co-culture BBB model

Primary bovine brain PCs were isolated, transfected and maintained as previously described^22^. PCs were initially cultured in 0.2% gelatin-coated 100 mm Petri dishes in Dulbecco’s Modified Eagle Medium (DMEM; Gibco, cat# 11885084) supplemented with 20% FBS (Sigma-Aldrich, cat# F4135), 1% GlutaMAX (Gibco, cat# 35050061) and 50 µg/mL gentamicin (Sigma-Aldrich, cat# G1397). PC-conditioned medium from 48 h cultures was sterile-filtered (0.2 µm) and used in a 1:1 ratio with EC medium to promote the acquisition of brain-like EC properties in experiments where co-culture was not suitable. For all other experiments involving human stem cell-derived ECs, a contact co-culture BBB model was used. PCs were gently trypsinized and seeded at the bottom side of collagen IV-coated (Sigma-Aldrich, cat# C5533, 100 µg/mL) Transwell inserts (Corning, cat# 3401 (12 mm inserts) and 3413 (6.5 mm inserts), polycarbonate membrane, 0.4 µm pore size) at a density of 1.8×10^4^ cells/cm^2^. PCs were allowed to attach to the bottom side of the insert (turned upside down) for 3 h at 37 °C, then the insert was placed in a 12 or 24-well plate containing EC medium. ECs were then seeded to the other, collagen IV (100 µg/mL) and fibronectin-coated (Sigma-Aldrich, cat# F1141, 25 µg/mL) side of the insert at a density of 7×10^4^ cells/cm^2^ in EC medium (ECM supplemented with 5% FBS and 1% ECGS, as described above). 48 h after the assembly of the co-culture, medium was replaced in both compartments. Four days after the assembly of the co-culture, medium was replaced in the bottom compartment and the upper, luminal (‘blood side’) compartment was treated with control EC medium or cARLA or its components for 48 h. Experiments were performed 6 days after the assembly of the co-culture and 48 h after treating the BBB model with soluble factors or their combinations.

### Transendothelial electrical resistance

Transendothelial electrical resistance (TEER) was measured first across EC monolayers grown in 50% PC-conditioned medium on 6.5 mm Transwell inserts as described above using a CellZScope+ instrument (nanoAnalytics). Confluent monolayers were treated with control medium, cARLA or its components and the impedance spectra of ECs between 1 Hz and 100 KHz was monitored every 3 h over a 96 h period at 37 °C. TEER was automatically calculated by the device and is presented in arbitrary units relative to baseline. TEER was also measured across EC-PC co-cultures using an EVOM voltohmmeter (World Precision Instruments) equipped with a hand-held STX2 chopstick electrode (for 12 mm inserts) or an EndOhm 6G chamber electrode (for 6.5 mm inserts). Electrodes were equilibrated in pre-warmed EC medium for 15 min and TEER was measured on a 37 °C heating pad. TEER in Ω×cm^2^ was calculated by subtracting the mean resistance of two coated, cell-free inserts from raw resistance values, multiplied by the surface area of the insert.

### Tracer permeability

After 48 h luminal (‘blood side’) treatment, the upper, donor compartment of the co-culture model was incubated with tracers sodium fluorescein (376 Da, Sigma-Aldrich, cat# 46960, 10 μg/mL) and Evans blue-albumin complex (67 kDa, 10 mg/mL bovine serum albumin (BSA) + 167.5 μg/mL Evans blue dye) in phenol red-free DMEM/F-12 medium (Gibco, cat# 21041025) supplemented with 1% FBS (Sigma-Aldrich, cat# F4135). Cultures were incubated on a PSU-2T horizontal shaker (Biosan, 150 rpm) for 1 h at 37 °C. Samples from both compartments were collected and the mean fluorescence of tracers was measured using a Fluorolog 3 spectrofluorometer (Horiba Jobin Yvon) with excitation/emission wavelengths 485/515 nm for sodium fluorescein and 584/663 nm for Evans blue-albumin complex. Concentration of tracers were determined by standard calibration curves. Apparent permeability coefficients (P_app_) were calculated as previously described (Veszelka *et al*., 2018, PMID: 29872378) using the following equation:

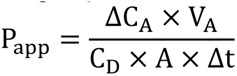

where C_A_ is the concentration measured in the acceptor compartment at 1 h, V_A_ is the volume of the acceptor compartment (1.5 and 0.9 mL for 12 and 6.5 mm inserts, respectively), C_D_ is the concentration in the donor compartment at 0 h, A is the surface area available for permeability (1.12 and 0.33 cm^2^ for 12 and 6.5 mm inserts, respectively) and Δt is the duration of the experiment (1 h).

### Immunocytochemistry and image analysis

After 48 h luminal (‘blood side’) treatment, the medium was removed from culture inserts and ECs were fixed with a 1:1 mixture of ice cold methanol-acetone solution for 2 min (for claudin-5, ZO-1 and VE-cadherin staining) or with 3% paraformaldehyde (all other stainings) for 15 min at room temperature. Cells were blocked with 3% BSA in phosphate-buffered saline (PBS; KCl 2.7 mM, KH_2_PO_4_ 1.5 mM, NaCl 136 mM, Na_2_HPO_4_×2 H_2_O 6.5 mM, pH 7.4) for 1 h at room temperature and were subsequently incubated with primary antibodies rabbit anti-claudin-5 (Sigma-Aldrich, cat# SAB4502981, AB_10753223, 1:300), mouse anti-claudin-5 (Invitrogen, cat# 35-2500, AB_2533200, 1:200), rabbit anti-ZO-1 (Invitrogen, cat# 61-7300, AB_2533147, 1:400), mouse anti-P-glycoprotein (Sigma-Aldrich, cat# 517310, AB_564389, 1:100), rabbit anti-P-glycoprotein (Abcam, cat# ab170904, AB_2687930, 1:100), mouse anti-α-SMA (Sigma-Aldrich, cat# A2547, AB_476701, 1:300), rabbit anti-PDGFRβ (Abcam, cat# ab32570, AB_777165, 1:200), goat anti-CD13 (R&D Systems, cat# AF2335, AB_2227288, 1:200), mouse anti-vimentin (Invitrogen, cat# MA511883, AB_10985392, 1:200), rabbit anti-Ki-67 (Millipore, cat# AB9260, AB_2142366, 1:200), rabbit anti-PECAM-1 (Santa Cruz, cat# sc-1506-R, AB_831096, 1:200), rabbit anti-β-catenin (Sigma-Aldrich, cat# C2206, AB_476831, 1:200), mouse anti-VCAM-1 (Invitrogen, cat# 13-1060-82, AB_529468, 1:100) and rabbit anti-VE-cadherin (Invitrogen, cat# PA519612, AB_10979589, 1:200) diluted in 3% BSA-PBS overnight at 4 °C. For intracellular antigens (F-actin, caveolin-1 and nuclear β-catenin), a permeabilization step with 0.2% Triton X-100 in PBS for 10 min at room temperature preceded blocking. Cells were then incubated with secondary antibodies Alexa Fluor 488 goat anti-rabbit IgG (Thermo Fisher, cat# A-11008, AB_143165, 1:400), Alexa Fluor 555 goat anti-rabbit IgG (Thermo Fisher, cat# A-21428, AB_2535849, 1:400) and Alexa Fluor 647 goat anti-mouse IgG (Thermo Fisher, cat# A-21235, AB_2535804, 1:300), the F-actin probe Alexa Fluor 488 phalloidin (Invitrogen, cat# A12379, 1:100) and the nuclear counterstain Hoechst 33342 (Thermo Fisher, cat# H1399, 1 μg/mL) diluted in PBS for 1 h at room temperature, protected from light. Between each step, cells were washed three times in PBS. Inserts were mounted on glass slides with Fluoromount-G (Southern Biotech, cat# 0100-01) and were examined using a Leica TCS SP5 AOBS confocal laser scanning microscope (Leica Microsystems) equipped with HC PL APO 20× (NA=0.7) and HCX PL APO 63× oil (NA=1.4) objectives. Z-stacks with approximately 0.75 μm spacing were acquired. Maximum intensity projections of each channel were analyzed using the ‘Mean gray value’ function in Fiji (ImageJ) software by an observer blinded to the experimental groups. Claudin-5 relative continuity was quantified in MATLAB software (The Mathworks Incorporated). For this, images were first corrected for intensity inhomogeneities by a normalized two-dimensional cross-correlation (*normxcorr2*) run with a gaussian template of 2 pixel variance approximating the point spread function of the microscope. Then, image masks corresponding to junctional regions were determined by thresholding the corrected and normalized images with a threshold value of 0.25. Finally, the fragmentation of junctional regions was quantified by calculating the ratio of the total number of distinct regions in the mask to the total masked area. Continuity was defined as the reciprocal of fragmentation or simply the mean size of distinct regions in the image masks. β-catenin nuclear/non-nuclear ratio was also determined using MATLAB software. Areas corresponding to nuclear and cytoplasmic staining were determined by a simple threshold-based segmentation carried out on images of Hoechst 33342-stained nuclei of the same Z-stacks. Using the obtained segmentation masks, mean fluorescence pixel intensity of β-catenin staining was calculated for areas corresponding to the nuclei and cytoplasm.

### Differentiation and culture of EECM-BMEC-like cells and SMLCs

EECM-BMEC-like cells and SMLCs were derived according to the protocol of Nishihara *et al*., 2020^21^ (described in detail in Nishihara *et al*., 2021, PMID: 34151293). Briefly, CD31^+^ endothelial progenitor cells (EPCs) were differentiated from the IMR90-4 human stem cell line (WiCell). Human induced pluripotent stem cells (hiPSCs) were seeded into Matrigel (Corning, cat# 354230)-coated plates in mTeSR1 medium (Stemcell Technologies, cat# 85850) supplemented with Y-27632 (Tocris, cat# 1254, 5 µM, 24 h, day –3). On days 0 and 1, the basal medium was changed to LaSR medium (Advanced DMEM/F-12, Life Technologies, cat# 12634, GlutaMAX, Thermo Fisher, cat# 35050-061, 2.5 mM, and L-ascorbic acid, Sigma-Aldrich, cat# A8960, 60 μg/mL) supplemented with CHIR99021 (Selleck, cat# S1263, 8 μM, 48 h). On day 5, CD31+ EPCs were purified using a FITC-conjugated human CD31 antibody (Miltenyi, cat# 130-117-390, clone AC128) and the EasySep Human FITC Positive Selection Kit II (Stemcell Technologies, cat# 18558). Purified EPCs were seeded into collagen IV (Sigma-Aldrich, cat# C5533, 100 µg/mL)-coated 6-well plates in hECSR medium (Human Endothelial SFM, Thermo Fisher, cat# 11111-044; B-27 Supplement (50×), Thermo Fisher, cat# 17504044; human fibroblast growth factor 2 (bFGF), Tocris, cat# 233-FB-500, 20 ng/mL) supplemented with Y-27632 (5 µM, 24 h). On the next day, culture medium was switched to hECSR without Y-27632, and was changed every other day. Upon reaching 80-100% confluency, ECs were selectively passaged using Accutase (Sigma-Aldrich, cat# A6964). After two selective passages, pure EC cultures were obtained. SMLCs left at the bottom of 6-well plates after selective passaging were cultured in DMEM/F-12 medium (Thermo Fisher, cat# 11320074) supplemented with 10% FBS (Thermo Fisher, cat# 10270106). To assemble non-contact BBB co-culture models, SMLCs were seeded into collagen IV-coated 12-well plates (Costar, cat# 3513) in hECSR medium, at a density of 1.8×10^4^ cells/cm^2^. Passage 3 ECs, referred to as EECM-BMEC-like cells, were seeded onto collagen IV (100 µg/mL) and fibronectin (Sigma-Aldrich, cat# F1141, 25 µg/mL)-coated Transwell inserts (Corning, cat# 3401, 12 mm, polycarbonate membrane, 0.4 µm pore size) in hECSR medium, at a density of 7×10^4^ cells/cm^2^. 48 h after the assembly of the co-culture, medium was replaced in both compartments. Four days after the assembly of the co-culture, medium was replaced in the bottom compartment, and the upper compartment was treated with control EC medium or cARLA (250 µM pCPT-cAMP + 17.5 µM Ro-20-1724 + 3 mM LiCl + 3 µM A83-01) for 48 h. Experiments were performed 6 days after the assembly of the co-culture and 48 h after treating the BBB model with cARLA.

### Animals

For primary cell isolation from mice, brain tissue was harvested from 18-24-week-old and newborn mice of both sexes according to the EU Directive 2010/63/EU about animal protection and welfare. Protocols were reviewed and approved by the Research Ethics Committees of Trinity College Dublin and/or the Royal College of Surgeons in Ireland under licenses from the Department of Health (HPRA: AE19136/PO80 and/or AE19127/P057), Dublin, Ireland. *Cldn5* heterozygous (*Cldn5^+/-^*) mice were generated at the the Department of Immunology, Genetics, and Pathology, Rudbeck Laboratory, Uppsala University, Sweden and were bred on-site at Trinity College Dublin onto a C57BL/6J background. Mice were housed under standard conditions (18–23 °C, 12 h light/dark cycle, 40-50% humidity) with regular rodent chow diet and water available *ad libitum*.

For primary cell isolation from rats, brain tissue was harvested from 3-week-old and newborn Wistar rats (Harlan Laboratories) of both sexes according to the regulations of the 1998. XXVIII. Hungarian law and the EU Directive 2010/63/EU about animal protection and welfare. Rats were housed in a conventional animal facility at the Biological Research Centre, Szeged, Hungary under standard conditions (22–24 °C, 12 h light/dark cycle) with regular rodent chow diet and water available *ad libitum*.

### Assembly of the primary mouse brain EC-PC-AC co-culture BBB model

The isolation of primary mouse brain capillary ECs, PCs and ACs was performed as previously described^57^. After isolation, ECs were seeded into 6-well culture plates (Corning, cat# 3516) coated with collagen IV (100 μg/mL) and fibronectin (25 μg/mL) and were cultured in DMEM/F-12 medium (Gibco, cat# 11320033) supplemented with 15% plasma-derived bovine serum (PDS; First Link, cat# 24-00-850), 10 mM HEPES (Sigma-Aldrich, cat# H4034), 1 ng/mL basic fibroblast growth factor (bFGF; Sigma-Aldrich, cat# F0291), 100 μg/mL heparin (Sigma-Aldrich, cat# 43149-100KU), 100× Insulin-Transferrin-Selenium reagent (ITS; Gibco, cat# 41400045) and 50 μg/mL gentamicin (Sigma-Aldrich, cat# G1397). During the first 3 days of culture, the medium of ECs also contained 4 μg/mL puromycin to eliminate P-glycoprotein negative, contaminating cell types. From day 3, PDS concentration in the medium of ECs was reduced to 10%. PCs were seeded into 6-well plates coated with collagen IV (100 μg/mL) and were cultured in DMEM (1 g/L glucose, Gibco, cat# 11885084) supplemented with 10% FBS (Sigma-Aldrich, cat# F4135) and 50 μg/mL gentamicin. ACs were seeded into 12-well plates (Corning, cat# 3513) coated with rat tail collagen (150 µg/mL) at a density of 8.5×10^4^ cells/cm^2^ and were cultured in the same medium as PCs. To assemble the triple co-culture model, PCs were passaged to the bottom side of Transwell inserts (Corning, cat# 3401, polycarbonate, 0.4 µm pore size) coated with collagen IV (100 μg/mL) at a density of 1.5×10^4^ cells/cm^2^. PCs were allowed to attach to the inserts for 3 h at 37 °C, then the insert was placed in a 12-well plate containing the ACs and the medium was replaced to EC medium. ECs were then seeded to the other, upper side of the culture insert coated with collagen IV (100 μg/mL) and fibronectin (25 μg/mL) at a density of 8.5×10^4^ cells/cm^2^ in EC medium. The three cell types were cultured together for 4 days before cARLA treatment. After that, the triple co-culture model was treated from the luminal (’blood side’) with cARLA. The optimal concentration of cARLA in the primary mouse BBB model was 125 µM pCPT-cAMP + 17.5 µM Ro-20-1724 + 1 mM LiCl + 3 µM A83-01 and the optimal treatment duration was 24 h. The measurement of TEER and claudin-5 immunocytochemistry was performed exactly as described above for human ECs.

### Mouse brain endothelial cell line bEnd.3

The mouse brain endothelial cell line bEnd.3 was purchased from ATCC (cat# CRL-2299) and cultured in DMEM/F-12 medium (Gibco, cat# 11320033) supplemented with 10% FBS (Sigma-Aldrich, cat# F4135) and 50 µg/mL gentamicin (Sigma-Aldrich, cat# G1397). Cells were used between passage numbers 25-28. For the measurement of barrier integrity by impedance, bEnd.3 cells were seeded at a density of 6×10^3^ cells/well in rat tail collagen-coated (150 µg/mL) 96-well plates (E-plate 96; Agilent, cat# 300600910) and confluent monolayers were treated as described above for human ECs. For the measurement of TEER and tracer permeability, bEnd.3 cells were passaged to Transwell inserts (Corning, cat# 3401 and 3413, polycarbonate, 0.4 µm pore size) at a density of 6.3×10^4^ cells/cm^2^ and grown in monoculture for 4 days. After that, bEnd.3 cells were treated from the luminal (’blood side’) with cARLA. The optimal concentration of cARLA in bEnd.3 cells was 125 µM pCPT-cAMP + 17.5 µM Ro-20-1724 + 10 mM LiCl + 1 µM A83-01 and the optimal treatment duration was 48 h. The measurement of TEER and tracer permeability as well as immunocytochemistry was performed exactly as described above for human ECs.

### Assembly of the primary rat brain EC-PC-AC co-culture BBB model

The isolation of primary rat brain capillary ECs, PCs and ACs was performed as previously described^52^. The assembly of the primary rat EC-PC-AC co-culture BBB model was performed exactly as described above for the primary mouse co-culture model, with the following modifications: (i) after isolation, rat ECs were seeded on 100 mm Petri dishes (instead of 6-well plates), (ii) during the first 3 days of culture, the medium of rat ECs contained 3 μg/mL puromycin (instead of 4 μg/mL), (iii) when assembling the model, rat ECs were seeded on the inserts at a density of 7×10^4^ cells/cm^2^ (instead of 8.5×10^4^ cells/cm^2^). For the measurement of barrier integrity by impedance, rat ECs were seeded at a density of 6×10^3^ cells/well in 96-well plates (E-plate 96; Agilent, cat# 300600910) coated with collagen IV (100 μg/mL) and fibronectin (25 μg/mL) and confluent monolayers were treated as described above for human ECs. The optimal concentration of cARLA in the primary rat BBB model was 250 µM pCPT-cAMP + 17.5 µM Ro-20-1724 + 1 mM LiCl + 1 µM A83-01 and the optimal treatment duration was 24 h. The measurement of TEER and tracer permeability as well as immunocytochemistry was performed exactly as described above for human ECs.

### Total RNA isolation

Human stem cell-derived brain-like ECs (in co-culture with PCs) were grown on Transwell inserts (Corning, cat# 3401) for 6 days. For the last 48 hours, ECs were treated luminally with cARLA or control medium containing the amount of solvents (Milli-Q water 100× and DMSO 1000× dilution) corresponding to cARLA. After treatment, PCs were removed and ECs were lysed directly with Buffer RLT Plus (Qiagen, cat# 1053393) on the inserts. For 1 sample in further analyses, cell lysates from 2 inserts were pooled. RNA was isolated using RNeasy Plus Micro Kit (Qiagen, cat# 74034) containing an integrated gDNA eliminator spin column according to the manufacturer’s protocol. RNA integrity was analyzed using automated capillary electrophoresis (RNA Pico Sensitivity Assay, LabChip GX II Touch HT instrument, Perkin Elmer). Samples were stored at –80°C until further analysis.

### Library preparation and 3’ RNA-sequencing (MACE-seq)

Genome-wide gene expression profiling was performed by massive analysis of cDNA ends (MACE-seq). For library preparation, samples with 1 µg of purified RNA were used. A total of 8 libraries (4 control and 4 cARLA) were constructed using the Rapid MACE-Seq kit (GenXPro GmbH, Frankfurt, Germany) according to the manufacturer’s protocol. Fragmented RNA was reverse transcribed into cDNA, using barcoded oligo(dT) primers containing TrueQuant unique molecular identifiers, followed by template switching. Library amplification was done using polymerase chain reaction (PCR), purified by solid phase reversible immobilization beads (Agencourt AMPure XP, Beckman Coulter, cat# A63882). Sequencing was performed on an Illumina NextSeq 500 platform.

### Bioinformatic analysis of MACE-seq data

Approximately 58 million single 75 bp MACE-seq reads were obtained across all 8 libraries. PCR duplicates were identified using TrueQuant technology and were removed from raw data. The remaining reads were further poly(A)-trimmed and low-quality reads were discarded. Clean reads were then aligned to the human reference genome (hg38, http://genome.ucsc.edu/cgi-bin/hgTables) using the Bowtie2 mapping tool, resulting in a dataset of 21,691 genes. Gene count data was normalized to account for differences in library size and RNA composition bias by calculating the median of gene expression ratios using the DESeq2 R/Bioconductor package. Testing for differential gene expression was also performed using DESeq2. As a result, *P*-value and log_2_FC (log_2_[fold change]) were obtained for each gene in the dataset. To account for multiple comparisons, false discovery rate (FDR) was calculated using the Benjamini-Hochberg method. Genes with an FDR < 0.01 and log_2_FC > 0.415 or log_2_FC < –0.415 were considered to be differentially expressed. To perform functional enrichment analysis of up– and downregulated gene sets upon cARLA treatment, the g:GOSt tool in g:Profiler was used to identify over-represented Gene Ontology Biological Process (GO:BP) terms and to calculate gene ratios and statistical significance for each pathway. Heatmaps were created using Min-Max scaling on normalized counts. For the correlation of gene expressional data with mouse^4,53^ and human^70^ *ex vivo* brain EC datasets, we plotted log_2_FC values as presented in the graphs, and calculated a global correlation coefficient (Pearson, two-tailed) using GraphPad Prism software (version 5.0). Our list of 100 BBB-specific genes is presented in **SI Appendix, Table S1**, whereas the list from Sabbagh *et al*.^53^ and Munji *et al*.^4^ are presented in **SI Appendix, Table S2**. RNA-sequencing datasets used for the correlations are available at the Gene Expression Omnibus (GEO): GSE95201 (Sabbagh *et al*.^53^), GSE111839 (Munji *et al*.^4^), GSE159851 (Schaffenrath *et al*.^70^, samples control_ 6, control_11, control_12, control_14, control_15, control_26).

### Glycocalyx staining and measurement of cell surface charge

After 48 h luminal (‘blood side’) treatment with cARLA or control medium, PCs were removed from culture inserts, human brain-like ECs were washed in PBS and fixed with 1% paraformaldehyde for 15 min at room temperature. Cells were then incubated with Alexa Fluor 488-conjugated wheat germ agglutinin (WGA) lectin (Invitrogen, cat# W11261, 5 mg/mL, 10 min at room temperature) that labels sialic acid and N-acetyl-d-glucosamine residues in the EC glycocalyx. Hoechst 33342 (Thermo Fisher, cat# H1399, 1 μg/mL) was used to counterstain nuclei. Cells were washed in PBS, mounted on glass slides with Fluoromount-G (Southern Biotech, cat# 0100-01) and were examined using a Leica TCS SP5 AOBS confocal laser scanning microscope equipped with a HCX PL APO 63× oil (NA=1.4) objective exactly as described above. Maximum intensity projections of Z-stacks from each channel were analyzed using the ‘Mean gray value’ function in Fiji (ImageJ) software by an observer blinded to the experimental groups. For the measurement of EC surface charge, EC-PC co-cultures were established exactly as described above, but on transparent Transwell inserts (Corning, cat# 3460, PET, 0.4 µm pore size) to be able to monitor the de-attachment of cells from the inserts. After 48 h treatment, ECs were washed with Ringer-HEPES solution (118 mM NaCl, 4.8 mM KCl, 2.5 mM CaCl_2_, 1.2 mM MgSO_4_, 5.5 mM d-glucose, 20 mM HEPES, pH 7.4), gently trypsinized and centrifuged at 700×*g* for 5 min. The cell pellet was resuspended in PBS supplemented with Ca^2+^ and Mg^2+^ (CaCl_2_ × 2H_2_O 1.2 mM, MgCl_2_ × 6H_2_O 0.5 mM) and zeta potential of 10^5^ cells per cuvette was measured using a Zetasizer Nano ZS instrument (Malvern) equipped with a He-Ne laser (λ=632.8 nm) as previously described^63^23.

### Measurement of efflux pump activity

After 48 h luminal (‘blood side’) treatment with cARLA or control medium, the permeability of rhodamine 123, a ligand of efflux pumps, was measured across the human brain-like EC-PC co-culture model in blood-to-brain and brain-to-blood directions. Either the luminal or abluminal (‘brain side’) compartment of the model was incubated with rhodamine 123 (Sigma-Aldrich, cat# R83702, 10 µM) diluted in phenol red-free DMEM/F-12 medium (Gibco, cat# 21041025) supplemented with 1% FBS (Sigma-Aldrich, cat# F4135) for 1 h at 37 °C on a horizontal shaker set to 150 rpm. To block efflux pump activity in some of the groups, verapamil (Sigma-Aldrich, cat# V4629, 2 µM) was added to the luminal compartment 30 minutes prior to rhodamine 123 treatment. After 1 h, samples from both compartments were collected and the mean fluorescence of rhodamine 123 was measured using a Fluorolog 3 spectrofluorometer (Horiba Jobin Yvon) with excitation/emission wavelengths 492/526 nm. Concentrations of rhodamine 123 were determined by standard calibration curves. Apparent permeability coefficients (P_app_) were calculated as described above.

Efflux ratio reflecting appropriate polarity of efflux pumps was calculated by dividing P_app_ values in brain-to-blood direction by P_app_ values in blood-to-brain direction.

We also performed accumulation assays with efflux pump ligands rhodamine 123 (5µM), doxorubicin (Sigma-Aldrich, cat# 44583, 5 µM) and Hoechst 33342 (Thermo Fisher, cat# H1399, 2 μg/mL), and efflux pump inhibitors cyclosporin A (CsA; Sigma-Aldrich, cat# C3662,10 µM) and Ko143 (Sigma-Aldrich, cat# K2144, 10 µM). Human stem cell-derived ECs were cultured with or without 50% PC-conditioned medium in 24-well plates (Corning, cat# CLS3527, flat-bottom) coated with collagen IV and fibronectin at a seeding density of 1.6×10^4^ cells/cm^2^. Confluent monolayers were treated with cARLA or control medium for 48 h. To block efflux pump activity, CsA or Ko143 were added to ECs 30 minutes prior to rhodamine 123, doxorubicin or Hoechst treatment. Efflux pump ligands were then added to ECs in the same medium containing the inhibitors in EC medium for 1 h at 37 °C on a horizontal shaker set to 150 rpm. After that, cells were washed three times with ice cold PBS supplemented with 0.1% BSA, once with acid stripping buffer (glycine 50 mM, NaCl 100 mM, pH 3.0) to remove cell surface-associated efflux pump ligands, and once with PBS. Finally, ECs were lysed with 1% Triton X-100 in Milli-Q water and the fluorescent signal of internalized efflux pump ligands were measured as described above with excitation/emission wavelengths 470/585 nm for doxorubicin and 360/497 nm for Hoechst. Concentrations of internalized efflux pump ligands were calculated using a standard calibration curve.

### Western blotting

For BCRP, human brain-like ECs were grown in monoculture or non-contact co-culture with PCs on Transwell inserts (cellQart, cat# 9300412, 24 mm, 0.4 µm pore size, PET, clear), and treated with cARLA exactly as described above. After 48 h cARLA treatment, ECs were lysed and fractionated using a subcellular protein fractionation kit (Thermo Fisher, cat# 78840), according to the manufacturer’s instructions. Membrane fractions were used for western blots. For active β-catenin, human brain-like ECs were grown in monoculture, with or without pericyte-conditioned medium, on 100 mm culture dishes (Corning, cat# 430167), and treated with cAMP+Ro, LiCl, A83-01 or cARLA exactly as described above. After 48 h treatment, ECs were lysed in RIPA buffer completed with PMSF, protease inhibitor cocktail and sodium orthovanadate according to the manufacturer’s instructions (RIPA Lysis Buffer System, Santa Cruz, cat# sc-24948). In this case, whole-cell lysates were used for further analyses. Western blots were performed according to Hoyk *et al.,* 2004 (PMID: 15390121), with minor modifications. A volume of 40 µL of each sample containing 25 µg protein was loaded in each lane of a 8 % sodium dodecyl sulfate-polyacrylamide gel. SDS-PAGE was carried out at 110 V for 2 h. Proteins were then transferred to PVDF membranes (Amersham™ Hybond®) for BCRP, or nitrocellulose membranes (Whatman®, Protran®) for active β-catenin detection at 260 mA for 2 h. The membranes were washed with 20 mM Tris-HCl, pH 7.5, 150 mM NaCl, 0.05% Tween-20 (TTBS, Sigma-Aldrich). Non-specific protein binding was blocked with 5% non-fat dry milk in TTBS for 1 h, then membranes were incubated overnight at 4 °C with rabbit anti-BCRP (Sigma-Aldrich, cat# ZRB1217, 1:1000) or mouse anti-active β-catenin (Sigma-Aldrich, cat# 05-665, 1:1000) antibodies. Following washing in TTBS, membranes were incubated for 2 h at room temperature with peroxidase-conjugated anti-mouse or anti-rabbit IgGs (Jackson, cat# 115-035-146 and 111-035-14, 1:10000), washed with TTBS and developed with Clarity Max Western ECL Substrate (Bio-Rad, cat# 1705062). As loading control, β-actin (Sigma-Aldrich, cat# A1978, 1:2000, overnight at 4°C) was used for each membrane. Membranes were analyzed with the Image Lab software version 6.0 (Bio-Rad Laboratories). Densities of the bands of interest from each membrane were normalized to β-actin values.

### Measurement of albumin internalization

Human brain-like ECs were cultured in 50% PC-conditioned medium on 35 mm glass bottom Petri dishes (Greiner Bio-One, cat# 627860) coated with collagen IV (Sigma-Aldrich, cat# C5533, 100 µg/mL) and fibronectin (Sigma-Aldrich, cat# F1141, 25 µg/mL) at a seeding density of 5×10^4^ cells/cm^2^. Confluent monolayers were treated with cARLA or control medium for 48 h. After that, culture medium was removed and cells were incubated with Texas Red-conjugated BSA (TR-albumin; Invitrogen, cat# A23017, 5 µg/mL) dissolved in control medium for 2 h at 37 °C. For the final 20 minutes of incubation, Hoechst 33342 (Thermo Fisher, cat# H1399, 2 μg/mL) was added to cells to counterstain nuclei. Cultures were washed twice in Ringer-HEPES supplemented with 1% FBS and live-cell imaging was performed immediately in the same buffer using a Leica TCS SP5 AOBS confocal laser scanning microscope equipped with a HCX PL APO 63× oil (NA=1.4) objective with the same parameters as described above. For measuring albumin internalization under inhibitory conditions, cells were cultured in 50% PC-conditioned medium in 24-well plates (Corning, cat# CLS3527, flat-bottom) coated with collagen IV and fibronectin at a seeding density of 1.6×10^4^ cells/cm^2^. Confluent monolayers were treated with cARLA or control medium for 48 h. To inhibit lipid raft/caveolin-mediated endocytosis, methyl-β-cyclodextrin (MβCD; Sigma-Aldrich, cat# C4555, 1 mM) was added to the medium of some groups for 1 h at 37 °C. Cells were then incubated with TR-albumin diluted in control EC medium containing 1% FBS for 2 h at 37 or 4°C. After that, cells were washed three times with ice cold PBS supplemented with 0.1% BSA, once with acid stripping buffer (glycine 50 mM, NaCl 100 mM, pH 3.0) to remove cell surface-associated TR-albumin and once with PBS. Finally, ECs were lysed with 1% Triton X-100 in Milli-Q water and the fluorescent signal of internalized TR-albumin was quantified with a Fluorolog 3 spectrofluorometer (Horiba Jobin Yvon) at 592 nm excitation and 612 nm emission wavelengths. Concentrations of internalized TR-albumin were calculated using a standard calibration curve. Values were normalized for total protein content in each well using a Pierce BCA Protein Assay Kit (Thermo Fisher, cat# 23225) according to the manufacturer’s instructions.

### Measurement of barrier integrity in response to inflammation

For the measurement of barrier integrity by impedance, human stem cell-derived ECs were cultured in the xCELLigence RTCA SP (Agilent) device and treated with cARLA or control medium exactly as described above. After 48 h of cARLA or control treatment, ECs were challenged with a combination of TNF-α (PeproTech, cat# 300-01A) and IL-1β (PeproTech, cat# 200-01B) in multiple concentrations (0.1-10 ng/mL). The impedance of monolayers was measured at 10 kHz at every 5 min for 48 h. For the measurement of TEER and tracer permeability, the human brain-like EC-PC co-culture model was assembled, maintained and treated with cARLA or control medium as described above. After 48 h of cARLA or control treatment, the luminal (‘blood side’) compartment of the model was treated with a combination of TNF-α (10 ng/mL) and IL-1β (10 ng/mL) for 24 h and TEER and permeability was assessed exactly as described above.

### Pharmacological inhibition of Wnt/β-catenin signaling

Wnt/β-catenin signaling was blocked at two different points using inhibitors ICG-001 (MedChemExpress, cat# HY-14428) and XAV 939 (MedChemExpress, cat# HY-15147). For the measurement of barrier integrity by impedance, human brain-like ECs were cultured as described above. Confluent monolayers were treated with ICG-001 (5-50 µM) or XAV 939 (0.5-3 µM) for 48 h at 37 °C, and impedance was measured at 10 kHz every 5 min using the xCELLigence RTCA SP device (Agilent) as described above. To investigate how the effect of cARLA is modulated by these inhibitors, ECs were co-treated with cARLA and either 5 µM ICG-001 or 1 µM XAV 939 for 48 h at 37 °C. For the measurement of TEER and tracer permeability, EC-PC co-cultures were assembled and maintained as described above. On day 4, the luminal (‘blood side’) compartment of the model was incubated with cARLA or control medium supplemented with either 5 µM ICG-001 or 1 µM XAV 939 for 48 h at 37 °C. TEER and tracer permebility were then measured and calculated as described above.

### Penetration of small molecule drugs across the human brain-like EC-PC BBB model

The penetration of ten selected small molecule drugs (atenolol, indomethacin, lamotrigine, loperamide, methotrexate, propranolol, salicylic acid, tacrine, verapamil and vinblastine) was measured in blood-to-brain and brain-to-blood directions across the human brain-like EC-PC co-culture model. Properties and catalog numbers of drugs are listed in **SI Appendix, Table S3**. The assembly, maintainance and treatment of co-cultures were performed as described above. After 48 h cARLA or control treatment, either the luminal (blood) or abluminal (brain) compartment of the model was incubated with 1 µM of drug compounds (stocks of 10 mM were prepared in DMSO) diluted in Ringer-HEPES solution (118 mM NaCl, 4.8 mM KCl, 2.5 mM CaCl_2_, 1.2 mM MgSO_4_, 5.5 mM D-glucose, 20 mM HEPES, pH 7.4) supplemented with 0.1% FBS (Sigma-Aldrich, cat# F4135) at 37 °C on a horizontal shaker set to 150 rpm. In brain-to-blood direction, samples from both compartments were collected at 60 min. In blood-to-brain direction, inserts were transferred to a new well with fresh buffer at 30 min, then at the end of the assay samples were collected from the donor (0-60 min) and acceptor compartments (0-30 min, 30-60 min). Samples were precipitated with –20 °C methanol (Molar Chemicals, cat# 05730-101-340) and drug content was measured using a Sciex API 4000 MS-Agilent 1260 Infinity UHPLC instrumental setup, except for methotrexate, which was measured using a Sciex x500r QTOF MS – Exion LC setup. Measurement parameteres are listed in **SI Appendix, Table S4**. Drug concentrations were determined by standard calibration curves. To calculate apparent permeability coefficients (P_app_) using multiple timepoints, clearance was first calculated^57,66^:

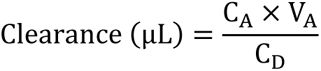

where C_A_ is the concentration measured in the acceptor compartment, V_A_ is the volume of the acceptor compartment (1.5 mL) and C_D_ is the treatment concentration in the donor compartment at a given timepoint. Cumulative clearance values were plotted against time at 0, 30 and 60 min and a simple linear regression was used to calculate clearance rate (PS_t_; µL/min) as the slope of the curve. P_app_ was then calculated as:

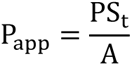

where A is the surface area of the insert available for permeability (1.12 cm^2^). Efflux ratio was calculated by dividing P_app_ values in brain-to-blood direction by P_app_ values in blood-to-brain direction. To calculate the *in vitro* unbound brain-to-plasma partition coefficients (K_p,uu,brain_)^22,25,55^ of drugs in the blood-to-brain direction, recovery for each compound was first calculated as:

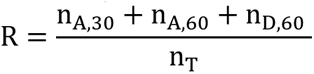

where n_A,30_, n_A,60_ and n_D,60_ are the amounts of substance (pmol) in the acceptor compartment at 30– and 60 min as well as the donor compartment at 60 min, respectively, and n_T_ is the amount of substance (pmol) present at the treatment solution at 0 min. *In vitro* K_p,uu,brain_ was then calculated as:

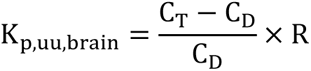

where C_T_ is the concentration of a drug in the treatment solution at 0 min, C_D_ is the concentration in the donor compartment at 60 min and R is recovery. Calculation of Pearson *r*^2^ and Spearman *r* coefficients was performed in GraphPad Prism software (version 5.0).

### Synthesis and penetration of nanoparticles across the human brain-like EC-PC BBB model

Poly(L-glutamic acid) nanoparticles (NPs) were synthesized as previously described^70^. As a targeting ligand for penetration across the BBB, reduced L-glutathione was added to NPs *via* the EDC/NHS coupling reaction in a molar ratio of L-glutamic acid:L-glutathione=2:1. NPs were grafted with *N*-(2-aminoethyl) rhodamine 6G-amide bis(trifluoroacetate) (rhodamine 6G) *via* EDC/NHS coupling in a weight ratio of NP:rhodamine 6G=1:20 to fluorescently label the particles. The assembly, maintainance and treatment of BBB co-cultures were performed as described above. After 48 h cARLA or control treatment, the luminal (‘blood side’) compartment of the model was incubated with non-targeted or GSH-targeted NPs (50 µg/mL) diluted in phenol red-free DMEM/F-12 medium (Gibco, cat# 21041025) supplemented with 1% FBS (Sigma-Aldrich, cat# F4135) at 37 °C on a horizontal shaker set to 150 rpm. Inserts were transferred to a new well with fresh buffer at 1 h and 4h, then at the end of the assay samples were collected from the donor (0-24 h) and acceptor compartments (0-1 h, 1-4 h, 4-24 h). The mean fluorescence of rhodamine 6G was measured using a Fluorolog 3 spectrofluorometer (Horiba Jobin Yvon) with excitation/emission wavelengths 525/-551 nm. NP concentrations were determined by standard calibration curves. To calculate endothelial permeability coefficients (P_e_) using multiple timepoints^57^, clearance and clearance rate were first calculated as described above for inserts with cells and for blank inserts without cells (PS_total_ and PS_blank_). The permeability × surface area product value for the endothelial monolayer (PS_e_) was calculated using the following formula:

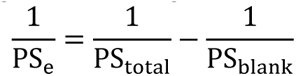

P_e_ was then calculated by dividing permeability × surface area product value for the endothelial monolayer by the surface area of the insert available for permeability (1.12 cm^2^):

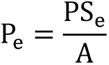

For the visualization of NP internalization, human brain-like ECs were grown in 50% PC-conditioned medium on 35 mm glass bottom Petri dishes (Greiner Bio-One, cat# 627960) coated with collagen IV (100 µg/mL) and fibronectin (25 µg/mL) at a seeding density of 5×10^4^ cells/cm^2^ until confluence. After 48 h treatment with cARLA or control medium, cells were incubated with non-targeted or GSH-targeted NPs (50 µg/mL) diluted in EC medium for 24 h at 37 °C. For the final 20 minutes of incubation, Hoechst 33342 (Thermo Fisher, cat# H1399, 2 μg/mL) was added to cells to counterstain nuclei. Cultures were washed twice in Ringer-HEPES supplemented with 1% FBS and live-cell imaging was performed immediately in the same buffer using a Leica TCS SP5 AOBS confocal laser scanning microscope equipped with a HCX PL APO 63× oil (NA=1.4) objective exactly as described above.

### Statistics

Individual wells or inserts of cultured cells that underwent the same experimental procedure are defined as replicates. For immunocytochemistry, individual fields of view (4-5 fields of view per culture insert) are defined as replicates. All key experiments were repeated independently and detailed information about error bars, sample size, replication strategy and statistical tests are provided in each figure legend. Statistical analyses were performed in GraphPad Prism software (version 5.0). Unpaired *t*-test or one-way ANOVA followed by Bonferroni’s post-hoc test were used to compare means from two or more groups, respectively, on normally distributed data determined by Shapiro-Wilk test. In experiments with two independent variables, two-way ANOVA with Bonferroni’s post-hoc test was used. Statistical significance was set at *P* < 0.05, and FDR < 0.01 for MACE-seq.

## Data availability

MACE-seq datasets generated and analyzed in this study have been deposited to the Gene Expression Omnibus (GEO) repository under the accession number GSE224846.

## Acknowledgments

Prof. Christer Betsholtz and Dr. Maarja A. Mäe from Uppsala University Sweden are gratefully acknowleged for providing the *Cldn5*^+/-^ mice. We are grateful for Dr. Hideaki Nishihara and Dr. Kinya Matsuo from Yamaguchi University for their advice and guidance on iPSC-derived EECM-BMEC-like cells. We would also like to thank the technical assistance of Katalin Kokavszky and Csilla Kovács from the HUN-REN BRC in the experiments. We are grateful for Dr. Imola Wilhelm and Dr. István Krizbai from the HUN-REN BRC for the use of a CellZScope+ instrument and Dr. Tibor Páli from the HUN-REN BRC for the use of a Horiba Jobin-Yvon Fluorolog 3 spectofluorometer. We also thank Ildikó Valkonyné Kelemen and Dr. Gábor Steinbach from the Cellular Imaging Laboratory at the HUN-REN BRC for providing support in microscopy.

This work was funded by the National Research, Development and Innovation Office of Hungary (K143766), and the Hungarian Academy of Sciences (NAP2022-I-6/2022) for M.A.D. G.P. was supported by the National Academy of Scientist Education Program of the National Biomedical Foundation under the sponsorship of the Hungarian Ministry of Culture and Innovation, the Stephen W. Kuffler Research Foundation, the Gedeon Richter Plc. Centenarial Foundation, and by the New National Excellence Program of the Ministry for Innovation and Technology (ÚNKP-22-3-SZTE-446). M.M. was supported by the the National Research, Development and Innovation Office, Budapest, Hungary (PD138930), the Gedeon Richter Plc. Centenarial Foundation and the “National Talent Program” with the financial aid of the Ministry of Human Resources (NTP-NFTÖ-22-B-0150). A.S. was supported by the New National Excellence Program of the Ministry for Innovation and Technology (ÚNKP-22-3-SZTE-458), and the Gedeon Richter Plc Centenarial Foundation (2022/R/26/2533). J.P.V. was supported by the „National Talent Program” with the financial aid of the Ministry of Human Resources (NTP-NFTÖ-22-B-0229). F.R.W. was supported by the Secretariat of Lorand Eotvos Research Network (SA-111/2021). S.V. was supported by the Premium Postdoctoral Research Program (Premium-2019–469) of the Hungarian Academy of Sciences and by National Research, Development and Innovation Office of Hungary (FK143233). The Campbell lab is supported by grants from SFI (Eye-D-21/SPP/3732), Enterprise Ireland, The Irish Research Council (IRC) and by a research grant from SFI under grant number 16/RC/3948 and co-funded under the European Regional Development fund by FutureNeuro industry partners. The lab is also supported by a European Research Council (ERC) grant, “Retina-Rhythm” (864522).

