## Supplementary information for "Synergistic induction of blood-brain barrier properties"

**SI Appendix includes:**

Supplementary Figures S1 to S18
Supplementary Tables S1 to S4

**Table of contents:**

**Figure S1.** Effect of small molecules and recombinant proteins targeting single pathways in human stem cell-derived ECs on barrier integrity measured by impedance.

**Figure S2.** Effect of simultaneous activation of Wnt and inhibition of TGF-β signaling on barrier integrity in human stem cell-derived ECs.

**Figure S3.** Effect of simultaneous activation of cAMP and Wnt as well as inhibition of TGF-β signaling on barrier integrity in human stem cell-derived ECs.

**Figure S4.** Additional measurements of barrier- and junctional integrity in brain-like ECs.

**Figure S5.** Testing the effect of cARLA on barrier tightness with or without pericyte influence.

**Figure S6.** The effect of cARLA on barrier tightness is reproducible in a human induced pluripotent stem cell-derived BBB model.

**Figure S7.** Optimizing concentrations for barrier integrity in mouse bEnd.3 cells.

**Figure S8.** Optimizing concentrations for barrier integrity in in rat primary brain ECs.

**Figure S9.** The statistically most significant as well as BBB-specific gene expressional changes induced by cARLA.

**Figure S10.** The effect of cARLA on vimentin and Ki-67 expression.

**Figure S11.** The effect of cARLA on barrier maturation and tight junction expression.

**Figure S12.** Correlating the effect of cARLA with in vivo data from mice and humans.

**Figure S13.** Changes in gene expression related to different aspects of BBB function in brain-like ECs.

**Figure S14.** The effect of cARLA and EC-PC co-culture on WGA lectin staining and VCAM-1 levels.

**Figure S15.** Further validation of the effect of cARLA on efflux pump expression, localization and activity.

**Figure S16.** The effect of cARLA and its constituents on Wnt signaling.

**Figure S17.** Additional experiments on Wnt/β-catenin signaling in brain-like ECs.

**Figure S18.** Comparison of $K_{p,uu,brain}$ values for selected drugs across species and BBB models.

**Table S1.** The list of 100 BBB-specific genes with robust expression at the human brain endothelium.

**Table S2.** The list of BBB-specific genes in mice as reported by previous studies.

**Table S3.** Properties of small molecule drugs tested in this study.

**Table S4.** Parameters for measuring the passage of small molecule drugs across the BBB co-culture model.

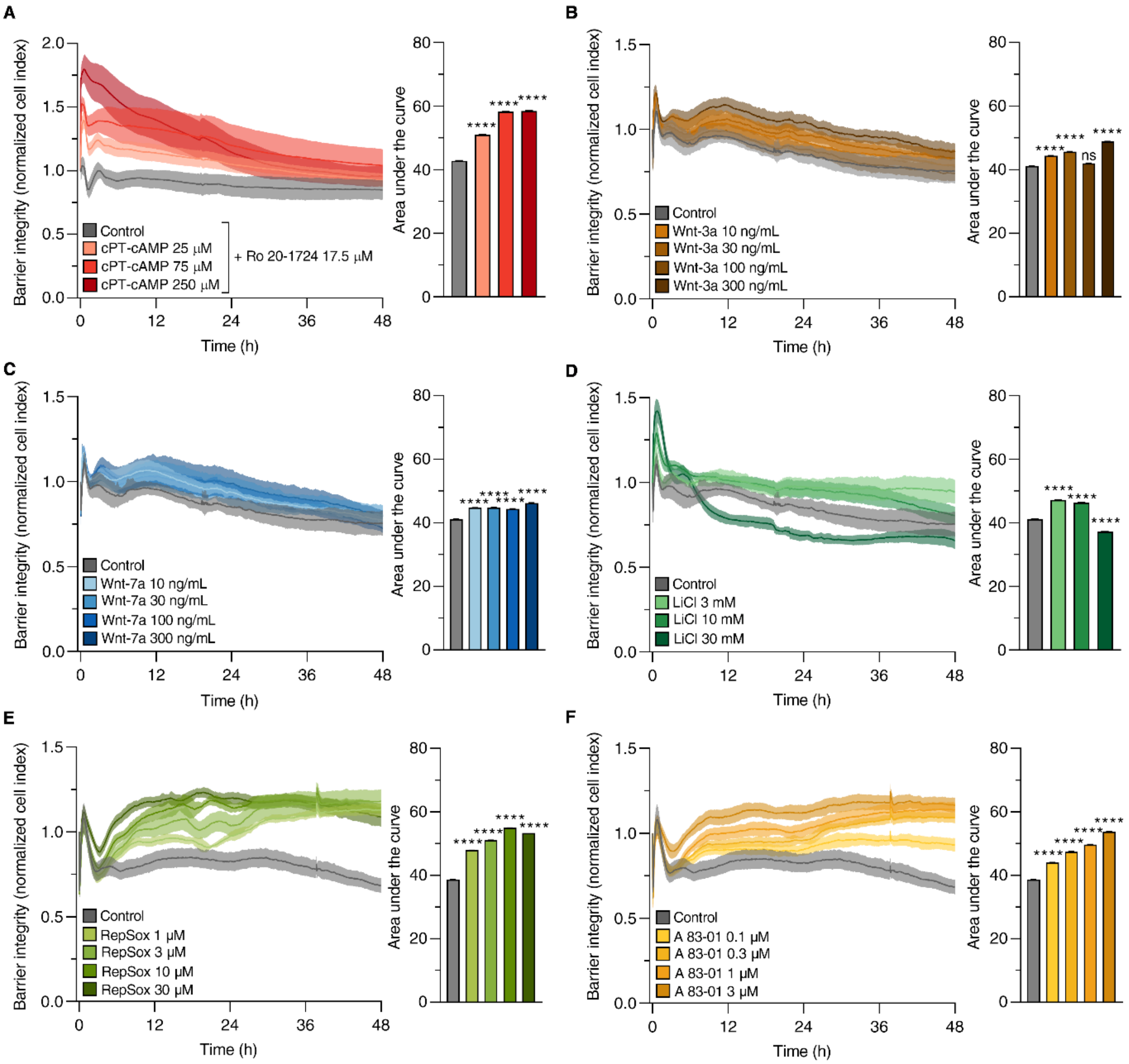


**Figure S1. Effect of small molecules and recombinant proteins targeting single pathways in human stem cell-derived ECs on barrier integrity measured by impedance.** Treatment concentrations and durations were optimized for **A)** pCPT-cAMP supplemented with the cAMP-specific phosphodiesterase inhibitor Ro 20-1724, **B-D)** activators of Wnt signaling and **E-F)** inhibitors of TGF-β signaling. Higher normalized cell index values and a higher area under the curve indicates increased barrier integrity. Mean ± SD, ANOVA with Bonferroni's post-hoc test, ****$P<0.0001$, ns: $P>0.05$ compared to the control group, $n=6$.

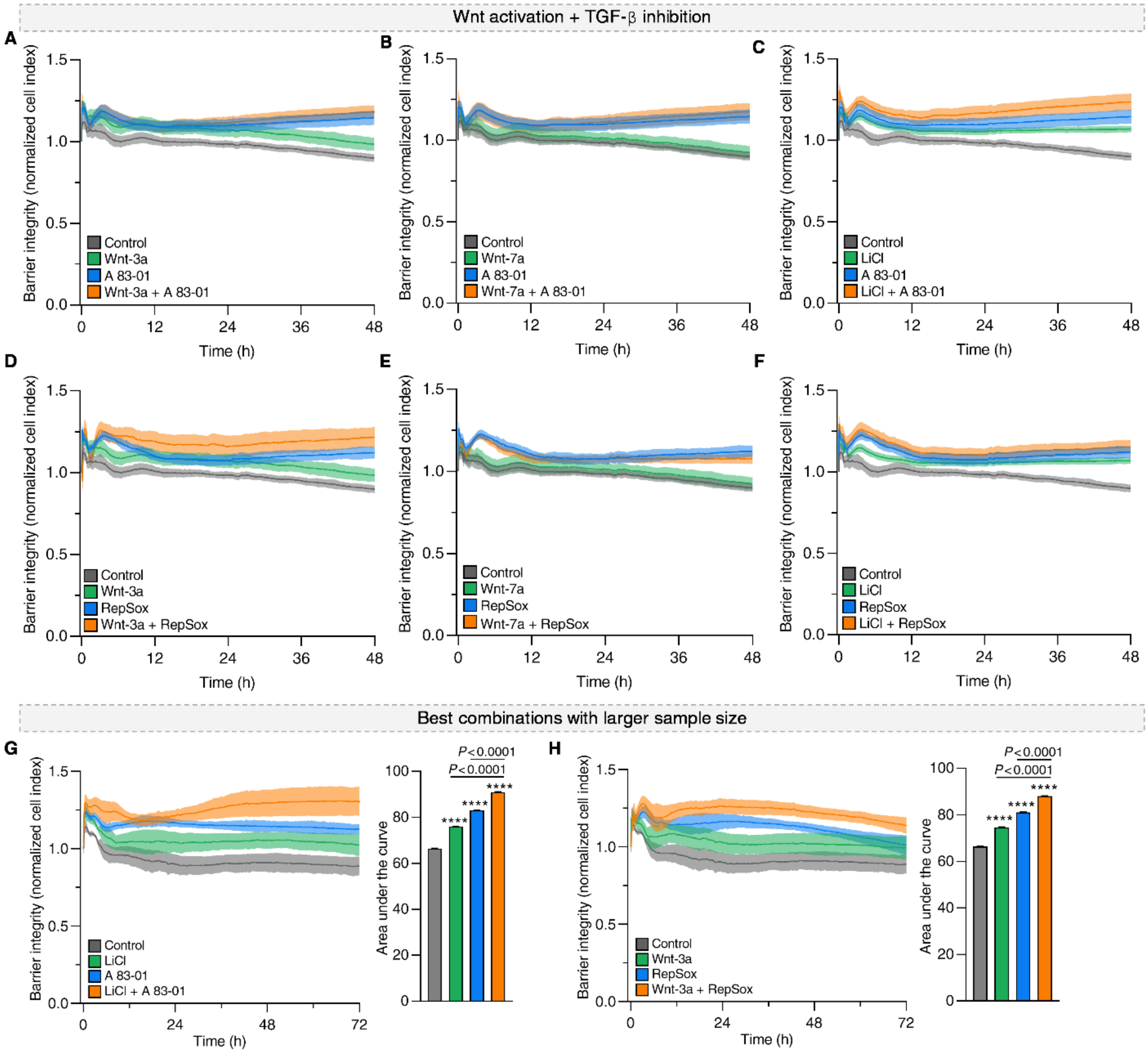


**Figure S2. Effect of simultaneous activation of Wnt and inhibition of TGF-β signaling on barrier integrity in human stem cell-derived ECs.** Impedance kinetics of ECs treated with **A-F)** Wnt activators (Wnt-3a 300 ng/mL, Wnt-7a 300 ng/mL, LiCl 3 mM), TGF-β receptor antagonists (RepSox 10 μM, A83-01 3 μM) and their combinations. Mean ± SD, *n*=5. **G-H)** The best combinations (LiCl+A83-01, Wnt-3a+RepSox) were also validated with a larger sample size. Higher normalized cell index values and a higher area under the curve indicates increased barrier integrity. Mean ± SD, ANOVA with Bonferroni's post-hoc test, ****$P$<0.0001 compared to the control group, *n*=10.

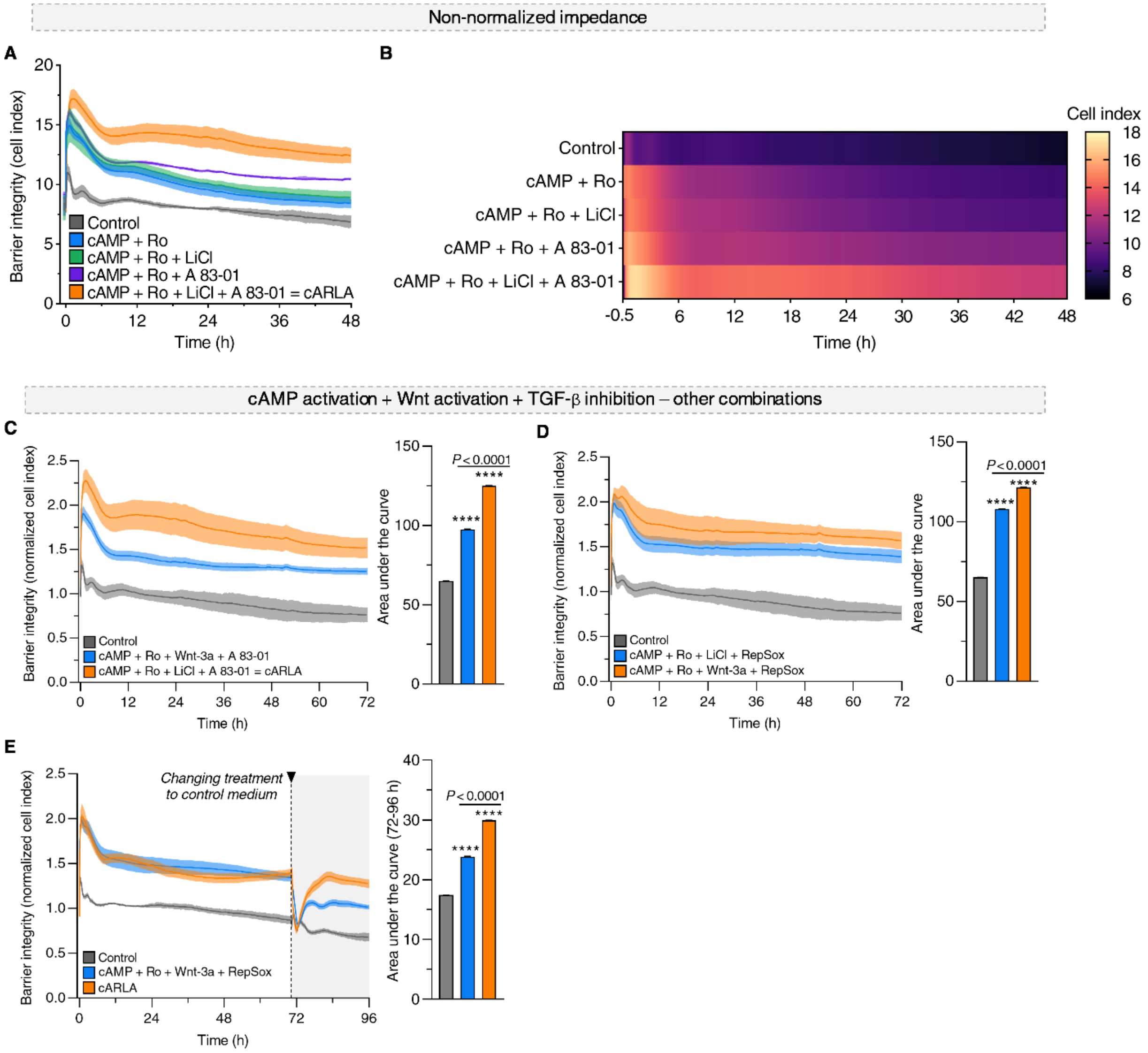


**Figure S3. Effect of simultaneous activation of cAMP and Wnt as well as inhibition of TGF-β signaling on barrier integrity in human stem cell-derived ECs. A)** Non-normalized data (raw cell index values) of data presented in Fig. 1C. **B)** The same non-normalized data presented as a heat map. **C)** Impedance kinetics of ECs treated with combinations containing A83-01 and **D)** RepSox, targeting all three pathways simultaneously. The same concentrations of compounds were used as described above. Of note, some combinations (cARLA, cAMP+Ro+Wnt-3a+RepSox) more effectively elevated barrier integrity than others (cAMP+Ro+Wnt-3a+A83-01, cAMP+Ro+LiCl+RepSox). **E)** Impedance kinetics of EC monolayers treated with the two best combinations. Treatment was changed to control medium at 72 h. Of note, cARLA outperformed cAMP+Ro+Wnt-3a+RepSox, since its effect was more resilient to the withdrawal of treatment. This makes cARLA a more suitable candidate to be used in permeability experiments, where the presence of small molecule compounds can potentially interfere with downstream analytics. Higher normalized cell index values and a higher area under the curve indicates increased barrier integrity. Mean ± SD, ANOVA with Bonferroni's post-hoc test, ****$P$<0.0001 compared to the control group, $n$=6 for all three panels.

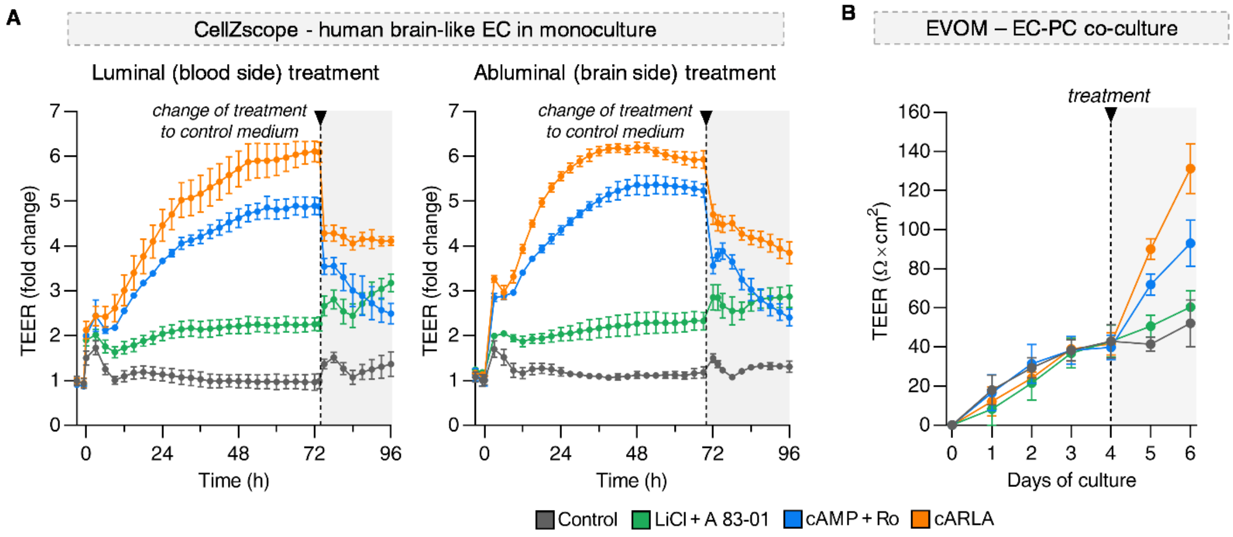


**Figure S4. Additional measurements of barrier- and junctional integrity in brain-like ECs. A)** Automated measurement of transendothelial electrical resistance (TEER) in brain-like ECs after treatment with cARLA or its components using the cellZscope instrument. Luminal and abluminal treatment was only applied for compounds LiCl and A83-01. Compounds cAMP+Ro were always administered luminally. **B)** Manual measurement of TEER across the EC-PC co-culture model over time using an EVOM voltohmmeter and hand-held STX-2 electrodes.

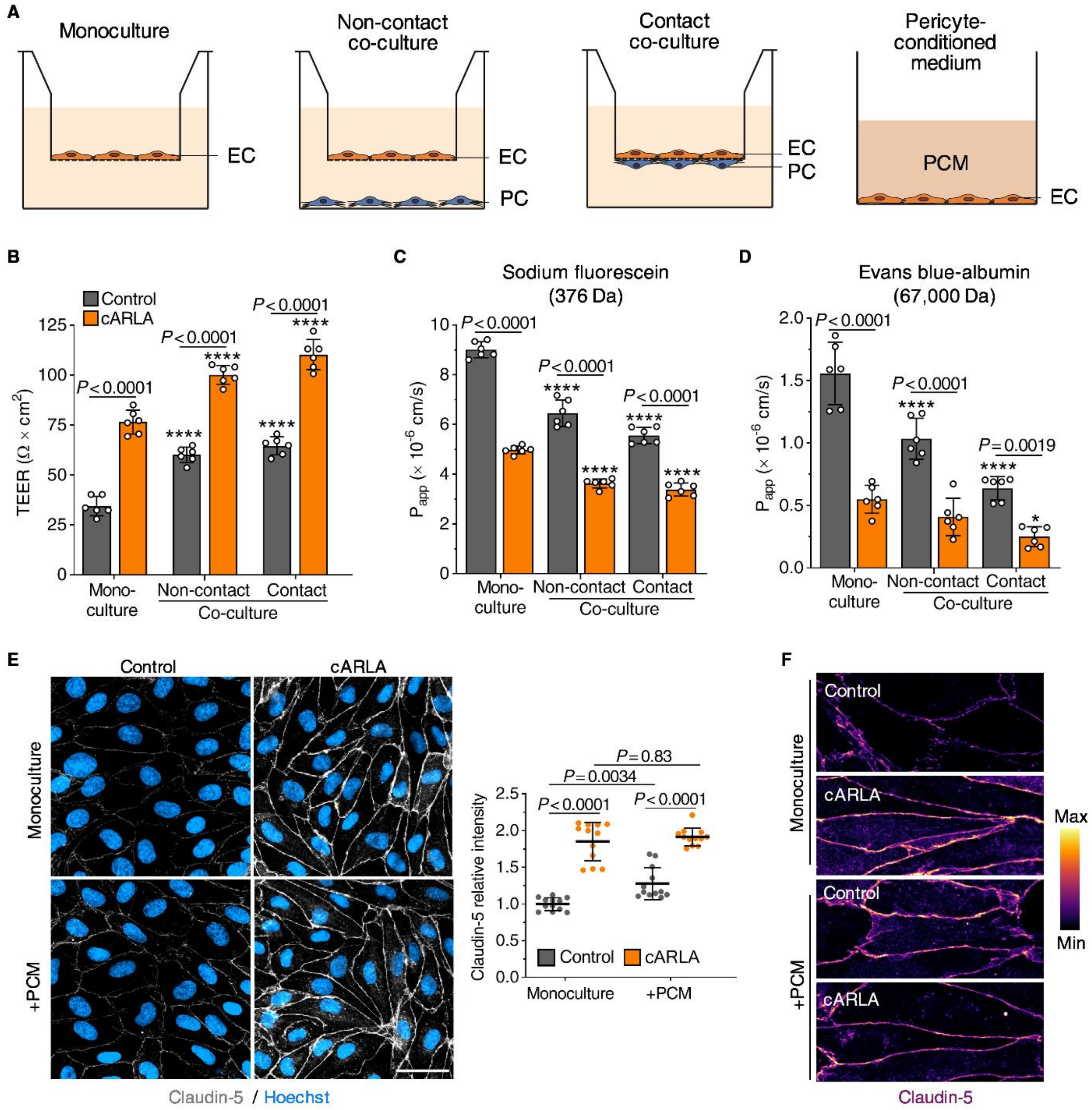


**Figure S5. Testing the effect of cARLA on barrier tightness with or without pericyte influence. A)** Schematic drawing of the types of culture conditions used. EC: endothelial cell, PC: pericyte, PCM: pericyte-conditioned medium. **B)** Transendothelial electrical resistance (TEER) after 48 h cARLA treatment. Mean ± SD, ANOVA with Bonferroni's post-hoc test, ****$P < 0.0001$ compared to its respective monoculture group, $n$=6. **C)** Permeability of sodium fluorescein and **D)** Evans blue-albumin across the co-culture model after 48 h cARLA treatment. $P_{app}$: apparent permeability coefficient. Mean ± SD, ANOVA with Bonferroni's post-hoc test, ****$P < 0.0001$ compared to its respective monoculture group, $n$=6. **E-F)** Claudin-5 immunostaining in control and cARLA-treated EC monolayers, with or without the addition of PCM. Bar: 50 µm. Quantification: Mean ± SD, ANOVA with Bonferroni's post-hoc test, $n$=12.

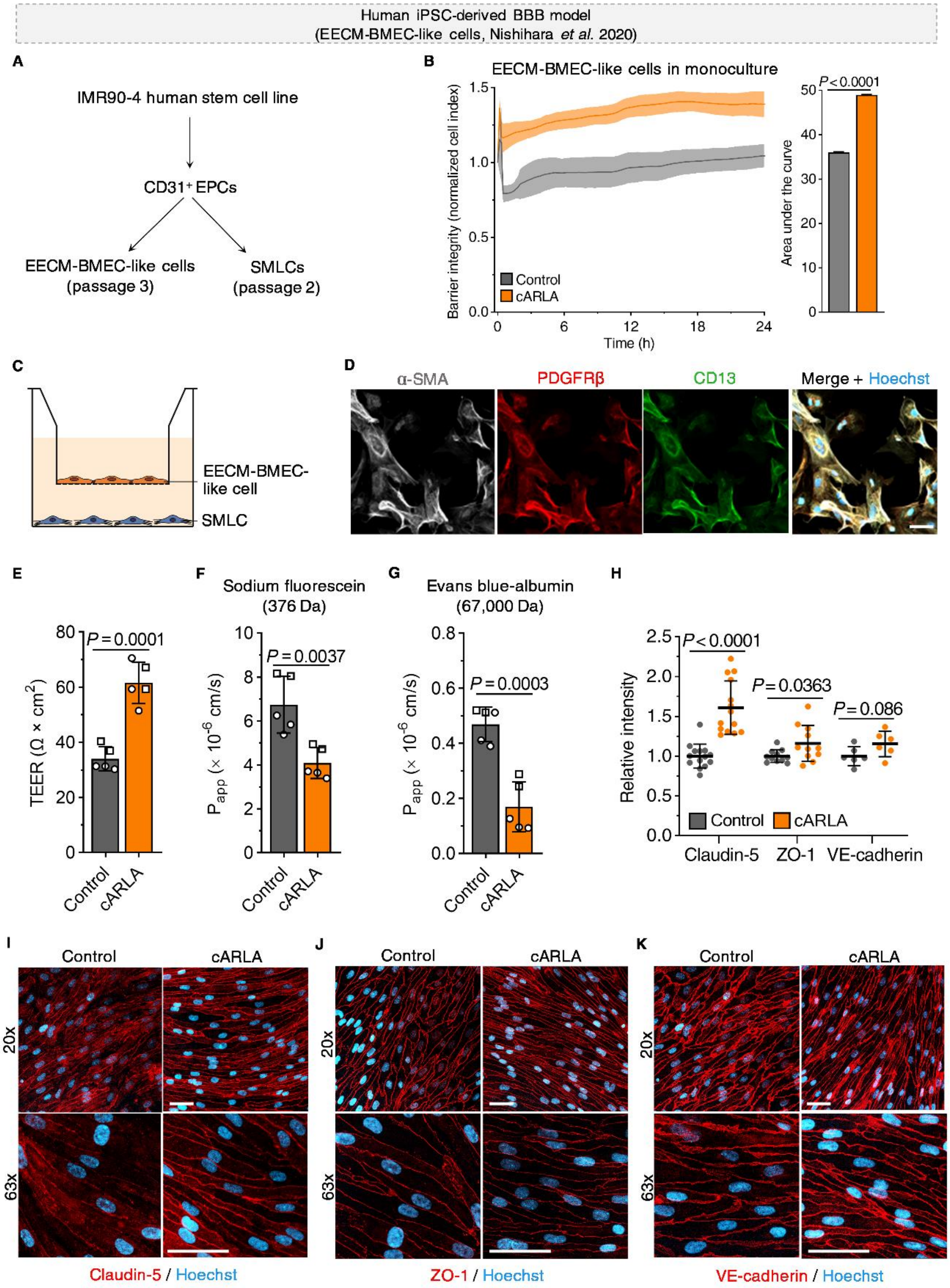


**Figure S6. The effect of cARLA on barrier tightness is reproducible in a human induced pluripotent stem cell-derived BBB model. A)** Overview of the differentiation protocol[21]. EPC: endothelial progenitor cell; EECM-BMEC: extended endothelial culture method - brain microvascular endothelial cell; SMLC: smooth muscle-like cell. **B)** Barrier integrity of EECM-BMEC-like cell monolayers measured by impedance in real time. Higher normalized cell index values and a higher area under the curve indicates increased barrier integrity. Mean ± SD, unpaired t-test, *n*=3. **C)** Schematic drawing of the co-culture model[21]. **D)** Validation of SMLC identity by immunostaining. α-SMA: alpha smooth muscle actin; PDGFRβ: platelet-derived growth factor receptor beta. Bar: 50 μm. **E)** Transendothelial electrical resistance (TEER) in the co-culture model after 48 h cARLA treatment. Mean ± SD, unpaired t-test, *n*=5 from 2 independent differentiations. **F)** Permeability of sodium fluorescein and **G)** Evans blue-albumin across the co-culture model after 48 h treatment. $P_{app}$: apparent permeability coefficient. Mean ± SD, unpaired t-test, *n*=5 from 2 independent differentiations. **H)** Intensity-based quantification of **I)** claudin-5, **J)** ZO-1, and **K)** VE-cadherin immunostainings in the co-culture model. Raw values were normalized to Hoechst intensity in each image. Mean ± SD, multiple unpaired t-tests, *n*=6-12. Bar: 50 μm.

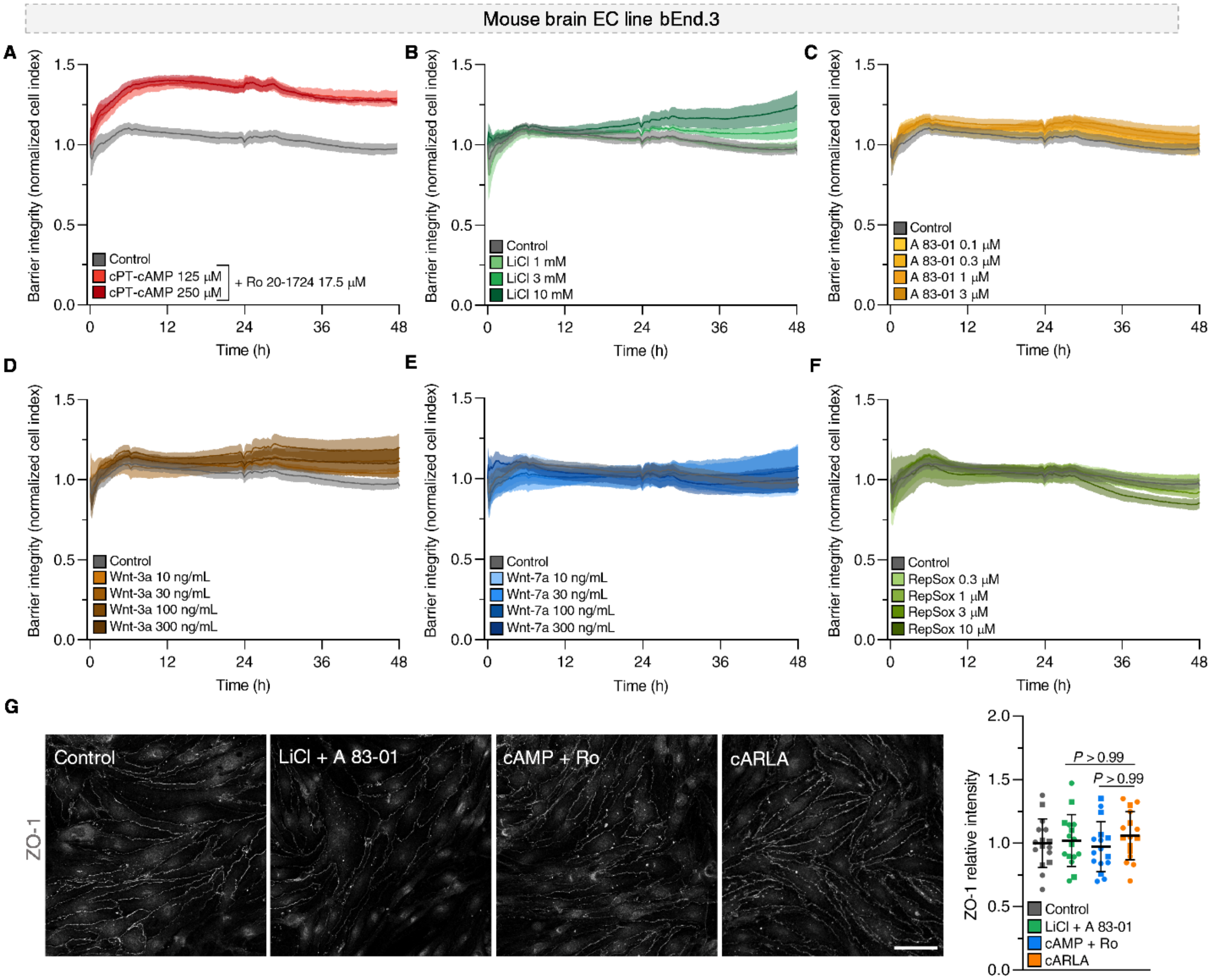


**Figure S7. Optimizing concentrations for barrier integrity in mouse bEnd.3 cells. A-F)** The effect of small molecules and recombinant proteins targeting single pathways on barrier integrity, measured by impedance. Higher normalized cell index values and a higher area under the curve indicates increased barrier integrity. Mean ± SD, *n*=5-6. **G)** Immunostaining of the tight junction protein ZO-1 in bEnd.3 cells after 48 h treatment. Bar: 50 µm. Quantification: Mean ± SD, ANOVA with Bonferroni's post-hoc test, *n*=16 from 2 independent experiments.

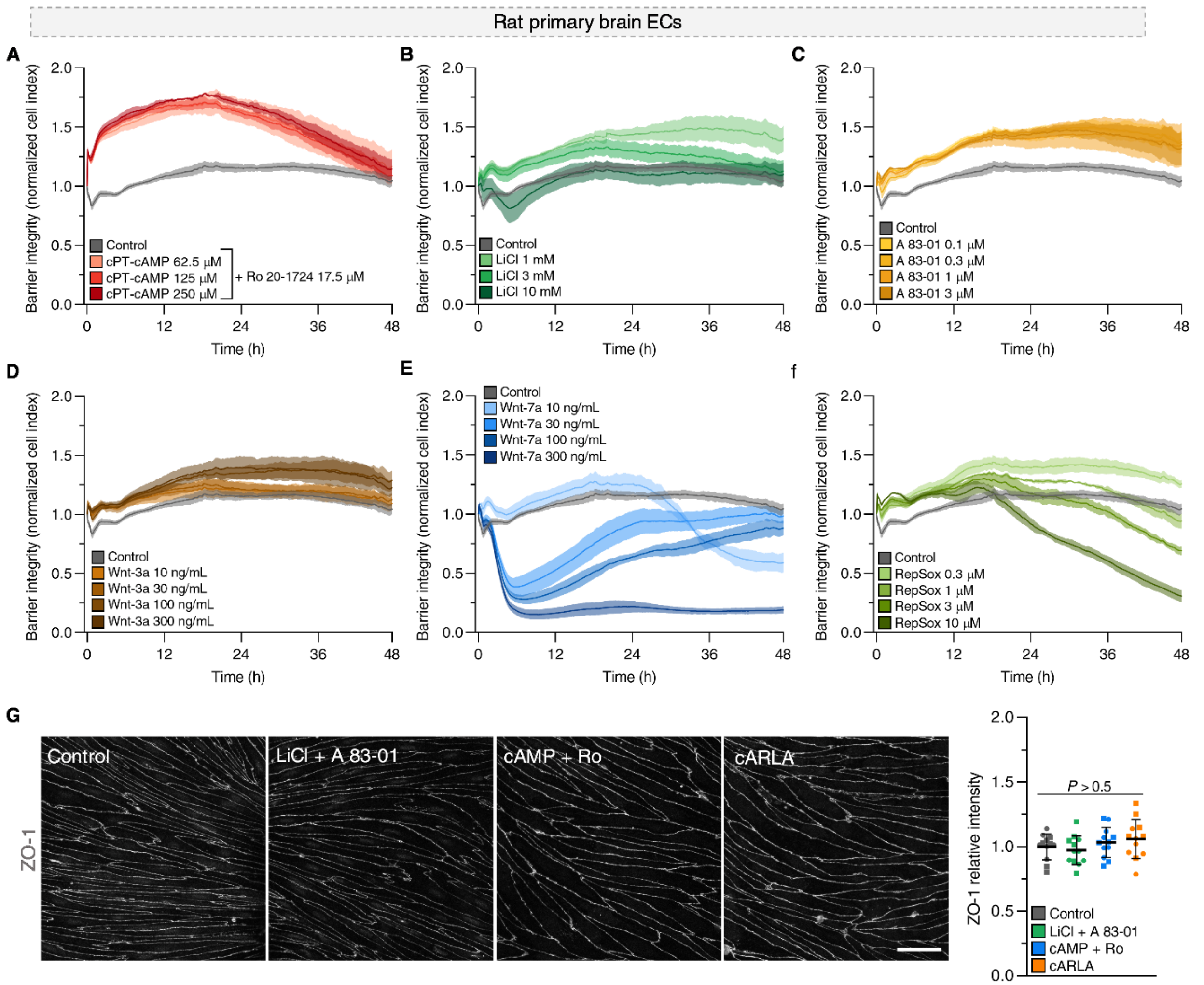


**Figure S8. Optimizing concentrations for barrier integrity in in rat primary brain ECs**. **A-F)** The effect of small molecules and recombinant proteins targeting single pathways on barrier integrity, measured by impedance. Higher normalized cell index values and a higher area under the curve indicates increased barrier integrity. Mean ± SD, *n*=5-6. **G)** Immunostaining of the tight junction protein ZO-1 in rat primary brain ECs from EC-PC-AC co-cultures after 48 h treatment. Bar: 50 µm. Quantification: Mean ± SD, ANOVA with Bonferroni's post-hoc test, *n*=12 from 2 independent experiments.

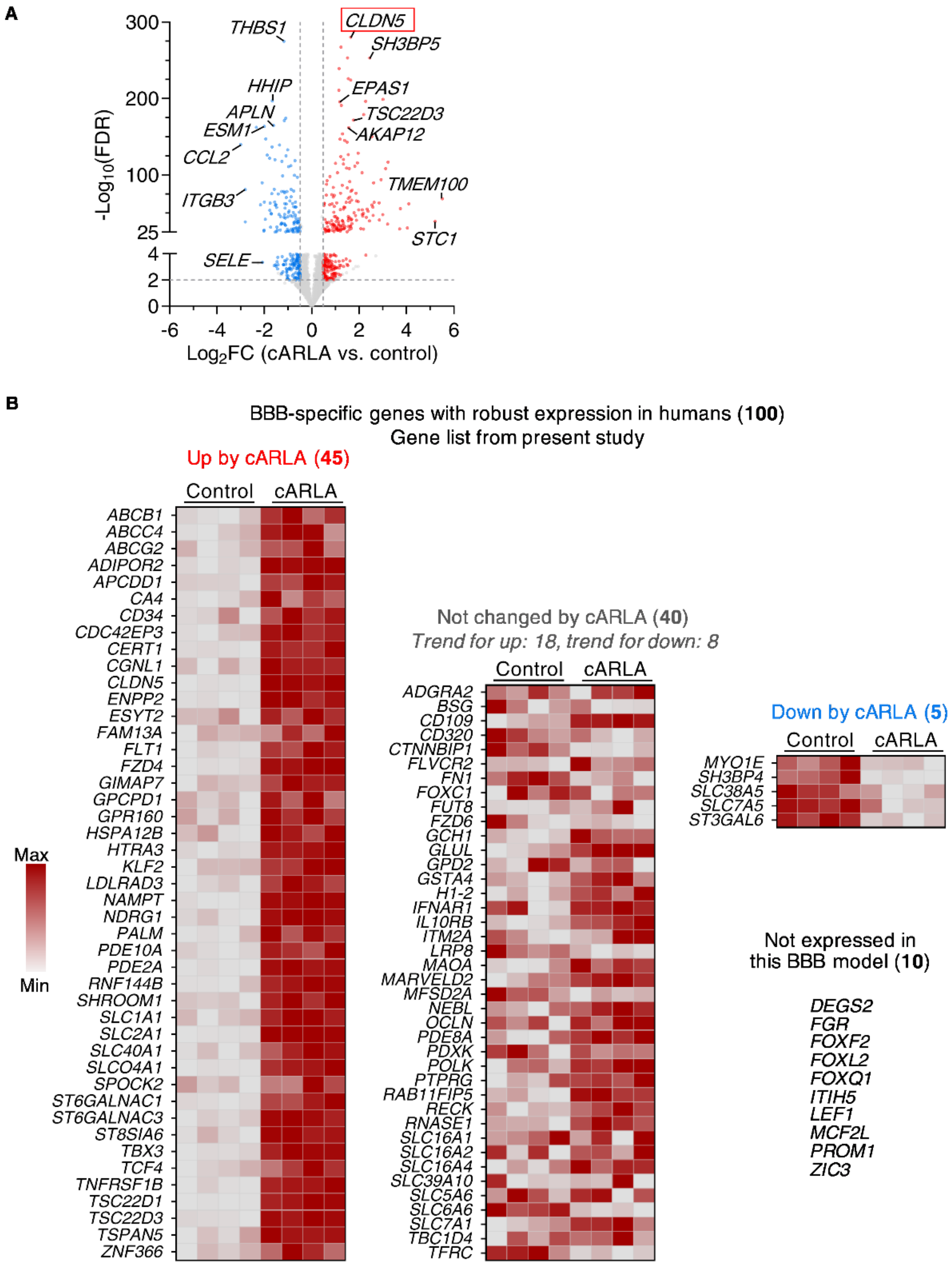


**Figure S9. The statistically most significant as well as BBB-specific gene expressional changes induced by cARLA. A)** Volcano plot of the statistically most significant up- and downregulated genes by cARLA. FDR: adjusted *P*-value (Benjamini-Hochberg method). $Log_2FC$: $log_2$(fold change). *CLDN5* is highlighted as having the lowest FDR value. **B)** Scaled heat map of our list of 100 BBB-specific genes identified by previous studies in mice[4,50,53] that are also robustly expressed at the human BBB *in vivo*[54,55,69,70]. Genes that are upregulated, not changed and downregulated by cARLA as well as genes that are not expressed in the human brain-like EC-PC co-culture model are shown in separate columns, in alphabetical order.

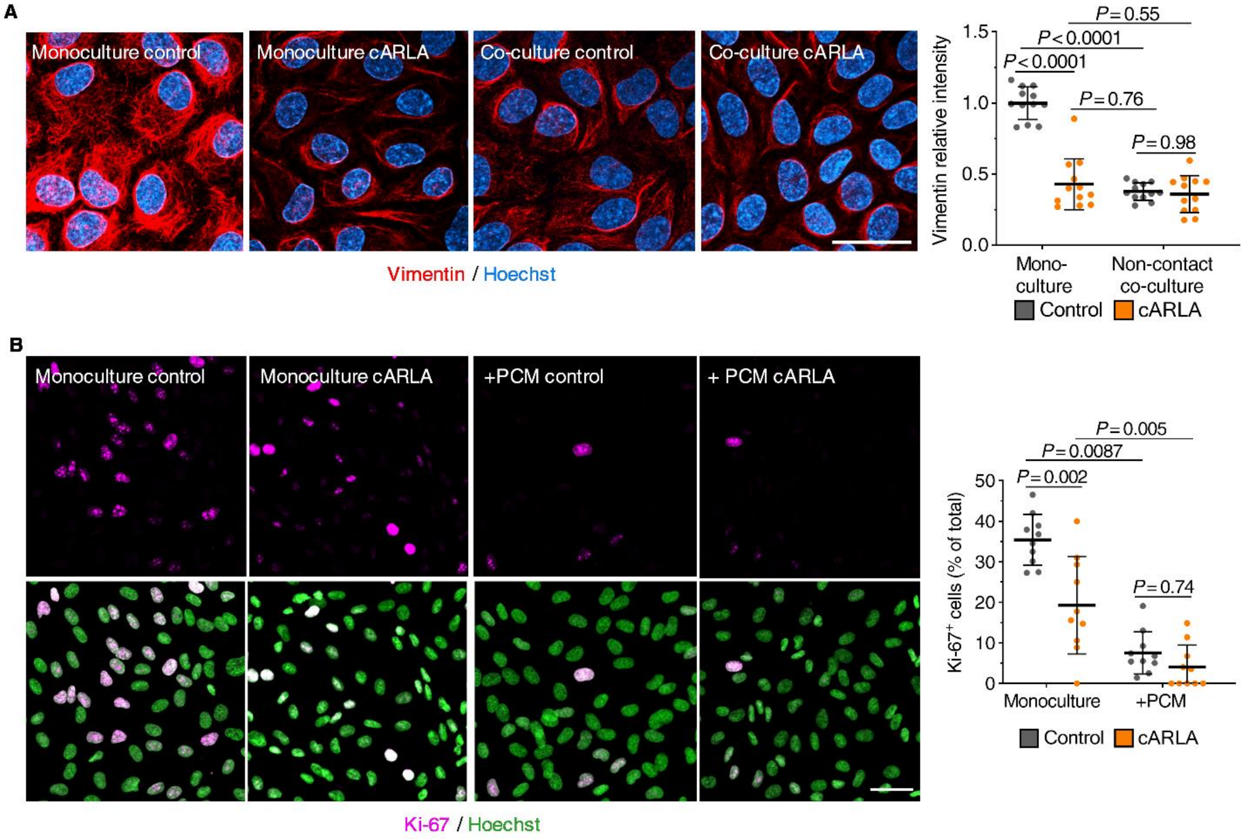


**Figure S10. The effect of cARLA on vimentin and Ki-67 expression. A)** Vimentin immunostaining in human ECs grown in monoculture or in non-contact co-culture with PCs. Quantification**:** Mean ± SD, ANOVA with Bonferroni's post-hoc test, *n*=12. **B)** Ki-67 immunostaining in human ECs grown in monoculture, with or without PCM. Quantification**:** Mean ± SD, ANOVA with Bonferroni's post-hoc test, *n*=10. Bar: 50 µm.

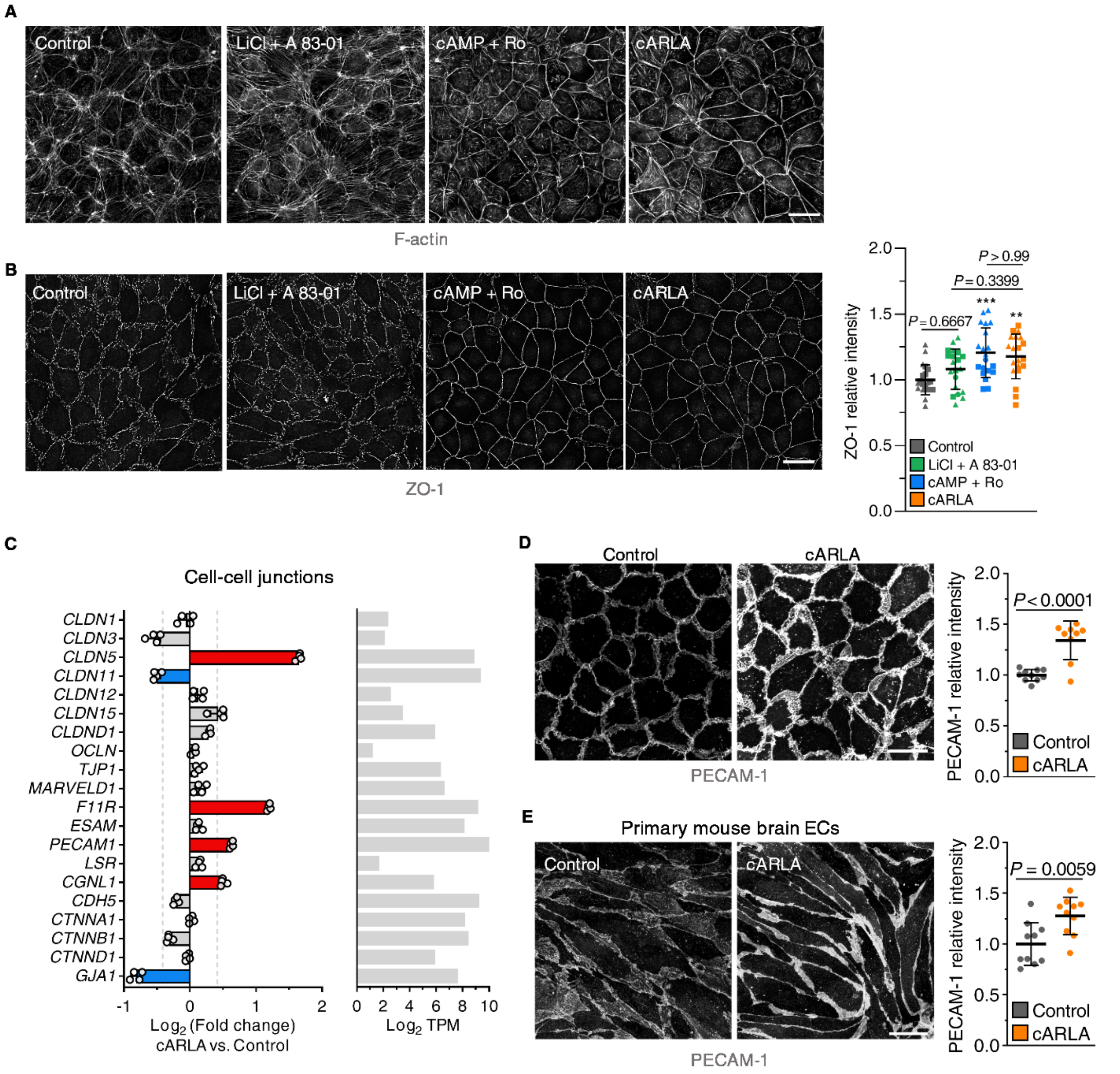


**Figure S11. The effect of cARLA on barrier maturation and tight junction expression. A)** F-actin and **B)** ZO-1 immunostaining in human stem cell-derived EC-PC co-cultures. Bar: 50 µm. Quantification: Mean ± SD, ANOVA with Bonferroni's post-hoc test, **$P$<0.01, ***$P$<0.001 compared to the control group, $n$=20-21 from 2 independent experiments. **C)** Changes in gene expression related to cell-cell junctions in human brain-like ECs. Mean ± SD, $n$=4. Red and blue color indicates up- and downregulation, respectively, upon cARLA treatment. Grey color indicates no differential expression. TPM: transcript per million. **D)** PECAM-1 (CD31) immunostaining in human brain-like ECs and **E)** in primary mouse brain ECs from EC-PC-AC co-culture. Bar: 50 µm. Quantification: Mean ± SD, unpaired t-test, $n$=10.

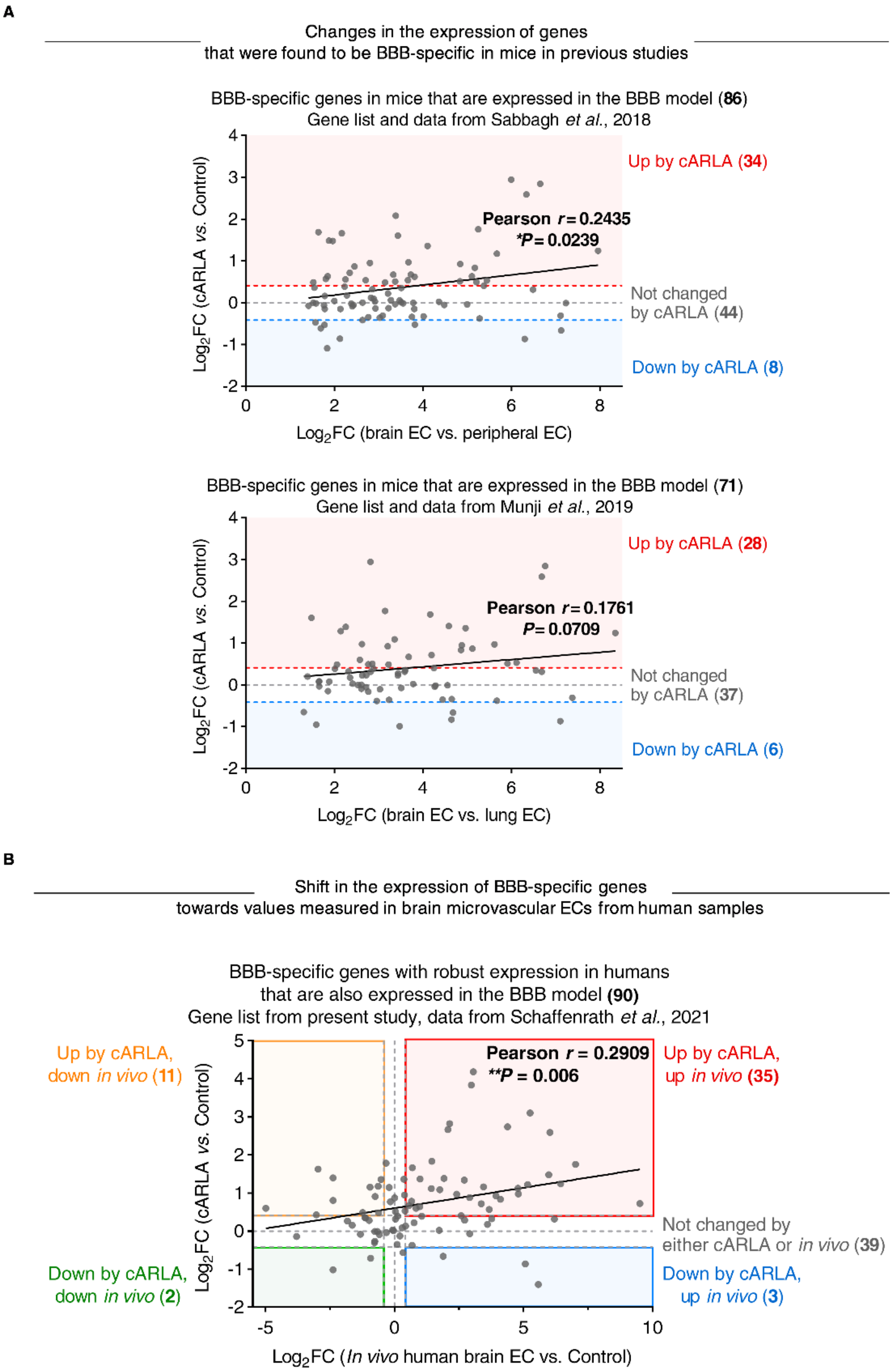


**Figure S12. Correlating the effect of cARLA with *in vivo* data from mice and humans. A)** Changes in gene expression induced by cARLA are plotted against changes between brain vs. peripheral ECs in mice *in vivo* using the gene list and data of Sabbagh *et al*.[53] and Munji *et al*.[4]. **B)** Changes in gene expression induced by cARLA are plotted against changes between human *in vivo* vs. control values using our gene list and data of Schaffenrath *et al.*[70]. Pearson correlation coefficients (two-tailed) and P-values are presented.

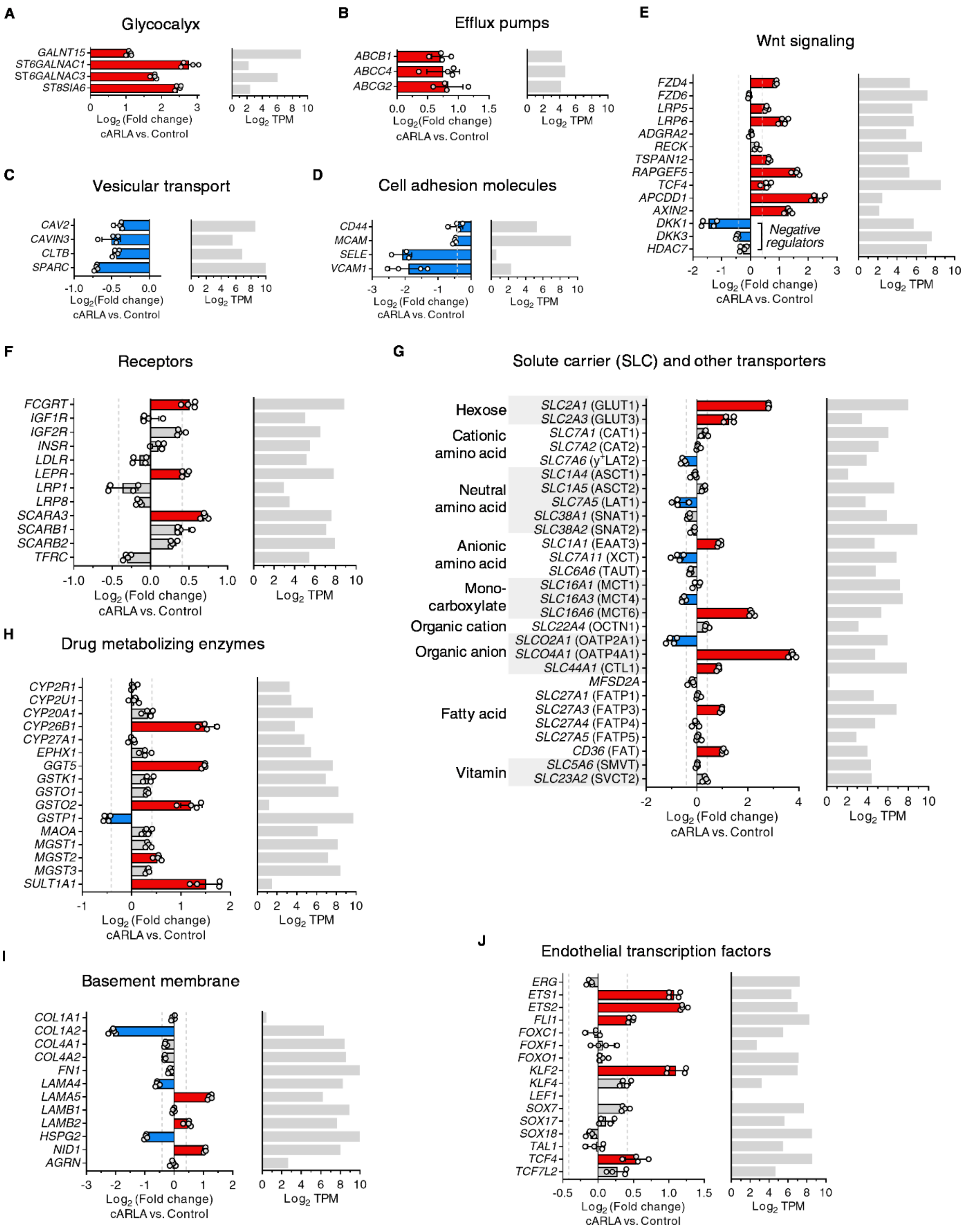


**Figure S13. Changes in gene expression related to different aspects of BBB function in brain-like ECs.** Gene expressional changes after 48 h cARLA treatment were determined using MACE-seq profiling and are presented in panels **A-J)** by category. Mean ± SD, *n*=4. Red and blue color indicates up- and downregulation, respectively, upon cARLA treatment. Grey color indicates no differential expression. TPM: mean transcript per million.

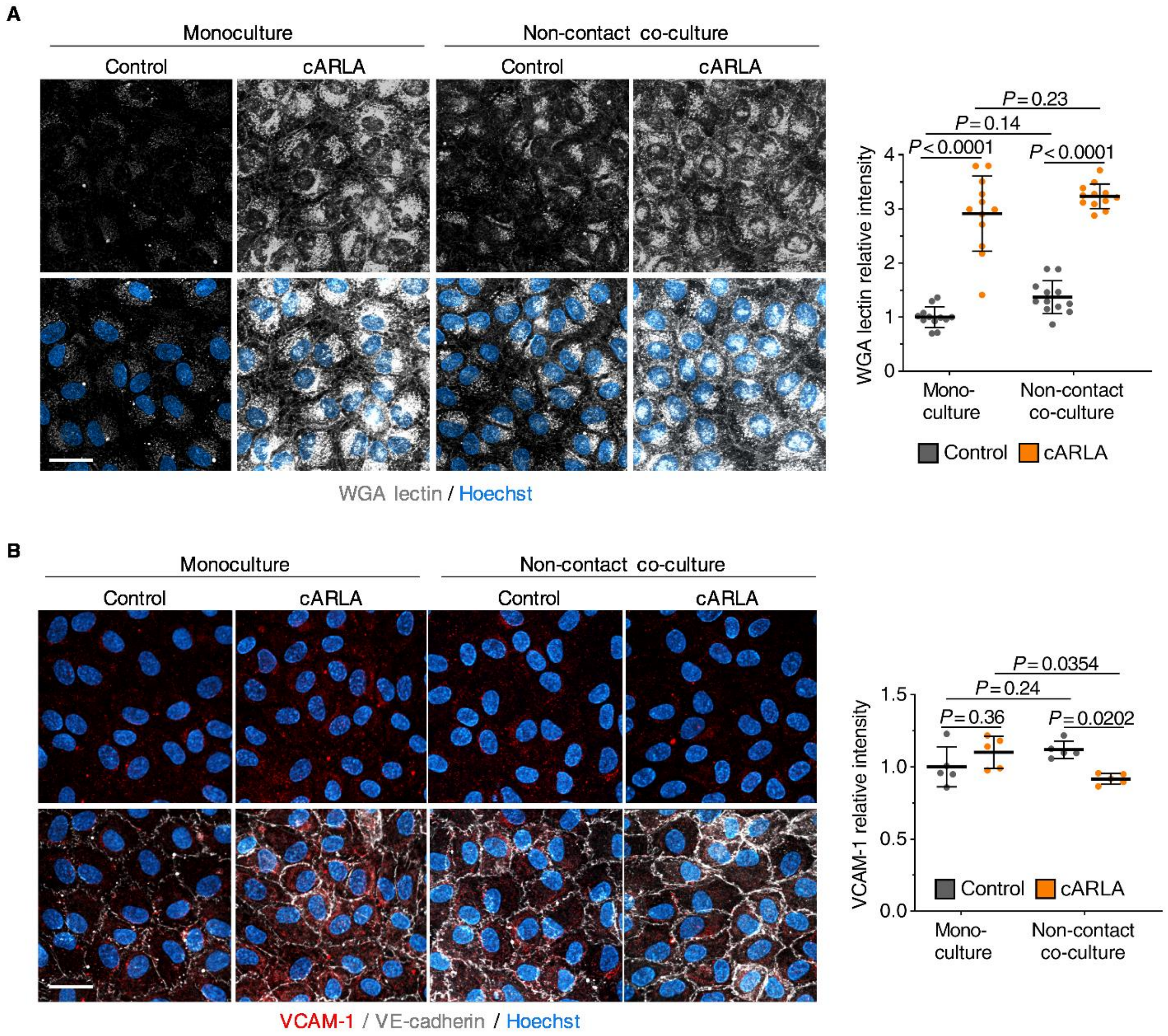


**Figure S14. The effect of cARLA and EC-PC co-culture on WGA lectin staining and VCAM-1 levels. A)** Wheat germ agglutining (WGA) lectin staining in human brain like ECs in monoculture or non-contact EC-PC co-culture. Bar: 50 µm. Quantification**:** Mean ± SD, ANOVA with Bonferroni's post-hoc test, *n*=12. **B)** VCAM-1 (red) and VE-cadherin (grey) immuno-staining in human brain like ECs in monoculture or co-culture. Bar: 50 µm. Quantification**:** Mean ± SD, ANOVA with Bonferroni's post-hoc test, *n*=5.

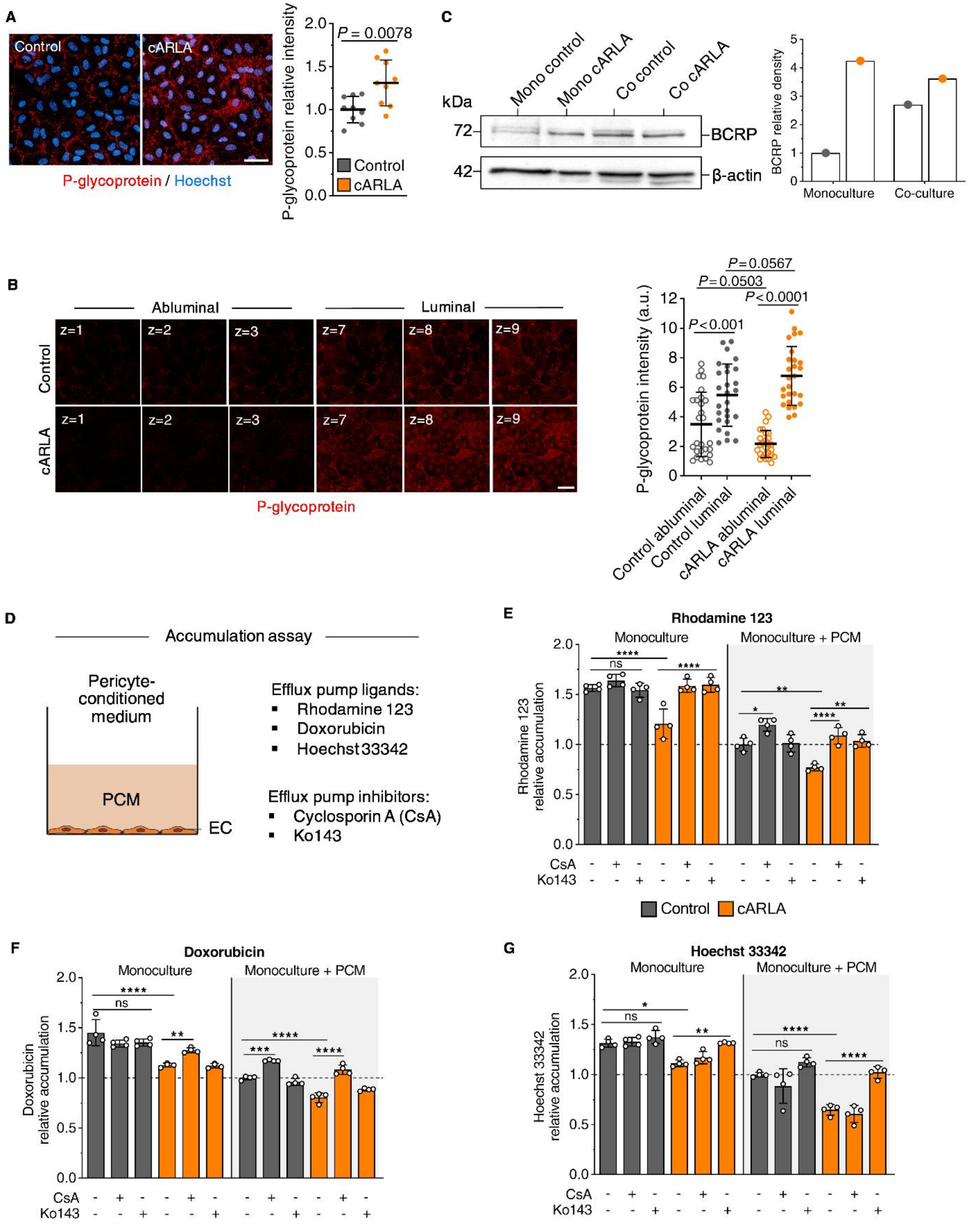


**Figure S15. Further validation of the effect of cARLA on efflux pump expression, localization and activity. A)** P-glycoprotein immunostaining in human brain-like ECs with an additional antibody. Quantification: Mean ± SD, unpaired t-test, *n*=9. Bar: 50 µm. **B)** Z-stacks from the same immunostaining demonstrating P-glycoprotein polarity in ECs. Representative abluminal (brain side) and luminal (blood side) slices (3 in each group) are shown. Quantification: Mean ± SD, ANOVA with Bonferroni's post-hoc test, *n*=27 slices from 9 z-stacks. Bar: 50 µm. **C)** Western blot of BCRP (*ABCG2*) in human brain-like ECs in monoculture and non-contact EC-PC co-culture using membrane-fractionated lysates. Quantification: Mean ± SD, ANOVA with Bonferroni's post-hoc test, *n*=1, normalized to β-actin band densities. **D)** Schematic drawing of the experimental setup, and the additional efflux pump ligands and inhibitors. **E)** Relative accumulation of rhodamine 123, **F)** doxorubicin, and **G)** Hoechst 33342 in ECs normalized to the PCM control group. Quantification: Mean ± SD, ANOVA with Bonferroni's post-hoc test, *n*=4. **P* < 0.05, ***P* < 0.01, ****P* < 0.001, *****P* < 0.0001.

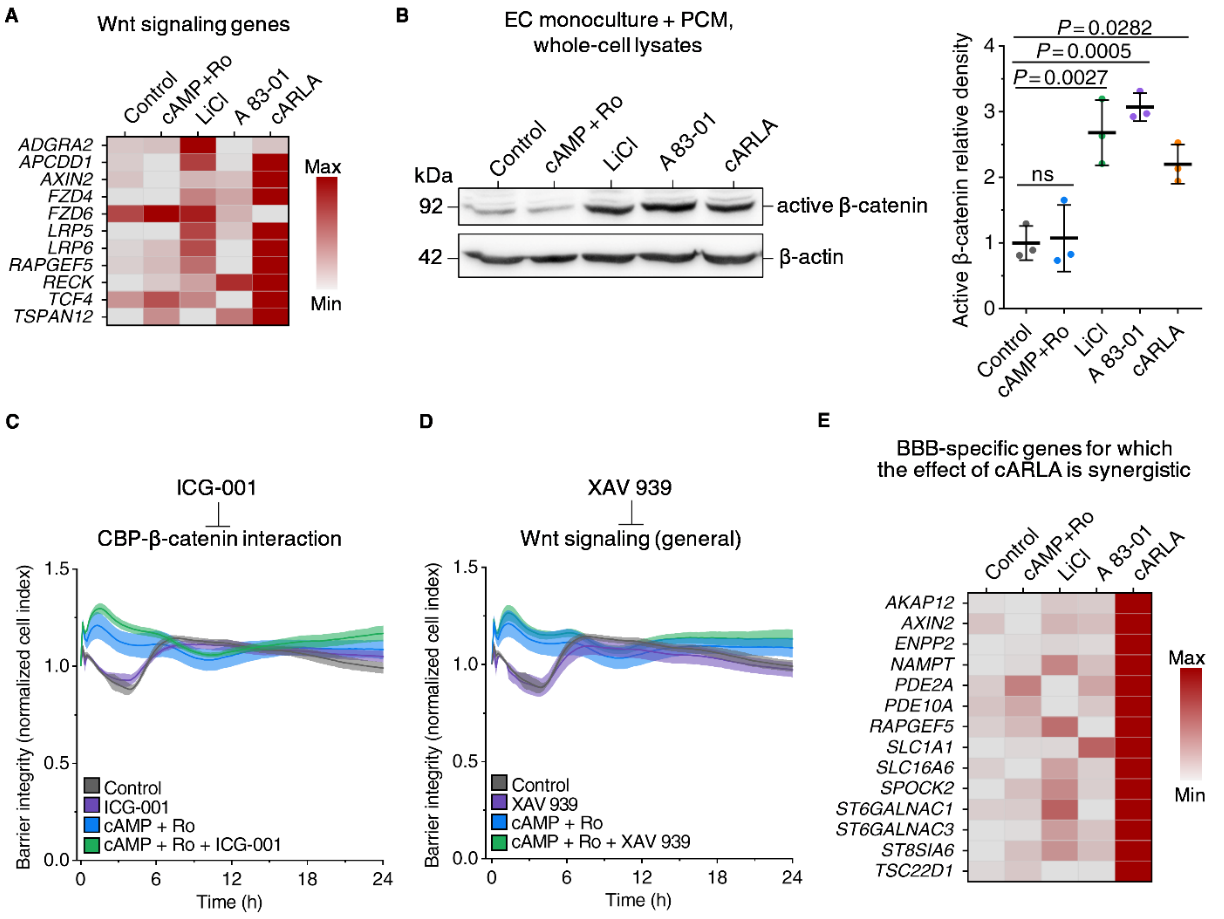


**Figure S16. The effect of cARLA and its constituents on Wnt signaling. A)** Scaled heat map of gene expression related to Wnt signaling. n=4 for control and cARLA samples, *n*=1 for cAMP+Ro, LiCl, and A 83-01 samples. **B)** Western blot of active β-catenin using whole-cell lysates. Quantification: Mean ± SD, ANOVA with Bonferroni's post-hoc test, *n*=3, normalized to β-actin band densities. **C)** Real-time impedance measurement of cAMP+Ro treatment, with or without ICG-001 and **D)** XAV 939. Higher normalized cell index values indicate higher barrier integrity. Mean ± SD, *n*=4. **E)** Scaled heat map of BBB-specific genes (gene list from present study) for which the effect of cARLA is synergistic.

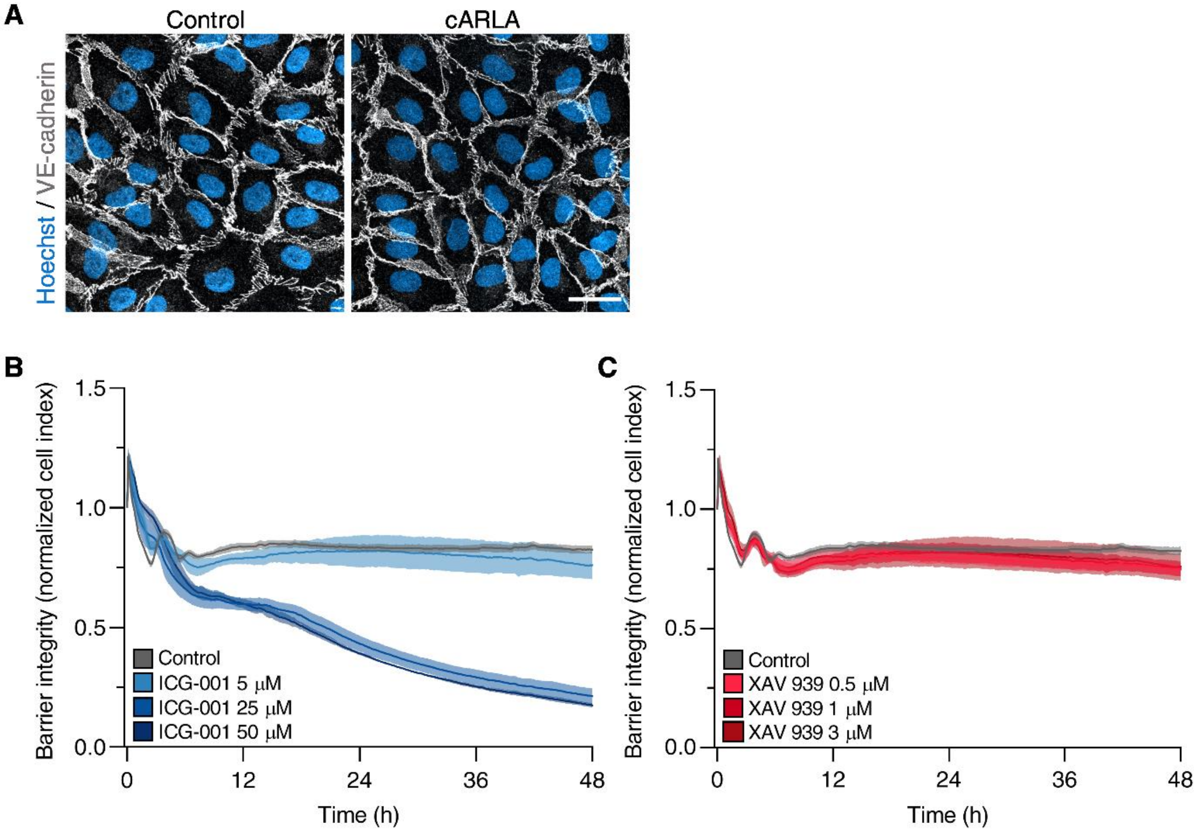


**Figure S17. Additional experiments on Wnt/β-catenin signaling in brain-like ECs. A)** Representative confocal microscopy images of VE-cadherin immunostaining (grey) show no disruption in VE-cadherin junctions upon 48 h cARLA treatment. Nuclei were stained with Hoechst (blue). Bar: 50 µm. **B)** Barrier integrity measured by impedance upon treatment with ICG-001, which specifically blocks the β-catenin-CBP interaction in the nucleus and **C)** XAV 939, which blocks the pathway upstream of nuclear translocation. Cells for this experiment did not receive cARLA treatment. Selected concentrations for further experiments were: ICG-001 5 µM, XAV 939 1 µM. Higher normalized cell index values and a higher area under the curve indicates increased barrier integrity. Mean ± SD, *n*=6.

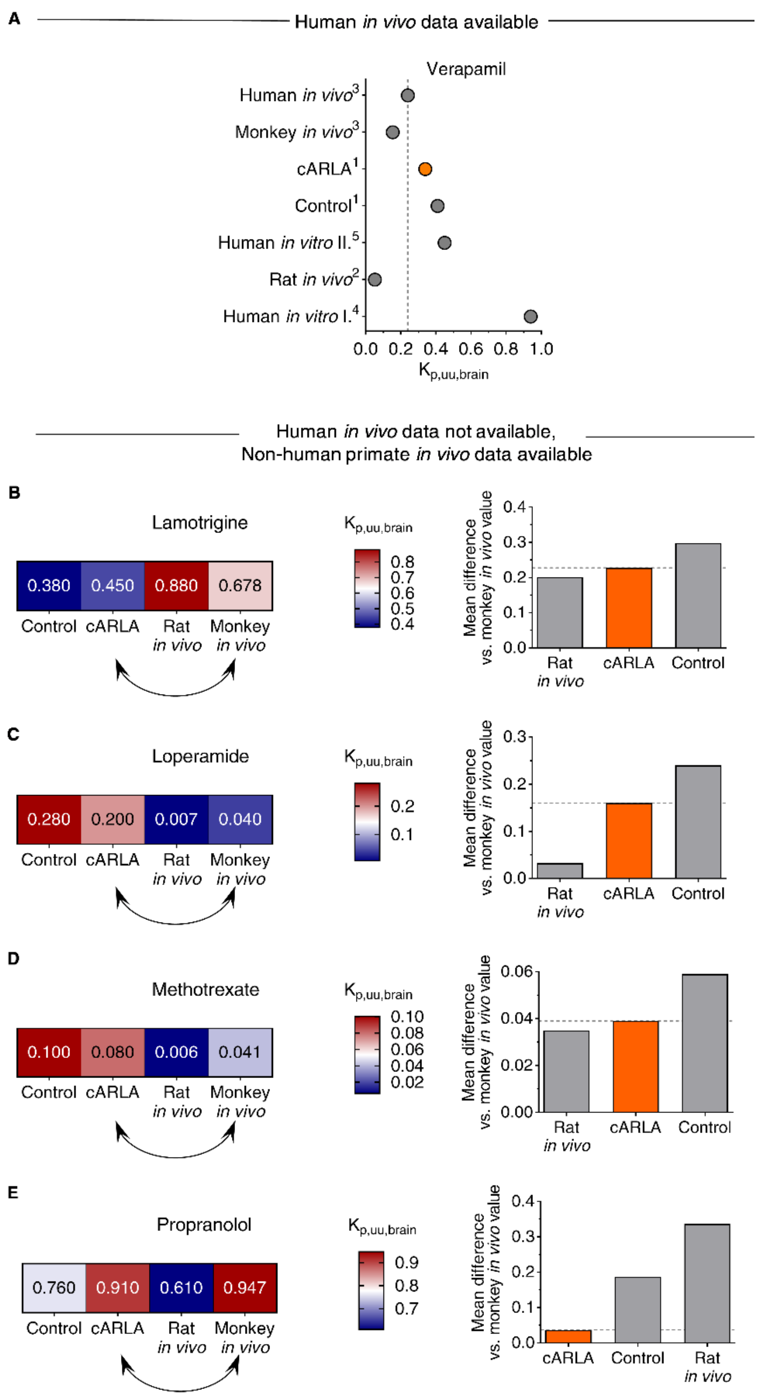


**Figure S18. Comparison of $K_{p,uu,brain}$ values for selected drugs across species and BBB models. A)** Comparison of $K_{p,uu,brain}$ measured *in vivo* in humans with its predictive models for the drug verapamil. Points are means, data obtained from articles above as well as from [4]Cecchelli *et al*., 2014 and [5]Moya *et al*., 2021. **B-E)** $K_{p,uu,brain}$ values for selected drugs which human *in vivo* data is not available. For these drugs, we used non-human primate data as a reference. Panels on the left: heat map of $K_{p,uu,brain}$ values. Panels on the right: Mean difference of $K_{p,uu,brain}$ values compared to non-human primate *in vivo* data. Lower values indicate better prediction.

**Table S1. The list of 100 BBB-specific genes with robust expression at the human brain endothelium.** $Log_2FC$: $log_2$(fold change), FDR: false discovery rate (adjusted P-value, Benjamini-Hochberg method), TPM: transcript per million.

| Gene symbol | Log2FC | FDR | TPM Control | | | | TPM cARLA | | | |
|---|---|---|---|---|---|---|---|---|---|---|
| *ABCB1* | 0,7145 | 5E-05 | 14,1767 | 13,3343 | 12,9603 | 15,3397 | 24,8375 | 28,4211 | 21,1541 | 25,2042 |
| *ABCC4* | 0,7549 | 1,14E-05 | 17,3592 | 16,7914 | 19,5503 | 21,9731 | 36,1856 | 38,9474 | 38,4115 | 24,9751 |
| *ABCG2* | 0,8355 | 3,49E-05 | 14,4660 | 9,6303 | 11,2030 | 14,0959 | 22,9104 | 23,5088 | 31,1745 | 19,7051 |
| *ADGRA2* | 0,0093 | 0,954093 | 99,2370 | 95,8096 | 104,7809 | 97,6352 | 89,7147 | 103,6842 | 103,2656 | 108,6072 |
| *ADIPOR2* | 1,3437 | 5,03E-56 | 58,4428 | 53,3373 | 60,6279 | 57,4203 | 150,9519 | 145,4386 | 147,8007 | 150,3086 |
| *APCDD1* | 2,3310 | 6,13E-09 | 2,0252 | 1,7285 | 1,7573 | 0,8292 | 8,5646 | 7,8947 | 11,4121 | 10,3108 |
| *BSG* | -0,0403 | 0,547823 | 415,7537 | 395,8318 | 377,8263 | 389,7115 | 396,5432 | 378,0703 | 379,6613 | 380,3542 |
| *CA4* | 0,4936 | 0,057109 | 0,5786 | 0,4939 | 0,6590 | 1,0365 | 3,2117 | 1,5789 | 2,5051 | 2,0622 |
| *CD109* | 0,3786 | 2,25E-06 | 86,7962 | 93,8341 | 99,9482 | 97,8425 | 125,2580 | 125,9650 | 131,6567 | 127,1666 |
| *CD320* | -0,1035 | 0,33261 | 93,7399 | 88,8955 | 82,5946 | 80,2225 | 84,3618 | 77,7193 | 74,3179 | 79,2786 |
| *CD34* | 0,5355 | 1,42E-07 | 61,0467 | 62,7207 | 79,0799 | 58,6640 | 89,5006 | 107,3685 | 96,0287 | 102,4207 |
| *CDC42EP3* | 1,0961 | 1,39E-17 | 26,3282 | 24,6932 | 31,6320 | 32,7524 | 65,0913 | 68,7720 | 59,2873 | 62,3231 |
| *CERT1* | 0,7539 | 3,92E-11 | 36,7437 | 37,2867 | 36,6843 | 35,4472 | 64,4490 | 61,5790 | 61,2357 | 68,2805 |
| *CGNL1* | 0,5013 | 2,94E-06 | 47,7379 | 42,2254 | 49,8643 | 43,7389 | 71,5148 | 67,5439 | 67,0809 | 66,2183 |
| *CLDN5* | 1,6487 | 3,9E-281 | 223,9342 | 241,7463 | 236,5808 | 225,3279 | 743,4115 | 738,7722 | 708,1073 | 752,0015 |
| *CTNNBIP1* | -0,2135 | 0,063614 | 58,7321 | 52,3496 | 55,7953 | 49,9577 | 44,5362 | 44,9123 | 43,9784 | 46,9715 |
| *DEGS2* | -0,0566 | - | 1,1573 | 0,2469 | 0,0000 | 0,0000 | 0,0000 | 0,1754 | 0,0000 | 0,2291 |
| *ENPP2* | 4,0400 | 2,31E-31 | 1,4466 | 2,2224 | 0,8787 | 1,4511 | 29,5480 | 29,1228 | 23,6592 | 27,2664 |
| *ESYT2* | 0,4231 | 7,64E-06 | 78,6952 | 78,2774 | 89,1846 | 69,0287 | 112,4110 | 100,0000 | 119,9663 | 108,8363 |
| *FAM13A* | 0,4319 | 0,035764 | 4,9185 | 7,4080 | 6,5900 | 7,0480 | 8,3505 | 12,4561 | 9,7420 | 14,2060 |
| *FGR* | 0,3184 | 0,128859 | 0,5786 | 0,4939 | 0,0000 | 0,6219 | 1,4988 | 1,5789 | 1,9484 | 1,1456 |
| *FLT1* | 1,0565 | 1,09E-29 | 65,6758 | 75,5612 | 70,2933 | 68,1995 | 142,8155 | 131,9299 | 167,2848 | 153,9747 |
| *FLVCR2* | 0,1987 | 0,310841 | 1,4466 | 1,4816 | 1,3180 | 0,8292 | 3,6400 | 1,7544 | 1,9484 | 2,2913 |
| *FN1* | -0,1530 | 0,000124 | 2210,9882 | 2354,2485 | 2416,5509 | 2282,5073 | 2061,5109 | 2113,8604 | 2183,8874 | 2018,3977 |
| *FOXC1* | -0,0553 | 0,739354 | 38,1903 | 50,3741 | 43,9333 | 48,7139 | 46,4632 | 38,2456 | 43,9784 | 43,9928 |
| *FOXF2* | - | - | 0,0000 | 0,0000 | 0,0000 | 0,0000 | 0,0000 | 0,0000 | 0,0000 | 0,0000 |
| *FOXL2* | 0,0156 | - | 0,0000 | 0,0000 | 0,0000 | 0,0000 | 0,0000 | 0,1754 | 0,2783 | 0,0000 |
| *FOXQ1* | -0,0276 | - | 0,0000 | 0,4939 | 0,2197 | 0,0000 | 0,0000 | 0,0000 | 0,0000 | 0,0000 |
| *FUT8* | 0,0828 | 0,593921 | 38,4796 | 37,7806 | 37,7826 | 42,9097 | 40,0397 | 43,6842 | 53,9987 | 37,1189 |
| *FZD4* | 0,8806 | 1,69E-35 | 98,0797 | 101,2421 | 99,7286 | 103,2321 | 186,2811 | 184,3860 | 191,2224 | 192,9266 |
| *FZD6* | -0,0570 | 0,712428 | 55,5496 | 49,6333 | 45,6906 | 44,9827 | 46,6773 | 47,3684 | 45,9268 | 44,9093 |
| *GCH1* | 0,2916 | 0,091759 | 10,4155 | 9,1365 | 8,5670 | 9,5355 | 16,0587 | 13,1579 | 12,5255 | 12,6021 |
| *GIMAP7* | 1,0873 | 1,01E-23 | 40,2156 | 58,0290 | 53,3790 | 50,3723 | 97,6370 | 121,9299 | 110,5026 | 111,1276 |
| *GLUL* | 0,3346 | 0,000205 | 70,5942 | 72,8449 | 77,9816 | 82,5028 | 93,5688 | 101,7544 | 102,1523 | 102,1916 |
| *GPD2* | -0,0566 | 0,784268 | 18,8058 | 17,5322 | 24,6026 | 23,4242 | 18,8422 | 21,4035 | 21,7108 | 17,1847 |
| *GPCPD1* | 0,4452 | 0,013783 | 12,4408 | 10,6181 | 11,8620 | 9,5355 | 19,4846 | 15,9649 | 20,5975 | 14,6643 |
| *GPR160* | 0,7980 | 0,008577 | 2,8932 | 1,7285 | 2,6360 | 1,4511 | 5,9953 | 5,0877 | 6,1236 | 4,8117 |
| *GSTA4* | 0,2012 | 0,188544 | 19,0952 | 18,0260 | 16,0357 | 17,8272 | 21,6257 | 23,8597 | 25,0510 | 19,7051 |
| *H1-2* | 0,3369 | 0,088587 | 4,0505 | 5,4325 | 3,9540 | 5,5969 | 7,4941 | 8,2456 | 6,4019 | 9,3943 |

| | | | | | | | | | | |
|---|---|---|---|---|---|---|---|---|---|---|
| *HSPA12B* | 0,7639 | 0,007794 | 2,8932 | 3,7040 | 2,1967 | 2,0729 | 6,8517 | 6,3158 | 5,2885 | 6,8739 |
| *HTRA3* | 1,1062 | 2,37E-11 | 15,6233 | 14,0751 | 15,8160 | 15,1324 | 33,4021 | 33,3333 | 34,2363 | 35,9733 |
| *IFNAR1* | 0,0841 | 0,515724 | 56,1282 | 58,7698 | 48,7660 | 49,7504 | 58,0255 | 58,2456 | 59,8440 | 57,9696 |
| *IL10RB* | 0,3377 | 0,108143 | 3,1825 | 2,4693 | 2,6360 | 3,3167 | 4,7106 | 4,9123 | 5,5669 | 6,1865 |
| *ITIH5* | -0,0196 | - | 0,2893 | 0,0000 | 0,2197 | 0,0000 | 0,0000 | 0,0000 | 0,0000 | 0,0000 |
| *ITM2A* | 0,1152 | 0,551239 | 10,4155 | 8,8895 | 6,8097 | 6,4261 | 7,4941 | 8,0702 | 12,5255 | 12,3730 |
| *KLF2* | 1,1050 | 1,34E-13 | 60,1787 | 86,4262 | 85,2306 | 89,9653 | 164,2271 | 162,8071 | 196,2326 | 198,6549 |
| *LDLRAD3* | 0,8969 | 1,7E-12 | 29,8000 | 32,5950 | 35,3663 | 30,2648 | 57,5973 | 68,7720 | 64,0191 | 56,8240 |
| *LEF1* | -0,0200 | - | 0,5786 | 0,2469 | 0,2197 | 0,2073 | 0,6423 | 0,0000 | 0,2783 | 0,0000 |
| *LRP8* | -0,1522 | 0,381158 | 14,7554 | 12,3466 | 11,6423 | 12,4376 | 10,9199 | 9,8246 | 10,5771 | 10,0817 |
| *MAOA* | 0,3129 | 0,002852 | 60,4680 | 57,2882 | 57,7723 | 69,2360 | 87,3594 | 72,6316 | 81,5548 | 78,8204 |
| *MARVELD2* | 0,2192 | 0,127953 | 19,3845 | 21,2361 | 24,3830 | 24,0460 | 26,9786 | 28,9474 | 30,0612 | 27,9537 |
| *MCF2L* | 0,0842 | - | 0,0000 | 0,2469 | 0,0000 | 0,0000 | 0,2141 | 0,3509 | 0,5567 | 0,2291 |
| *MFSD2A* | -0,2059 | 0,276884 | 2,3146 | 1,4816 | 1,5377 | 0,8292 | 0,2141 | 0,8772 | 0,5567 | 0,6874 |
| *MYO1E* | -0,4797 | 0,000513 | 39,9262 | 35,0643 | 40,4186 | 50,1650 | 28,4775 | 28,9474 | 29,7828 | 24,9751 |
| *NAMPT* | 1,3427 | 5,4E-115 | 149,2894 | 137,7880 | 137,5112 | 142,2033 | 360,9999 | 356,4914 | 374,0944 | 364,5443 |
| *NDRG1* | 1,1335 | 3,14E-36 | 58,1534 | 65,4369 | 54,4773 | 53,0671 | 126,7568 | 130,1755 | 131,6567 | 127,3957 |
| *NEBL* | 0,1980 | 0,309634 | 2,6039 | 2,2224 | 1,5377 | 3,1094 | 3,2117 | 4,0351 | 4,4535 | 3,8952 |
| *OCLN* | 0,0607 | 0,793976 | 2,3146 | 1,9755 | 2,4163 | 1,4511 | 2,5694 | 2,1053 | 2,7834 | 2,7495 |
| *PALM* | 0,6992 | 1E-08 | 35,5864 | 36,7928 | 35,5860 | 42,4951 | 70,0160 | 56,8421 | 68,1943 | 61,6357 |
| *PDE10A* | 1,3068 | 6,72E-10 | 10,1262 | 8,8895 | 8,1277 | 9,3282 | 25,0516 | 23,3333 | 20,3191 | 27,4955 |
| *PDE2A* | 1,6030 | 8,95E-65 | 41,9515 | 44,9416 | 41,5170 | 40,6295 | 129,5403 | 136,3158 | 126,6465 | 130,6036 |
| *PDE8A* | 0,3007 | 0,055866 | 16,7806 | 13,8282 | 13,6193 | 19,0710 | 20,7693 | 23,1579 | 21,1541 | 23,1420 |
| *PDXK* | -0,0146 | 0,930939 | 76,6700 | 80,0059 | 74,0276 | 67,7849 | 76,0113 | 72,4562 | 74,0395 | 73,5504 |
| *POLK* | 0,1615 | 0,271322 | 20,5418 | 24,9401 | 25,4813 | 22,3877 | 27,6210 | 27,0176 | 29,2261 | 29,3285 |
| *PROM1* | -0,0060 | - | 0,2893 | 0,2469 | 0,0000 | 0,2073 | 0,4282 | 0,0000 | 0,0000 | 0,2291 |
| *PTPRG* | 0,1557 | 0,315601 | 16,7806 | 19,0138 | 17,7930 | 22,3877 | 22,4822 | 23,1579 | 21,9892 | 24,9751 |
| *RAB11FIP5* | 0,2179 | 0,015915 | 81,5884 | 83,9568 | 78,2013 | 77,1131 | 97,8511 | 99,4737 | 93,8019 | 93,7138 |
| *RECK* | 0,2256 | 0,070509 | 38,7690 | 32,5950 | 36,2450 | 35,4472 | 41,1103 | 44,2105 | 48,9886 | 43,9928 |
| *RNASE1* | 0,2000 | 0,000709 | 275,7226 | 272,8597 | 273,0454 | 253,1052 | 305,7579 | 331,2282 | 316,4772 | 301,9921 |
| *RNF144B* | 0,6059 | 9,28E-15 | 90,8467 | 90,8709 | 89,8436 | 100,3300 | 139,8179 | 144,5615 | 150,3058 | 147,5591 |
| *SH3BP4* | -0,6789 | 2,88E-08 | 56,4175 | 57,7821 | 62,6049 | 74,6256 | 38,1127 | 40,0000 | 37,5764 | 37,3480 |
| *SHROOM1* | 1,3275 | 6,22E-30 | 36,4544 | 34,5705 | 36,4646 | 27,3627 | 82,4347 | 92,8071 | 78,2147 | 91,8807 |
| *SLC16A1* | -0,0153 | 0,90832 | 139,1632 | 141,4920 | 146,2979 | 157,3357 | 138,9614 | 152,8071 | 131,1000 | 157,4116 |
| *SLC16A2* | 0,0426 | 0,867181 | 3,7612 | 2,4693 | 3,7343 | 2,2802 | 3,4259 | 3,5088 | 2,7834 | 4,1243 |
| *SLC16A4* | 0,2517 | 0,146158 | 7,8117 | 9,8773 | 8,1277 | 10,7793 | 14,3458 | 11,4035 | 13,3605 | 13,0604 |
| *SLC1A1* | 0,8891 | 3,47E-08 | 19,0952 | 14,8159 | 18,0127 | 19,4856 | 32,9739 | 36,4912 | 36,7414 | 33,6820 |
| *SLC2A1* | 2,8217 | 1,19E-09 | 55,5496 | 62,4738 | 63,7033 | 66,7485 | 452,2134 | 445,9651 | 447,2989 | 443,3647 |
| *SLC38A5* | -0,4972 | 0,004465 | 24,5923 | 21,4831 | 21,5273 | 17,2053 | 15,6305 | 11,0526 | 11,9688 | 14,8934 |
| *SLC39A10* | 0,0056 | 0,979899 | 42,2408 | 35,8051 | 36,9040 | 36,4836 | 37,4703 | 37,0176 | 43,7000 | 35,7441 |
| *SLC40A1* | 0,6154 | 2,69E-10 | 64,5185 | 72,3510 | 63,0443 | 71,3089 | 95,0676 | 107,8948 | 116,6262 | 111,8150 |
| *SLC5A6* | 0,0033 | 0,991051 | 19,0952 | 21,4831 | 20,2093 | 18,4491 | 20,7693 | 21,2281 | 17,8140 | 20,1634 |
| *SLC6A6* | -0,2608 | 0,055178 | 33,5612 | 30,1257 | 30,9730 | 32,5451 | 25,6939 | 24,5614 | 25,3293 | 24,0585 |

| | | | | | | | | | | |
|---|---|---|---|---|---|---|---|---|---|---|
| *SLC7A1* | 0,3213 | 0,003351 | 50,6311 | 54,8189 | 59,0903 | 63,4318 | 76,4395 | 76,8421 | 83,2249 | 65,3018 |
| *SLC7A5* | -0,7048 | 0,000643 | 18,5165 | 17,7791 | 16,4750 | 19,2783 | 13,7034 | 7,5439 | 9,1854 | 9,1652 |
| *SLCO4A1* | 3,7475 | 1,5E-56 | 2,6039 | 4,4448 | 3,5147 | 3,3167 | 44,3221 | 48,9474 | 50,1019 | 55,4492 |
| *SPOCK2* | 0,5343 | 0,039923 | 2,8932 | 1,7285 | 2,1967 | 1,0365 | 3,6400 | 3,5088 | 6,6803 | 4,8117 |
| *ST3GAL6* | -0,9570 | 1,3E-08 | 31,5359 | 27,4094 | 33,8286 | 29,8502 | 14,9881 | 16,6667 | 12,8038 | 16,0390 |
| *ST6GALNAC1* | 2,7692 | 3,69E-09 | 1,1573 | 0,9877 | 0,6590 | 1,0365 | 7,2800 | 6,8421 | 8,9070 | 10,0817 |
| *ST6GALNAC3* | 1,7871 | 2,81E-51 | 23,4350 | 32,3481 | 27,2386 | 34,4107 | 106,2016 | 110,1755 | 106,0491 | 96,4633 |
| *ST8SIA6* | 2,4617 | 4,77E-09 | 0,8680 | 2,7163 | 1,3180 | 0,6219 | 9,6352 | 9,8246 | 8,6287 | 8,9360 |
| *TBC1D4* | 0,0438 | 0,779575 | 56,4175 | 69,8817 | 61,9459 | 70,4797 | 67,2325 | 68,4211 | 77,3796 | 61,8648 |
| *TBX3* | 2,5939 | 1,09E-47 | 6,3651 | 8,8895 | 8,1277 | 9,9501 | 48,3903 | 55,9649 | 53,9987 | 53,3871 |
| *TCF4* | 0,5404 | 7,01E-08 | 273,6973 | 296,3182 | 305,3364 | 330,4256 | 391,6185 | 454,2107 | 519,9467 | 462,3824 |
| *TFRC* | -0,3073 | 0,009355 | 50,6311 | 51,3618 | 53,5986 | 43,5316 | 37,8986 | 40,1755 | 36,7414 | 38,7228 |
| *TNFRSF1B* | 1,0935 | 6,11E-44 | 75,8020 | 87,1670 | 77,3226 | 77,1131 | 164,8695 | 171,0527 | 171,1816 | 182,1576 |
| *TSC22D1* | 1,3604 | 2,28E-58 | 61,0467 | 63,2146 | 61,2869 | 61,5661 | 152,8790 | 158,9474 | 169,7899 | 165,2020 |
| *TSC22D3* | 1,7663 | 1,6E-172 | 127,8797 | 123,9598 | 117,9609 | 108,8290 | 413,2443 | 392,4563 | 428,6498 | 412,8905 |
| *TSPAN5* | 0,4351 | 5,21E-07 | 73,4874 | 79,5121 | 75,3456 | 85,8195 | 113,2675 | 109,4737 | 108,8325 | 109,7528 |
| *ZIC3* | - | - | 0,0000 | 0,0000 | 0,0000 | 0,0000 | 0,0000 | 0,0000 | 0,0000 | 0,0000 |
| *ZNF366* | 0,7166 | 0,017506 | 0,8680 | 1,7285 | 1,3180 | 1,8656 | 3,2117 | 5,2632 | 4,4535 | 3,2078 |

**Table S2. The list of BBB-specific genes in mice as reported by previous studies.** For correlations, we selected 86 genes from Sabbagh *et al.,* 2018, ref. 53 and 71 genes from Munji *et al.*, 2019, ref. 4, which are expressed at the human brain endothelium and also in the human BBB model. $Log_2FC$: $log_2$(fold change).

| Sabbagh et al., 2018 | | Munji et al., 2019 | |
|---|---|---|---|
| Gene symbol | $Log_2FC$ brain vs. peripheral EC | Gene symbol | $Log_2FC$ brain vs. lung EC |
| *Abcb1a* | 5,171 | *Abcb1a* | 4,859 |
| *Abcc4* | 2,455 | *Abcc4* | 5,119 |
| *Abhd2* | 3,368 | *Abcg2* | 2,618 |
| *Acox1* | 1,580 | *Abhd2* | 2,852 |
| *Alas1* | 2,134 | *Ablim1* | 2,627 |
| *Apcdd1* | 6,335 | *Alas1* | 1,850 |
| *Arl4a* | 2,702 | *Apcdd1* | 6,687 |
| *Axin2* | 1,634 | *Axin2* | 4,164 |
| *Bsg* | 4,997 | *Bsg* | 4,265 |
| *Car4* | 5,251 | *Cobll1* | 2,709 |
| *Cd320* | 3,219 | *Esyt2* | 2,056 |
| *Cdkn2b* | 3,659 | *Extl3* | 2,608 |
| *Cgnl1* | 1,805 | *Far2* | 4,648 |
| *Cldn5* | 2,162 | *Flt1* | 3,359 |
| *Cthrc1* | 2,634 | *Fosb* | 2,142 |
| *Ctsh* | 3,471 | *Foxc1* | 3,506 |
| *Def6* | 3,781 | *Fry* | 2,766 |
| *Efr3b* | 2,928 | *Fyco1* | 3,588 |
| *Esyt2* | 1,520 | *Glul* | 2,880 |
| *Extl3* | 2,634 | *Gpcpd1* | 3,676 |
| *Faim* | 2,380 | *Gpd2* | 2,607 |
| *Far2* | 3,750 | *Gsta4* | 6,552 |
| *Flvcr2* | 4,841 | *Igf1r* | 4,552 |
| *Fn1* | 3,800 | *Isyna1* | 3,231 |
| *Fzd6* | 2,403 | *Itga6* | 1,880 |
| *Gbe1* | 1,773 | *Itm2a* | 6,682 |
| *Golim4* | 1,778 | *Kif26a* | 3,140 |
| *Gpcpd1* | 3,331 | *Ldlrad3* | 4,878 |
| *Gpr4* | 1,834 | *Lrp8* | 7,372 |
| *Htra3* | 5,668 | *Lsr* | 6,113 |
| *Isyna1* | 3,025 | *Ly75* | 2,765 |
| *Itm2a* | 6,484 | *Maoa* | 2,005 |
| *Kcnj2* | 3,811 | *Max* | 2,533 |
| *Ldlrad3* | 2,800 | *Mfsd7c* | 3,202 |
| *Lipa* | 1,875 | *Mkl2* | 1,388 |
| *Lrp8* | 7,108 | *Nebl* | 4,192 |
| *Lsr* | 5,441 | *Nes* | 1,590 |
| *Ly75* | 5,207 | *Net1* | 1,650 |
| *Maoa* | 2,231 | *Nr4a1* | 3,030 |
| *Max* | 1,525 | *Ocln* | 2,317 |
| *Mospd1* | 1,759 | *Palmd* | 1,307 |
| *Mpzl1* | 1,995 | *Parvb* | 1,651 |

| | | | |
|---|---|---|---|
| *Nampt* | 4,101 | *Pdxk* | 4,284 |
| *Nebl* | 2,344 | *Pltp* | 1,477 |
| *Nomo1* | 3,371 | *Polk* | 3,438 |
| *Nos3* | 1,843 | *Rab11fip1* | 3,600 |
| *Ocln* | 2,855 | *Reck* | 2,819 |
| *Pcdh19* | 3,086 | *Sgms1* | 3,429 |
| *Pdxk* | 3,552 | *Sgms2* | 2,957 |
| *Plpp6* | 2,250 | *Shroom1* | 4,961 |
| *Pltp* | 3,433 | *Slc16a2* | 2,352 |
| *Pmaip1* | 3,817 | *Slc16a4* | 5,909 |
| *Ppard* | 2,882 | *Slc1a1* | 5,618 |
| *Proser2* | 4,011 | *Slc2a1* | 6,765 |
| *Rab11fip1* | 3,509 | *Slc30a1* | 2,411 |
| *Rnpep* | 1,412 | *Slc38a5* | 4,679 |
| *Rras2* | 1,687 | *Slc39a10* | 3,779 |
| *Slc16a1* | 7,228 | *Slc6a6* | 4,443 |
| *Slc16a2* | 2,279 | *Slc7a1* | 4,247 |
| *Slc16a4* | 4,836 | *Slc7a5* | 7,103 |
| *Slc1a1* | 3,673 | *Slco2b1* | 2,812 |
| *Slc2a1* | 6,649 | *Sorbs2* | 3,471 |
| *Slc30a1* | 2,915 | *Sparcl1* | 4,586 |
| *Slc31a1* | 2,826 | *Spock2* | 8,349 |
| *Slc38a5* | 7,121 | *Tfrc* | 5,672 |
| *Slc39a10* | 4,358 | *Tmc7* | 4,643 |
| *Slc3a2* | 4,484 | *Tob1* | 2,570 |
| *Slc40a1* | 3,727 | *Tsc22d1* | 2,252 |
| *Slc46a3* | 3,385 | *Tspan5* | 3,222 |
| *Slc5a6* | 3,506 | *Usp33* | 2,714 |
| *Slc7a1* | 5,377 | *Utrn* | 1,656 |
| *Slc7a5* | 6,302 | | |
| *Slco2b1* | 5,993 | | |
| *Smox* | 3,208 | | |
| *Spock2* | 7,960 | | |
| *St3gal6* | 2,120 | | |
| *Stard13* | 2,426 | | |
| *Swap70* | 1,572 | | |
| *Tcf7* | 2,766 | | |
| *Tdrp* | 3,274 | | |
| *Tfrc* | 5,279 | | |
| *Tmem123* | 1,547 | | |
| *Tspan5* | 3,144 | | |
| *Ttyh2* | 5,107 | | |
| *Wwtr1* | 1,779 | | |
| *Zfp366* | 1,959 | | |

**Table S3. Properties of small molecule drugs tested in this study.** MW: molecular weight, CNS indicates if a compound readily enters (+) or does not enter (-) the central nervous system *via* the systemic circulation.

| Name | MW | Manufacturer, catalog number | CNS | BBB transport | Recovery (%) |
|---|---|---|---|---|---|
| Atenolol | 266.3 | Sigma-Aldrich, cat# 330892 | - | Passive hydrophilic, (efflux) | 88.16 ± 2.87 |
| Indomethacin | 357.8 | Sigma-Aldrich, cat# I7378 | + | Passive lipophilic | 102.08 ± 2.40 |
| Lamotrigine | 256.1 | Sigma-Aldrich, cat# PHR1392 | + | Passive lipophilic | 88.70 ± 0.82 |
| Loperamide | 513.5 | Sigma-Aldrich, cat# L4762 | - | Passive hydrophilic, efflux | 90.22 ± 9.38 |
| Methotrexate | 454.4 | Sigma-Aldrich, cat# M9929 | - | Efflux | 95.46 ± 1.63 |
| Propranolol | 295.8 | Sigma-Aldrich, cat# P0884 | + | Passive lipophilic | 87.47 ± 6.36 |
| Salicylic acid | 138.1 | Sigma-Aldrich, cat# 27301 | - | Efflux, influx | 94.25 ± 3.33 |
| Tacrine | 234.7 | Sigma-Aldrich, cat# A3773 | + | Influx | 92.41 ± 1.57 |
| Verapamil | 491.1 | Sigma-Aldrich, cat# V106 | + | Passive lipophilic, efflux | 100.14 ± 2.91 |
| Vinblastine | 909.1 | Sigma-Aldrich, cat# V1377 | - | Efflux | 82.49 ± 1.93 |

**Table S4. Parameters for measuring the passage of small molecule drugs across the BBB co-culture model.** For all compounds, a Kinetex C8 2.6 µm, 100 A, 100×4.6 mm analytical column was used. The eluents were A: $H_2O$ + 0.1% formic acid and B: ACN + 0.1% formic acid. DP:declustering potential, EP: entrance potential, CE: collision energy, CXP: collision cell exit potential, CG: curtain gas, IS: ionspray voltage, T: temperature.

| Name | Instru-ment | Scan mode | Pola-rity | Transi-tion | DP (V) | EP (V) | CE (V) | CXP (V) | CG | CAD | Gas 1 | Gas 2 | IS (V) | T (°C) |
|---|---|---|---|---|---|---|---|---|---|---|---|---|---|---|
| Atenolol | Sciex API 4000 | MRM | + | 267/145 | 51 | 10 | 35 | 10 | 40 | 45 | 45 | 4 | 5500 | 450 |
| Indomethacin | Sciex API 4000 | MRM | + | 358/139 | 81 | 10 | 21 | 16 | 40 | 45 | 45 | 4 | 5500 | 450 |
| Lamotrigine | Sciex API 4000 | MRM | + | 256/211 | 96 | 10 | 37 | 14 | 40 | 45 | 45 | 4 | 5500 | 450 |
| Loperamide | Sciex API 4000 | MRM | + | 477/266 | 101 | 10 | 33 | 18 | 40 | 45 | 45 | 4 | 5500 | 450 |
| Methotrexate | Sciex x500r QTOF | TOF-MS | + | 494/455 | 100 | - | 30 | - | 45 | 40 | 45 | 7 | 5500 | 500 |
| Propranolol | Sciex API 4000 | MRM | + | 260/116 | 51 | 10 | 25 | 22 | 40 | 45 | 45 | 4 | 5500 | 450 |
| Salicylic acid | Sciex API 4000 | MRM | - | 137/93 | -80 | -10 | -22 | -15 | 40 | 45 | 45 | 4 | -4500 | 450 |
| Tacrine | Sciex API 4000 | MRM | + | 199/171 | 56 | 10 | 43 | 12 | 40 | 45 | 45 | 4 | 5500 | 450 |
| Verapamil | Sciex API 4000 | MRM | + | 455/165 | 116 | 10 | 35 | 12 | 40 | 45 | 45 | 4 | 5500 | 450 |
| Vinblastine | Sciex API 4000 | MRM | + | 811/355 | 196 | 10 | 47 | 10 | 40 | 45 | 45 | 4 | 5500 | 450 |
